# Elevated nuclear TDP-43 induces constitutive exon skipping

**DOI:** 10.1101/2023.05.11.540291

**Authors:** Rogger P. Carmen-Orozco, William Tsao, Yingzhi Ye, Irika R. Sinha, Koping Chang, Vickie Trinh, William Chung, Kyra Bowden, Juan C. Troncoso, Seth Blackshaw, Lindsey R. Hayes, Shuying Sun, Philip C. Wong, Jonathan P. Ling

## Abstract

Cytoplasmic inclusions and loss of nuclear TDP-43 are key pathological features found in several neurodegenerative disorders, suggesting both gain- and loss-of-function mechanisms of disease. To study gain-of-function, TDP-43 overexpression has been used to generate *in vitro* and *in vivo* model systems. Our study shows that excessive levels of nuclear TDP-43 protein lead to constitutive exon skipping that is largely species-specific. Furthermore, while aberrant exon skipping is detected in some human brains, it is not correlated with disease, unlike the incorporation of cryptic exons that occurs after loss of TDP-43. Our findings emphasize the need for caution in interpreting TDP-43 overexpression data, and stress the importance of controlling for exon skipping when generating models of TDP-43 proteinopathy. Understanding the subtle aspects of TDP-43 toxicity within different subcellular locations is essential for the development of therapies targeting neurodegenerative disease.

## Introduction

Transactivation response element DNA-binding protein 43 (*TARDBP*, TDP-43) is an RNA-binding protein implicated in a variety of age-related disorders such as ALS-FTD (*1–3*), Alzheimer’s Disease (*4*), LATE (*5*), Paget’s Disease of Bone (*6*), and IBM (*7*). TDP-43 proteinopathy is characterized by large cytoplasmic inclusions and loss of nuclear TDP-43 staining, suggesting that both gain- and loss-of-function effects could contribute to disease pathogenesis.

Genetic deletion models have demonstrated that TDP-43 is an essential gene (*8–13*). Loss of TDP-43 function has been linked to neurodegeneration via disruptions in the splicing repression of nonconserved cryptic exons (*7, 14–28*). Indeed, compelling evidence from recent studies suggests that cryptic exons found in the genes *STMN2* (*16–18*) and *UNC13A* (*21, 22*) may contribute significantly to disease pathogenesis. Repression of cryptic exons in these and other genes represents a promising therapeutic strategy for ALS-FTD and other neurodegenerative disorders.

By contrast, transgenic models of TDP-43 overexpression also exhibit toxicity, but do not reproduce the cytoplasmic aggregates observed in human disease (*29–41*). Despite the absence of cytoplasmic aggregation, these TDP-43 transgenic models exhibit a dose-dependent toxicity that may be linked to excessive levels of nuclear TDP-43. Indeed, TDP-43 autoregulates the stability of its own mRNA by binding to an ultraconserved element in its 3’UTR (*42–44*). Only a single genomic copy of *Tardbp* is required for survival, as *Tardbp* heterozygotes exhibit the same levels of TDP-43 protein as wildtypes (*10*). Collectively, these data indicate that elevated levels of TDP-43 protein are harmful, even in non-disease contexts.

To circumvent the toxic effects associated with TDP-43 transgenic mice, alternatives have been developed that involve the use of adeno-associated virus (AAV)-mediated delivery of TDP-43 (*45*) and the generation of transgenic TDP-43 models that feature mutated nuclear localization signals, such as rNLS8 (*46–49*). Findings from these studies suggest a potential common mechanism of toxicity involving microglial activation (*45–47*). However, the molecular mechanisms that underlie toxicity due to TDP-43 accumulation remain poorly understood. Fully characterizing these mechanisms will be crucial to accurately interpret TDP-43 overexpression models and their relevance to human disease.

In this work, we demonstrate that excessive levels of nuclear TDP-43 protein result in the splicing repression of coding exons that are normally incorporated into mRNAs. Most of these exons are constitutively spliced under normal cellular conditions. With sufficiently high levels of overexpression, TDP-43 autoregulation can be overwhelmed and excessive nuclear TDP-43 can lead to aberrant constitutive exon repression. TDP-43 preferentially binds to long repetitive UG repeats (*15, 50*), but with higher concentrations in the nucleus, we theorize that TDP-43 can bind to shorter, less optimal UG-containing motifs present at these constitutive exons.

Interestingly, the constitutive exons repressed by excessive TDP-43 are divergent between mouse and human neurons. We also find that, while aberrant constitutive exon skipping can be detected in some human brain samples, constitutive exon skipping does not correlate with disease. By contrast, TDP-43 dependent cryptic exons are found only in human disease tissues or biofluids. Our findings suggest that constitutive exon skipping plays a role in the toxicity observed with TDP-43 overexpression and this aberrant exon skipping should be carefully mitigated when generating models of TDP-43 proteinopathy.

## Results

Transgenic models of TDP-43 overexpression exhibit a dose-dependent toxicity, but do not recapitulate the nuclear clearance and cytoplasmic aggregation that are hallmarks of TDP-43 proteinopathy. TDP-43 overexpression models also do not exhibit any disruptions in cryptic exon repression, as found in models of TDP-43 loss-of-function (*14*). These findings suggest that increasing TDP-43 protein to a degree that significantly exceeds physiological levels introduces gain-of-function toxicity that may not be applicable to the pathogenesis of human disease. Therefore, to determine whether slight elevations in TDP-43 levels could produce better models of disease, we generated transgenic mice that express wildtype TDP-43 (TDP-43^WT^) or TDP-43 carrying the G298S mutation (TDP-43^G298S^) that is associated with familial ALS (*51*), under the control of the Thy1.2 promoter (Fig. 1A). Immunoblotting analysis indicates that total TDP-43 protein levels in these transgenic models was approximately 50% higher than in non-transgenic controls (Fig. 1B). Indeed, transgenic animals developed a progressive motor deficit that began with hind-limb clench and hemiparesis, eventually leading to end-stage paralysis. Behavioral testing showed marked reduction on hanging time in transgenic lines compared with littermate controls (Fig. 1C) and animal weights for both transgenic mouse lines markedly decreased over time (Supplementary Fig. 1). The Kaplan-Meier survival curve indicated that lifespans of individuals from both transgenic lines were significantly shorter than their littermate controls (Fig. 1D). Pathological examination showed that TDP-43 overexpression was confined to the nucleus of neurons from the cortex and spinal cord, without any cytoplasmic inclusions in either transgenic line. Furthermore, histological analysis revealed reduction in axonal diameter, neuromuscular denervation and muscle degeneration in both transgenic lines, reflecting the severity of their condition (Supplementary Fig. 1). In summary, both TDP-43^WT^ and TDP-43^G298S^ mouse lines exhibited similar phenotypes of mild motor deficits that appeared to be mutation-independent.

**Figure 1.**
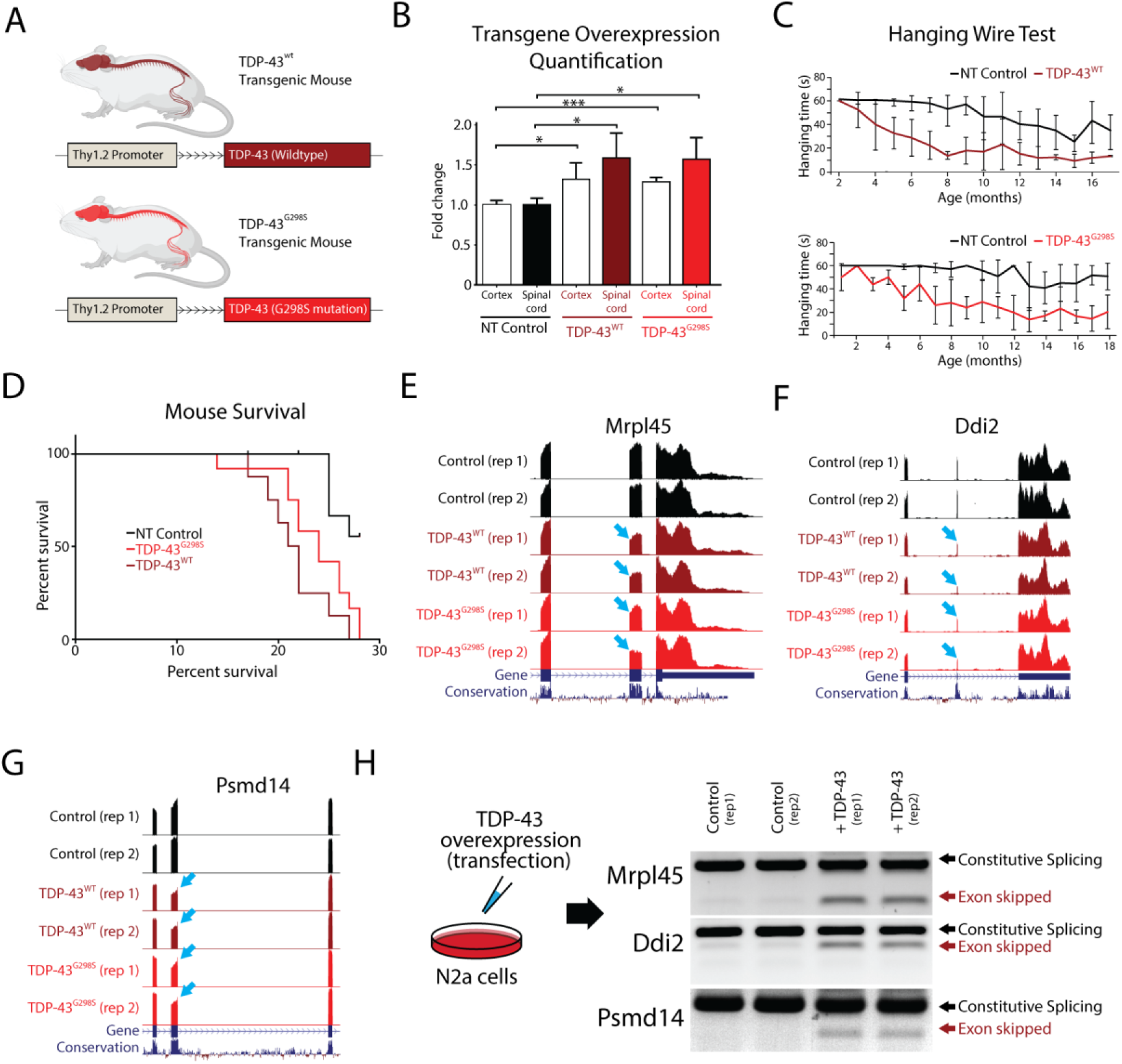
TDP-43 overexpression in mice leads to skipping of constitutive exons. Overexpression of TDP-43 has been shown to be toxic when under the control of different promoters, leading to premature death in animals. (**A**) In this study, we investigated the effects of overexpressing both human TDP-43 (TDP-43^WT^) and TDP-43 carrying a G298S mutation (TDP-43^G298S^), under the control of the weak Thy1.2 promoter in mice. (**B**) We compared TDP-43 levels in the spinal cord and cortex of transgenic mice to controls using immunoblotting. We observed approximately 1.5x and 1.3x overexpression of both transgenes in the spinal cord and cortex, respectively (* p<0,05, ***p<0.001). (**C**) We measured the hanging time of transgenic mice compared to their littermate controls and found a reduction associated with age, indicative of a motor neuron deficit. (**D**) Both transgenic lines had shorter survival times compared to nontransgenic (NT) controls (NT vs WT: p=0.0005, NT vs G298S: p=0.0035), but there were no differences between the two transgenic lines (WT vs G298S: p=0.1260). (**E**-**G**) To investigate the underlying mechanism of TDP-43 toxicity, we performed RNA-Seq analysis on isolated mouse spinal cords and found several instances of exon skipping (arrows) in both transgenic lines. (**H**) To confirm that these skipping events were directly caused by TDP-43 overexpression and not an indirect effect of disease progression, we transfected mouse N2a cells with human TDP-43 and evaluated different targets using double band RT-PCR. We observed exon skipping events only in transfected cells.

We next hypothesized that the observed phenotypes could be explained by changes in gene expression or alternative splicing due to the relatively low levels of TDP-43 overexpression. To profile these transcriptomic changes, we surgically isolated the ventral horn of the spinal cord and conducted bulk RNA sequencing (RNA-Seq) on the TDP-43^WT^ and TDP-43^G298S^ lines, as well as non-transgenic controls. TDP-43 has been well-described as a splicing repressor and loss of TDP-43 function leads to the inclusion of nonconserved cryptic exons. Interestingly, analysis of transgenic RNA-Seq data revealed that overexpression of both TDP-43^WT^ and TDP-43^G298S^ leads to the skipping of constitutive conserved exons (Fig. 1E-G). To verify that these splicing events were indeed due to TDP-43 overexpression and not due to the secondary effects of neuronal degeneration, we overexpressed TDP-43 in mouse N2a cells and performed a double band RT-PCR to detect the same targets identified from TDP-43^WT^ and TDP-43^G298S^ transgenic RNA-Seq data. As predicted, we observed prominent constitutive exon skipping only following TDP-43 overexpression (Fig. 1H). Together, these data indicate that even mild overexpression of TDP-43 (50% increase over controls) is sufficient to induce aberrant exon skipping and that exon skipping could be mediating the dose-dependent toxicity of TDP-43 overexpression.

Given the evolutionary conservation of the coding exons skipped by TDP-43 overexpression in mice, we hypothesized that syntenic exons in the human genome might also be skipped by TDP-43 overexpression in human cells. To profile exon skipping, we transduced human i3Neurons with lentivirus expressing TDP-43 and examined the splicing patterns of the equivalent exons observed to be skipped in mice (Fig. 2A). Unexpectedly, a large majority of skipped exons in mice were not skipped in human cells following TDP-43 overexpression, despite a ∼1.6x fold increase in TDP-43 protein (Supplementary Fig. 2, Fig. 3A). In order to understand these differences, we expanded our RNA-Seq analysis on transduced i3Neurons to include all splicing events and found that exon skipping still occurred across multiple genes (78 skipped exons, Supplementary Fig. 3) but at sites entirely different from those found in mice (Supplementary Fig. 3). The genes affected by exon skipping regulate a variety of cellular pathways including those associated with intellectual disability, synaptic activity, and mitochondrial proteins (Fig. 2B-C). Three of the clearest splicing repression events (∼90% reduction) include exons in *HYOU1*, *NUP93*, and *XPNPEP1*, where the corresponding exon in mice remains constitutively spliced (Fig. 2D-F). These RNA-Seq results were validated by double and single band RT-PCR analysis (Fig. 2G-H). UG repeats serve as the consensus binding site for TDP-43. An analysis of exons repressed by TDP-43 overexpression reveals the presence of short UG motifs that may be responsible for species-specific exon skipping, but these UG motifs are not as long as UG repeats associated with cryptic exons (Fig. 2I) (*14, 52*). This suggests that smaller UG repeats may be available for TDP-43 to bind when protein concentrations of nuclear TDP-43 exceed a certain threshold. Overall, our data indicate that while TDP-43 overexpression leads to aberrant exon skipping across the transcriptome, the exons that are skipped appear to be species-specific and generally exhibit shorter UG repeats than cryptic exons (Fig. 3A).

**Figure 2.**
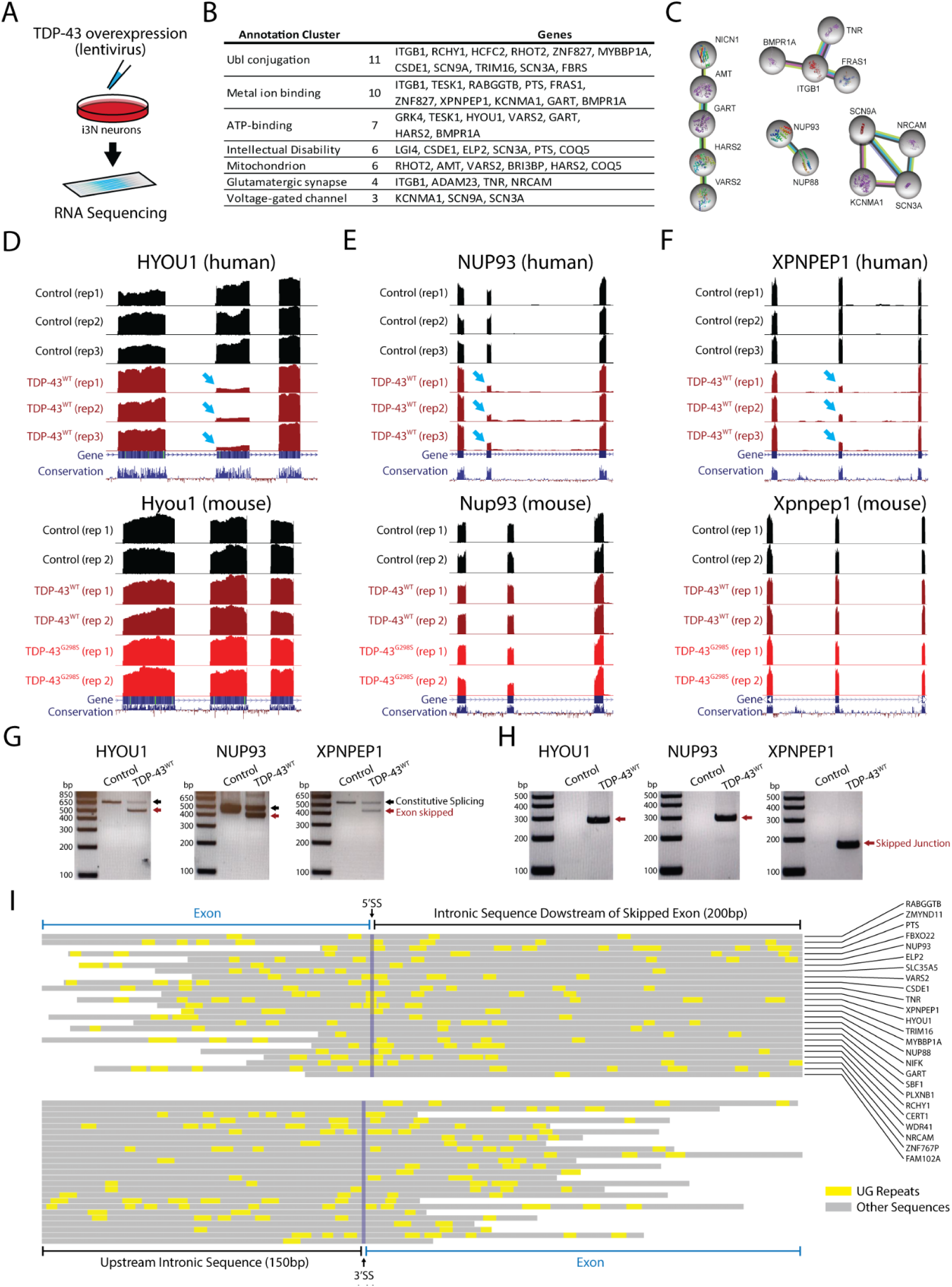
TDP-43 overexpression induces exon repression in humans. (**A**) To investigate whether exon skipping events also occur in humans, we infected human iPS cells with a lentivirus expressing human TDP-43 and analyzed the resulting RNA-Seq dataset. (**B**) Our analysis revealed several genes with skipping events, including genes involved in several molecular pathways. (**C**) Network analysis identified genes related to intellectual disability, synaptic activity, and mitochondrial proteins. We identified genes that had exons with particularly high levels of exon skipping, including HYOU1, NUP93, and XPNPEP1 (arrows). (**D**-**F**) When we cross-referenced these results with samples from transgenic mice, we found that exons repressed in humans were not repressed in mice. We validated these findings using RT-PCR in i3Neurons. We used double band RT-PCR (**G**) with primers located in the adjacent exon to the repressed exon or single band RT-PCR (**H**) with primers spanning the skipped junction. (**I**) Analysis of UG repeats in 25 targets revealed that these TDP-43 recognition motives are found in both repressed exons and intronic sequences. UG motifs appear slightly more frequently around the downstream 5’ splice site, but with far shorter UG repeat lengths than those found adjacent to cryptic exons (*73*).

**Figure 3.**
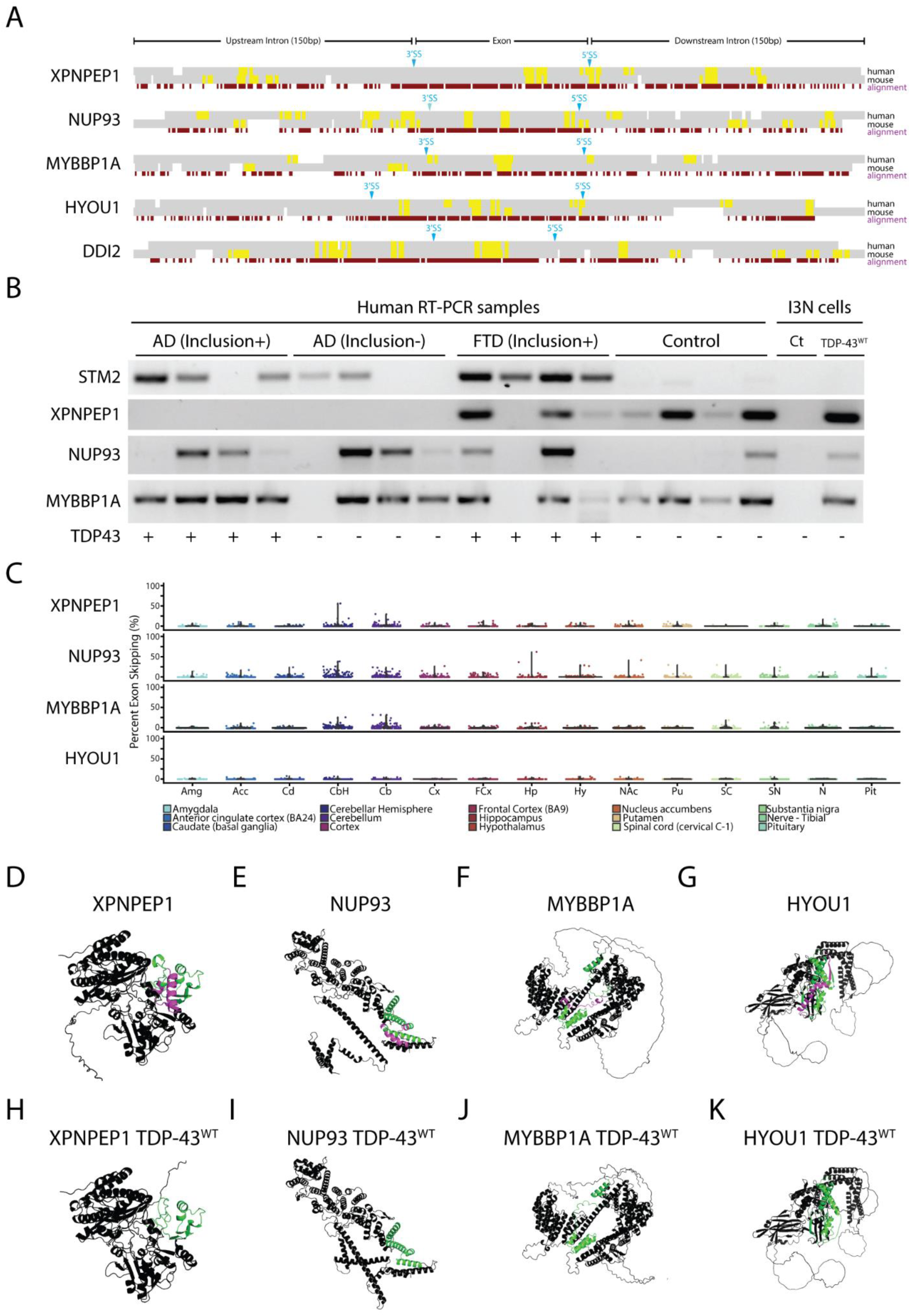
TDP-43 exon skipping events are found in aging human brains but do not correlate with disease. (**A**) Alignment of syntenic mouse (mm10) and human (hg38) genomic sequences surrounding exons repressed by TDP-43 overexpression in human cells. Constitutively spliced exons in the genes XPNPEP1, NUP93, MYBBP1A, and HYOU1 are skipped in human cells but not mouse cells when TDP-43 is overexpressed. By contrast, the exon in DDI2 is repressed in both mouse and human cells. An analysis of UG motifs (yellow highlights) reveals that slightly higher frequencies in UG repeat frequency surrounding the 3’ and 5’ splice sites may account for species-specific exon skipping. (**B**) We performed single-band RT-PCR to amplify cryptic junctions or exon-repressed junctions in human brain samples from patients with AD pathology with or without TDP-43 inclusions, frontotemporal dementia with inclusions, and control patients who did not have TDP-43 inclusions. As previously reported, the cryptic exon in the gene STMN2 is highly correlated with TDP-43 proteinopathy (*16, 17*). RT-PCR analysis showed that exon skipping occurred in both control and disease samples, indicating that aberrant exon skipping does not correlate with TDP-43 proteinopathy. (**C**) Since aging is a primary factor for developing neurodegenerative disease, we wanted to explore whether skipping events appear normally in the CNS and whether skipping events correlated with aging. We analyzed publicly available human RNA-Seq datasets (GTEx) and measured the percent spliced-in values of skipping events in patients aged between 60 to 69 years old. Exon skipping was found at low levels in most of the different brain areas analyzed, with slightly higher levels in the cerebellum. Using AlphaFold 2, we modeled protein structures with (**D**-**G**) and without (**H**-**K**) exons repressed by TDP-43 overexpression. Purple highlights indicate the repressed exons while green highlights indicate flanking amino acid sequences. TDP-43 induced exon skipping can dramatically influence protein structure and may lead to functional deficits.

Next, we wanted to explore whether TDP-43-mediated exon skipping represents a potential mechanism underlying the pathogenesis of ALS and TDP-43-related dementias. To study this, we designed single-band RT-PCR primers that could selectively amplify the unique junction produced by constitutive exon skipping. We profiled brain samples from individuals diagnosed with frontotemporal dementia (FTD), Alzheimer’s disease (AD) with or without TDP-43 inclusions, TDP-43 negative human controls, and i3Neurons. Previous studies have demonstrated that a cryptic exon found in the gene *STMN2* is a robust biomarker for ALS-FTD (*16, 18, 53*). As expected, the *STMN2* cryptic exon was detected in all four FTD-TDP-43 human samples and in 3 out of 4 AD TDP-43 positive samples (Fig. 3B). However, skipping events in at least three genes (*HYOU1*, *NUP93*, and *XPNPEP1*) did not exhibit a correlation with disease (Fig. 3B) and could be detected in both disease cases and controls.

Since exon skipping was detected in some control cases, we explored whether aberrant exon skipping could be correlated with aging. To study this, we explored publicly available RNA-Seq archives using the Snaptron search engine (*54*), queried each skipped exon across thousands of brain samples available through the GTEx repository (*55*), and generally observed low levels of exon skipping, with slightly higher skipping frequencies in cerebellum tissue (Fig. 3C, Supplementary Fig. 4). Our data suggest that aberrant exon skipping, induced by excessive levels of nuclear TDP-43, is unlikely to play a significant role in disease pathogenesis.

Since there is no significant correlation between aberrant exon skipping and disease, it will be important to avoid exon skipping when modeling TDP-43 proteinopathy. Overexpression of TDP-43 can be a valuable tool for investigating cytoplasmic toxicity, gain-of-function mechanisms, and post-translational modifications that may impact TDP-43 binding and other pathological interactions (*56–59*). However, dose-dependent toxicity resulting from exon skipping may confound results when analyzing phenotypes from these model systems, particularly since TDP-43 induced exon skipping appears to be highly species specific. Using the AlphaFold 2 protein prediction artificial intelligence system (*60*), we investigated the consequences on protein structure due to exon skipping and found significant impacts on protein structure and folding. For example, the XPNPEP1 isoform that results from exon skipping leads to complete disruption of the second domain of the protein (Fig. 3D,H). For HYOU1, we observe a loss of alpha helical domains in the C-terminal region of the protein (Fig. 3G, K). In NUP93, we predict that exon skipping would lead to dramatic changes in the C-terminal region that is crucial for nuclear pore complex assembly (Fig. 3E, I). These results suggest that exon skipping can dramatically affect protein structure and function, thereby leading to cellular toxicity.

A promising alternative to overexpression of wildtype TDP-43 (TDP-43^WT^) that could avoid exon skipping is the overexpression of TDP-43^NLSm^, where the nuclear localization signal is mutated (*46, 48, 61*). Transgenic mice (rNLS8) have been generated where expression of TDP-43^NLSm^ can be induced by doxycycline (Dox) under the control of a neuron-specific driver line, the human NEFH-tTA promoter (*49*). However, since TDP-43^NLSm^ can enter the nucleus passively (*62, 63*), we wanted to determine the maximum level of TDP-43^NLSm^ expression at which exon skipping can be detected. For this we used previously established Dox-inducible HEK-293 stable cell lines (*64*) to conditionally express TDP-43^WT^ (iGFP-WT) or TDP-43^NLSm^ (iGFP-NLSm) (Fig. 4A). Using RT-PCR, we found that exon skipping was present following doxycycline-dependent induction of either TDP43^WT^ or TDP43^NLSm^ Dox induction. However, the percentage of exon skipping for iGFP-NLSm induction reached only approximately 9% while iGFP-WT induction led to exon skipping of ∼90% (Fig. 4B). As previously described, induction of either iGFP-WT or iGFP-NLSm causes autoregulation of endogenous TDP-43, but also increases the total amount of TDP-43 protein (Fig. 4C-D). We estimate that a 50% increase in total TDP-43 compared to its endogenous level is enough to induce at least 40% of exon skipping. By comparison, a 100% increase in total TDP-43, due to iGFP-NLSm induction, leads to only 9% exon skipping (Fig. 4E).

**Figure 4.**
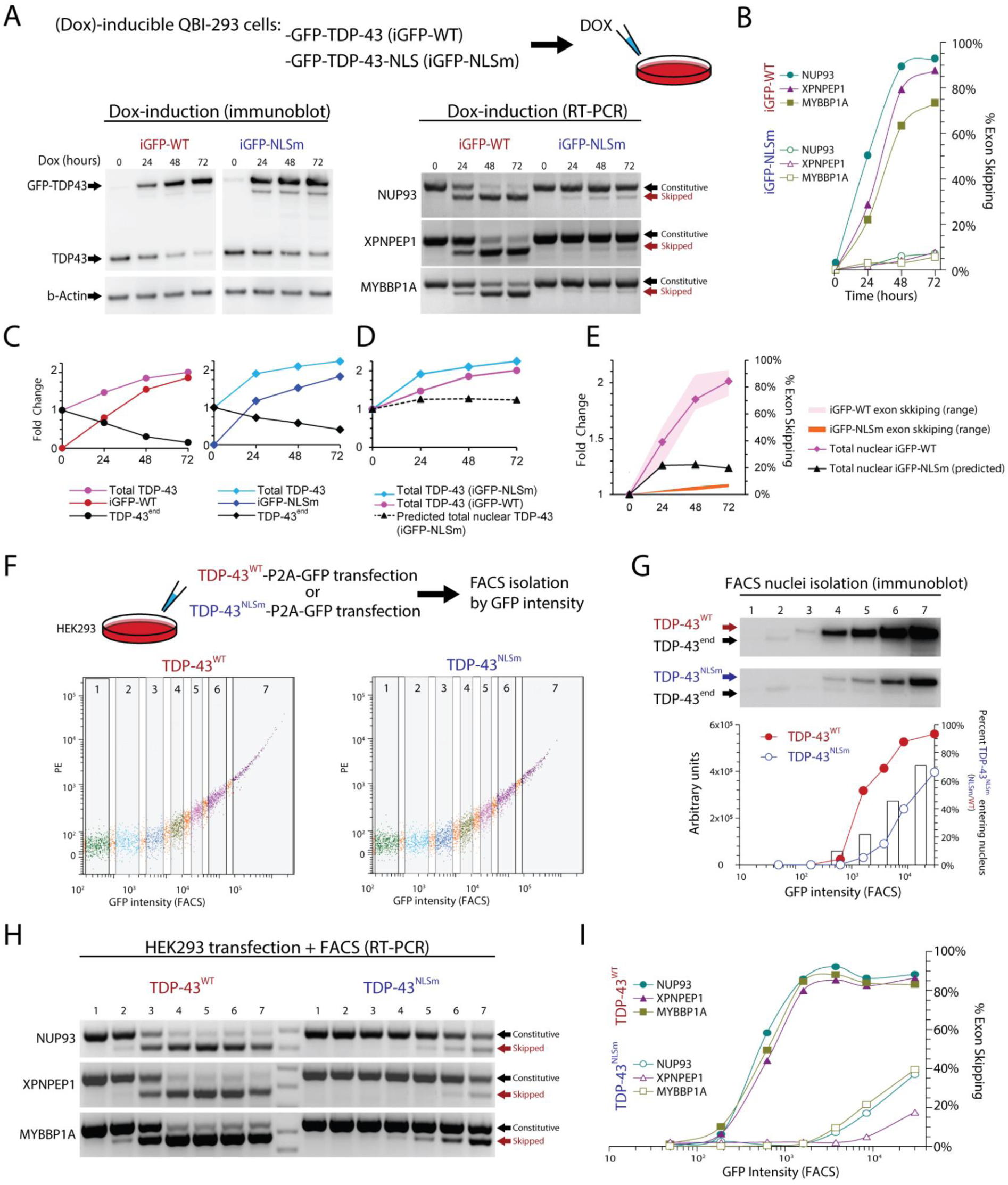
Overexpression of the ΔNLS mutant TDP-43 (TDP-43^NLSm^) induces exon repression when expressed at higher levels than wildtype TDP-43. To mimic mislocalization of TDP-43 into the cytoplasm, we used mutant forms of TDP-43. We wanted to explore to what extent this studied protein can passively migrate into the nucleus and repress exons. We used QBI-293 cells that have a stable cassette with TDP-43 tagged with GFP (iGFP-WT) and a TDP-43 mutated form in its nuclear localization signal (iGFP-NLSm), both lines under the control of the Doxycycline (Dox) inducible promoter. Cultures were exposed to Dox for different times (0, 24, 48, 72 hours). (A) Exon skipping events were seen when wild type or NLS TDP-43 forms were induced. (B) Quantification from RT-PCR results in iGFP-NLSm Dox induction shows that exon repression only reached ∼9% exon skipping compared to ∼95% in iGFP-WT. (C) Protein quantification shows TDP-43 endogenous self-regulation and total levels of TDP-43 for both WT and NLSm reached approximately ∼2 times the normal levels of TDP-43. (D and E) Approximate levels of total nuclear TDP-43 based on estimates of TDP-43^NLSm^ passive diffusion into the nucleus. (F) To evaluate to what extent TDP-43^NLSm^ was passively diffusing in the nucleus, we transfected HEK-293 cells with TDP-43^WT^-P2A-GFP or TDP-43^NLSm^-P2A-GFP under the control of a constitutive promoter and FACS isolated seven fractions according to GFP intensity. (G) We isolated nuclei from all seven fractions and detected TDP-43 by immunoblotting, TDP-43 endogenous is reduced while TDP-43^WT^ or TDP-43^NLSm^ are increasing. Raw intensity value in arbitrary units is plotted and the bar graph shows the ratio between overexpressed TDP-43^NLSm^ and TDP-43^WT^, which suggest that TDP-43^NLSm^ passive diffusion rate to the nucleus increases as the expression of this protein also progressively increases. (H) Double band RT-PCR from whole cell fractions showed the progressive increase of exon skipping when either TDP-43^WT^ or TDP-43^NLSm^ are overexpressed. (I) TDP-43^WT^ can almost completely repress exons evaluated (∼95%), however, TDP-43^NLSm^ only reached up to ∼40% of exon repression. Using these data, we estimate the proportion of TDP-43^NLSm^ in the Dox-inducible system, dotted line (D). Levels of total TDP-43^WT^ and total predicted TDP-43^NLSm^ in the nucleus are plotted together with their respective exon expression levels (E).

In animal models, first month of doxycycline induction results in overexpression of about 8 to 10-fold total TDP-43 (TDP-43^NLSm^ + endogenous TDP-43), suggesting that excessive levels of TDP-43^NLSm^ may induce higher levels of exon skipping (*65*). To test whether higher levels of TDP-43^NLSm^ overexpression promotes higher passive diffusion into the nucleus, we transfected HEK-293 cells with TDP-43^WT^-P2A-GFP or TDP-43^NLSm^-P2A-GFP and used FACS to obtain seven population fractions based on GFP intensity (Fig. 4F). We performed nuclei isolation on each fraction and quantified TDP-43 levels by immunoblotting and found that endogenous TDP-43 (TDP-43^end^) was eliminated by autoregulation from TDP-43^WT^ transfection in FACS populations with higher GFP intensity. This autoregulation was less evident in TDP-43^NLSm^, as nuclear TDP-43^NLSm^ did not reach levels that were sufficient to completely repress TDP-43^end^ even in the highest GFP intensity population. By comparing the ratio of TDP-43^WT^/TDP-43^NLSm^ at equivalent FACS populations, we estimated that 10-45% of TDP-43^NLSm^ can enter the nucleus. The wide range in diffusion percentages reflects the non-linear increase in TDP-43^WT^ transfection, as total TDP-43 levels begin to saturate and plateau despite an increase in GFP fluorescence (Fig. 4G). To estimate the minimal level of TDP-43 required to induce exon skipping, we also isolated RNA from each of the seven FACS populations isolated by GFP intensity and evaluated exon skipping using 2-band RT-PCR (Fig. 4H). We found that TDP-43^WT^ overexpression induced 90% exon skipping while TDP-43^NLSm^ saturated at 40% exon skipping (Fig 4H). Since TDP-43^NLSm^ induces exon skipping in FACS fraction six at equivalent levels as the iGFP-NLSm Dox inducible system, i.e. both induce 9% exon skipping, we can use immunoblot analysis of isolated nuclei to estimate that nuclear TDP-43^NLSm^ is approximately 40% of nuclear TDP-43^WT^ (Fig. 4G). With these estimates, we extrapolated that total TDP-43 levels 1.1 to 1.5-fold above normal can begin to induce exon skipping. Together this suggests that cell line and animal models using TDP-43^NLSm^ could cause exon skipping toxicity if expression reaches sufficiently high levels. Estimations on the levels of TDP-43 required to induce exon skipping and the diffusion of TDP-43^NLSm^ are summarized in Table 1.

**Table 1.**
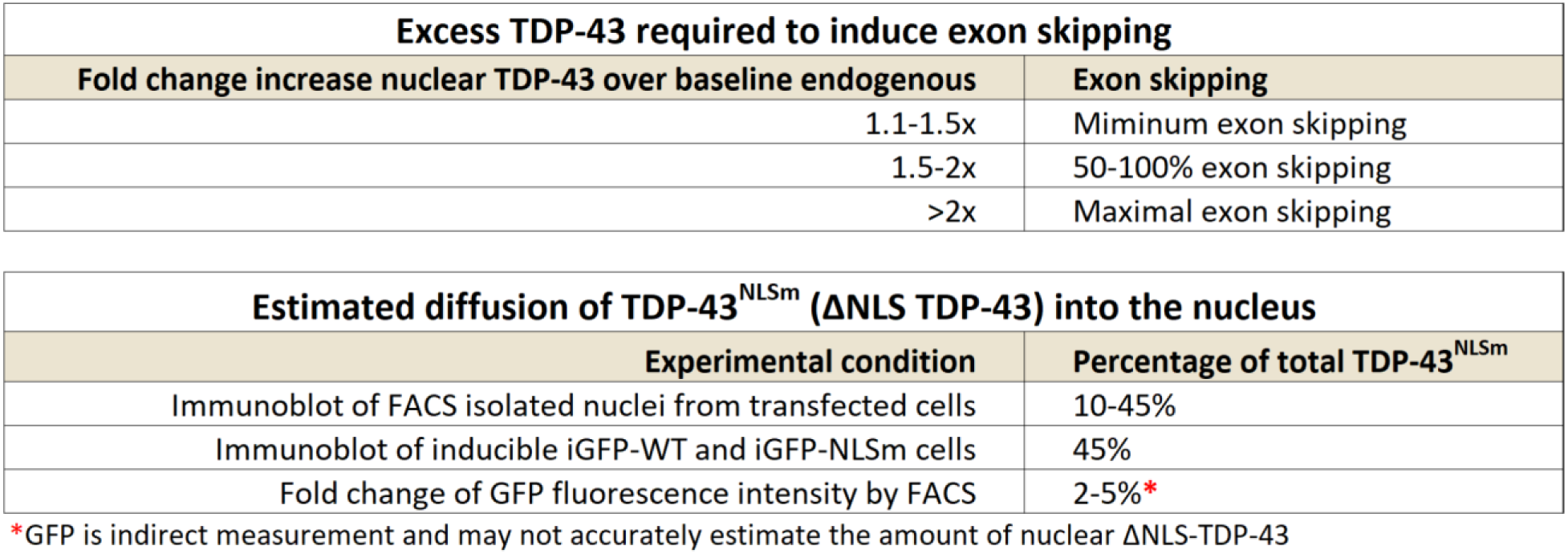
Estimates for the amount of excess nuclear TDP-43 that is required to induce exon skipping across different experimental conditions. Our results suggest that an increase in TDP-43 that is greater than 1.1 to 1.5 fold over normal protein levels may lead to exon skipping toxicity. Models for TDP-43 overexpression should avoid increasing nuclear TDP-43 and aim to strictly limit excess TDP-43 to only the cytoplasm.

| Excess TDP-43 required to induce exon skipping |  |
| --- | --- |
| Fold change increase nuclear TDP-43 over baseline endogenous | Exon skipping |
| 1.1-1.5x | Minimum exon skipping |
| 1.5-2x | 50-100% exon skipping |
| >2x | Maximal exon skipping |

**Table 1. Estimates for the amount of excess nuclear TDP-43 that is required to induce exon skipping across different experimental conditions.**
| Estimated diffusion of TDP-43<sup>NLSm</sup> (ΔNLS TDP-43) into the nucleus | |
| --- | --- |
| Experimental condition | Percentage of total TDP-43<sup>NLSm</sup> |
| Immunoblot of FACS isolated nuclei from transfected cells | 10-45% |
| Immunoblot of inducible iGFP-WT and iGFP-NLSm cells | 45% |
| Fold change of GFP fluorescence intensity by FACS | 2-5%* |

\*GFP is indirect measurement and may not accurately estimate the amount of nuclear ΔNLS-TDP-43

## Discussion

In this study, we explore the molecular mechanisms by which excessive levels of TDP-43 in the nucleus can induce cellular toxicity. Our findings reveal that TDP-43 overexpression leads to skipping of constitutive exons that are normally incorporated into mRNAs under steady state conditions. This aberrant exon skipping appears to be driven by high concentrations of nuclear TDP-43 binding to suboptimal UG motifs located near these constitutive exons, although further studies are necessary to understand the exact mechanism of splicing repression.

Indeed, we observe that nearly all exons skipped due to TDP-43 overexpression are species-specific, at least when comparing between TDP-43 overexpression in mouse and human neurons. Moreover, while aberrant constitutive exon skipping could be detected in some human brain samples, there was no clear correlation with disease status. By contrast, cryptic exons were exclusively detected in human disease samples, indicating their potential significance in disease pathogenesis. These findings have important implications for the interpretation and use of TDP-43 overexpression models in neurodegenerative disease research.

Our study underscores the need for caution when interpreting data obtained from TDP-43 overexpression models. We believe that TDP-43 induced exon skipping can be avoided when generating such models by ensuring that overexpressed TDP-43 is restricted to the cytoplasm and limiting nuclear TDP-43 protein levels to 130% to 150% above steady state. Cytoplasmic gain of function and nuclear loss of function of TDP-43 may both contribute to the pathogenesis of neurodegenerative diseases, but toxicity due to exon skipping may confound the interpretation of some model systems. For example, we observe minimal exon skipping in cell lines that stably express ΔNLS-TDP-43, but rNLS8 mice exhibit levels of nuclear TDP-43 that may induce increased exon skipping. Likewise, it remains to be determined whether AAV-mediated delivery of TDP-43 can also induce exon skipping, or whether TDP-43 autoregulation can minimize excessive nuclear TDP-43.

In conclusion, our study provides new insights into the complex molecular mechanisms underlying TDP-43 gain- and loss-of-function models. Our findings suggest that TDP-43 autoregulation is a highly conserved mechanism because both reduction and increases in TDP-43 protein levels lead to toxicity; depletion of nuclear TDP-43 leads to cryptic exon incorporation, while excess nuclear TDP-43 leads to constitutive exon skipping (Fig. 5). We believe that future studies that control for aberrant exon skipping may further refine gain-of-function toxicities due to excess cytoplasmic TDP-43. Understanding the nuances of TDP-43 toxicity that are specific to various subcellular compartments will be critical for advancing therapeutic approaches to treat neurodegenerative diseases.

**Fig 5.**
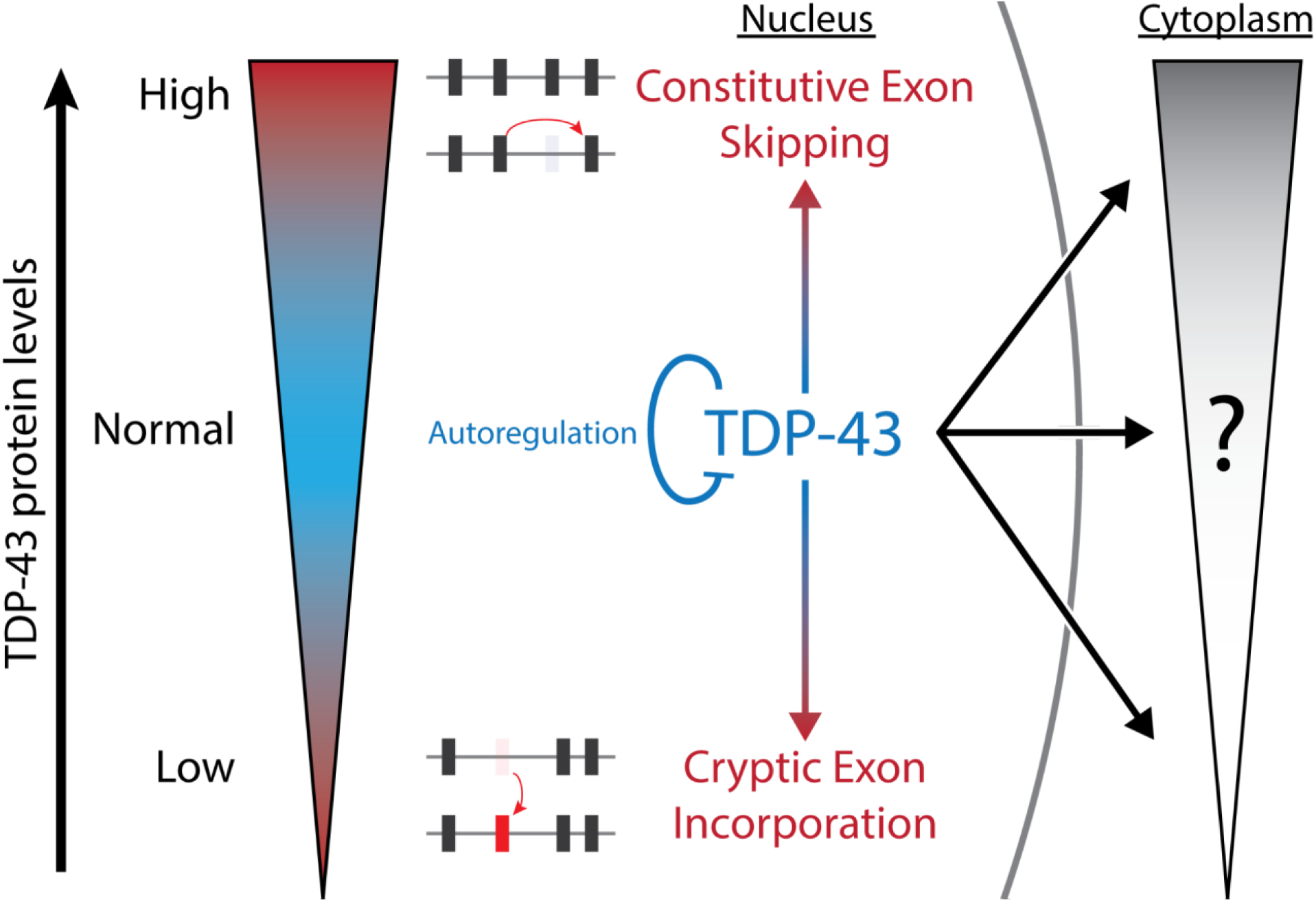
Summary Diagram. TDP-43 is a highly autoregulated protein due to different forms of cellular toxicity when TDP-43 protein levels are either too low (cryptic exon incorporation) or too high (aberrant exon skipping). Our study has demonstrated that these splicing deficits are linked specifically to nuclear TDP-43, whereas toxicity due to cytoplasmic TDP-43 remains to be fully elucidated. Future studies that avoid constitutive exon skipping may identify biomarkers for cytoplasmic-specific TDP-43 toxicity.

## Methods

### Ethics Declarations

The authors declare no competing interests.

### Antibodies

The following antibodies were used for mouse tissues immunohistochemistry and immunofluorescence: Mouse anti-Human TDP-43 (Novus 2E2-D3); Rabbit anti-Mouse TDP-43 N-Term (Proteintech 10782-2); Rabbit anti-Mouse TDP-43 C-Term (Proteintech 12892-1);Rabbit anti-Synaptophysin (Life Technologies Z66, 1:500); Mouse anti-Pan-Axonal neruofilament (Covance, SMI-312R, 1:1000). The following antibodies were used for cortex and spinal cord immunoblot: Mouse anti-Human TDP-43 (Novus 2E2-D3); Rabbit anti-Mouse TDP-43 N-Term (Proteintech 10782-2); Rabbit anti-Mouse TDP-43 C-Term (Proteintech 12892-1); Mouse anti-GAPDH (Sigma, GAPDH-71.1).The following antibodies were used for i3Neuron stable cell lines immunoblot: Rabbit anti-TDP-43 (Proteintech 10782-2-AP 1:1000); Mouse anti-FLAG (Sigma F1804, 1:5000); Rabbit anti-GAPDH (CST 2118, 1:1000); Mouse anti-Tubulin (Thermofisher 14-4502-82, 1:1000). The following antibodies were used for QBI-293 stable cell lines immunoblot: Mouse anti-TDP-43 (abcam ab104223, 1:2000); Rabbit anti-GFP (Cell signalling mAb 2956, 1:1000); Mouse anti-Beta Actin (Proteintech 66009-1, 1:2000). The following antibody was used for HEK-293 cells immunoblot: Rabbit anti-Mouse TDP-43 N-Term (Proteintech 10782-2, 1:2000).

### Transgenic mouse generation

A Thy1.2 expression cassette on a pUC18 backbone was used to generate transgenic mice. Plasmids pTSC-TDP-WT and pTSC-TDP-G298S were constructed and submitted to the Transgenic Core Laboratory at the Johns Hopkins University School of Medicine for pronuclear injection using the hybrid mouse strain C57B6;SJL. Potential founders were screened by tail cutting, genomic DNA extraction, and PCR. The WT and G298S lines were maintained in the hybrid background and phenotypic characterization was started at the F3 generation. All mouse experiments were approved by the Johns Hopkins University Animal Care and Use Committee.

### Hanging wire tests

F3 generation was followed to the end-stage (death or inability to right from a lateral decubitus position), and survival data was recorded. A subset of this population was selected for monthly hanging wire tests. Briefly, a mouse was placed on a wire grid, and the grid was shaken to encourage the mouse to grip the wires. The grid was then turned upside down and held level while the time for the mouse to fall was recorded, the endpoint was 60 sec. Each mouse was tested 3 times, with the maximum hang time recorded as the result.

### Tissue Preparation for Sectioning

Mice were anesthetized with an intraperitoneal injection of 15% chloral hydrate and anesthesia was monitored with limb and corneal reflex checks. Portions of quadriceps muscles were snap-frozen for sectioning. Mice were transcardially perfused with 50-100 mL of cold PBS, and then fixed on 4% PFA. The L3 and L4 dorsal root ganglia and attached dorsal and ventral roots were identified, removed, placed in a fixation buffer with 2% glutaraldehyde overnight, washed in PBS and embedded in Epon for EM. The brain and spinal cord were dissected and separated into right/left halves. One half of each was placed in a fixation buffer overnight at 4°C with gentle agitation, and then held in PBS prior to embedding in paraffin and sectioning. The other half brain was half spinal cord was placed in a fixation buffer for 2 hours, and then switched to sterile PBS with 30% sucrose overnight at 4°C to be embedded in O.C.T and frozen in isopentane for cryostat sectioning.

### Immunofluorescence and Immunohistochemistry

Paraffin-embedded brain samples were sectioned in the sagittal axis and spinal cord samples were sectioned in the cross-sectional axis onto slides. Slides used for immunohistochemistry were incubated at 60°C for 30 minutes, then deparaffinized in xylene and ethanol. Antigen retrieval was accomplished by incubating slides for 5 minutes in boiling 10mM sodium citrate buffer (pH 6.0). Slides were washed with PBS and blocked and stained using appropriate primary antibodies and reagents from the Vectastain Elite ABC kit (Vector Labs). Diaminobenzidine exposure was titrated to optimal contrast, sections were counterstained with Mayer’s hemalaun, and then slides were dehydrated using ethanol and xylene and mounted.

Frozen gastrocnemius muscle was cut into 40μm longitudinal sections using a freezing sliding microtome (Leica). Sections were separated into wells on a 12-well plate and blocked in IF blocking buffer: PBS with 5% normal goat serum and 0.5% Triton-X, slides were incubated overnight at 4°C on primary antibody (rabbit anti-Synaptophysin and mouse anti-SMI-312), diluted in blocking solution. Sections were then washed with PBS with 0.5% Triton-X, incubated with α-bungarotoxin and goat anti-rabbit and anti-mouse secondary antibodies conjugated to Alexa Fluor-488 (Invitrogen), and washed again. Sections were spread on slides and coverslips were attached using Prolong Gold Antifade Reagent (Life Technologies). Slides were examined using a Zeiss LSM 510 Meta confocal microscope.

### Muscle Histology

Frozen quadriceps muscle embedded in O.C.T. were cut into 10μm cross-sections onto slides using a cryostat (Leica). For H&E staining of muscle, sections were covered for 20 minutes with a fixation buffer (4% PFA in PBS). Sections were rinsed with distilled water, stained with Mayer’s Hemalaun, rinsed, and stained with Eosin. After a final rinse, samples were dehydrated in ethanol and xylene, and coverslips were applied. For esterase staining, 25% α-naphthyl acetate, 5% acetone, 0.1% Pararosaniline HCL, and 0.1% Sodium Nitrate in 0.2M Sodium Phosphate solution was used. The solution was applied to quadriceps sections for 5 minutes, rinsed in running tap water for several minutes and slides were then dehydrated in ethanol and xylene, mounted. Esterase and H&E-stained sections were analyzed under light microscopy.

### I3Neurons TDP-43 lentivirus transduction

i3Neurons were transduced with lentivirus containing N-terminal Flag-tagged wild type TDP-43 at 1MOI, 2MOI, and 4MOI, respectively, at Brain physiological stage day 11. Neurons were harvested on Day 14 and dried ice frozen. Total RNA was isolated using Trizol extraction and used for downstream RNA sequencing analyses.

### TDP-43 Dox-inducible QBI-293 stable cell lines

iGFP-WT and iGFP-NLSm inducible cell lines were kindly provided by Silvia Porta and Virginia Lee (*48*) and cultured on Dulbecco’s Modified Eagle Medium supplemented with 10% FBS (Corning, 35-010-CV), and 1% Penicillin-Streptomycin (ThermoFisher Scientific, 15070063), L-glutamine (20 mM, Corning Cellgro, Manassas, VA) with G418 (400 µg ml−1, Calbiochem, La Jolla, CA). Cells were induced with Dox ug/mL and collected after 24h, 48h or 72h of induction.

### HEK-293 cell culture, transfection, FACS separation and nuclei isolation

HEK-293 cells were cultured in Dulbecco’s Modified Eagle Medium supplemented with 10% FBS (Corning, 35-010-CV), 1x GlutaMAX (ThermoFisher Scientific, 35050061), and 1% Penicillin-Streptomycin (ThermoFisher Scientific, 15070063). For overexpression of TDP-43^WT^ and TDP-43^NLS^ ORF expression cassettes were cloned into pAAV-CBh-mKate2-IRES-MCS (a gift from Marcella Patrick, Addgene plasmid # 105921) and transfected on HEK-294 cells with Lipofectamine 3000 (Thermo Fisher Scientific, L3000-008) following the manufacturer’s protocol. Two days of transfection single cell suspensions were obtained using TrypLE (ThermoFisher Scientific, 12604013) and sorted by GFP fluorescence intensity on a BD FACSCalibur in the JHMI Ross Flow Cytometry Core Facility. Nuclei were isolated following the 10x Genomics nuclei isolation protocol from cell suspensions.

### Immunoblotting

Tissues and cells were digested in RIPA Buffer with 1%, protease inhibitor (Roche Complete ULTRA mini tablet + EDTA), and phosphatase inhibitor (Roche PhosStop). Samples were centrifuged, and supernatants were saved. Protein concentrations in the supernatants were determined using a BCA assay (Pierce), and 20μg of total protein was loaded into each well of a 10-20% Tris-Glycine gel or a 4-12% Bis-Tris gel (Invitrogen). Protein was transferred to a PVDF membrane (Millipore), blocked with 5% BSA in TBS with 0.1% Tween-20. Membranes were incubated overnight at 4°C with primary antibody diluted in blocking buffer, then washed three times with TBS with 0.1% Tween-20, and incubated for 2 hours with secondary antibody (Goat anti-mouse IgG-HRP or Goat anti-rabbit IgG-HRP, Sigma) diluted in blocking buffer. Three more washes with TBS with 0.1% Tween-20 followed, and then membranes were soaked in ECL solution (EMD Millipore Immobilon), dried, imaged on a Biorad image. Densitometric analysis was performed using Quantity One software (Bio Rad).

### RNA extraction, library preparation, and RNA sequencing

Two ventral halves of spinal cords from TDP-43^WT^ and TDP-43^G298S^ transgenic animals and one littermate control for each line were dissected and transferred immediately into RNAlater storage reagent (Life Technologies). Total RNA from spinal cord tissues was extracted using the RNeasy Mini Kit (Qiagen). Total RNA from human tissue, QBI-293 stable cell lines, and HEK-293 cells were extracted using Monarch Total RNA Miniprep Kit (New England BioLabs, T2010S). To prepare RNA-Seq libraries, the TruSeq Stranded Total RNA Library Prep Kit (Illumina) was employed. Subsequently, the sample libraries were sequenced on an Illumina HiSeq for spinal cord samples and NextSeq for cultured cells. The obtained data was transformed into FASTQ files after demultiplexing.

### RT-PCR and Primers

Mouse animals were genotyped using the following primers: 5’-CGGAAGACGATGGGACGG TG, 5’-GCCAAACCCCTTTGAATGACCA, and 5’-AAGATGGCACGGAAGTCTAACCATG, was used to generate a 386 base pair band (spanning TARDBP exons 2, 3, and 4) in transgenic mice only and a 241 base pair internal control band in all mice. LunaScript RT SuperMix Kit was used for cDNA synthesis. The following primers were used for double band PCR in mouse tissue: Ddi2-F1: CAGAGTGTGCTCGTTTGGCA, Ddi2-R1: GACTCGTCGGGCTACCAAC, product: 284bp (Full) or 222bp (Skip); Mrpl45-F1: ACACTGTTTTCCGGACATGGT, Mrpl45-R1: TCGTACTCCTCCCAGGGTTT, product: 384bp (Full) or 210bp (Skip); Psmd14-F1: GAGCCAGGTCCTTGTTGAGT, Psmd14-R1: TTGGCTTGGAACACTGGATCA, product: 473bp (Full) or 353bp (Skip). The following primers were used for double band PCR in human cells: HYOU1-F2: TTCTATGACATGGGCTCAGGC, HYOU1-R2: ACTGCATCTCGGACGACAAA, product: 635bp (Full) or 500bp (Skip); NUP93-F2: GAGGCTGAAGAACATGGCAC, NUP93-R2: TCCCACAAAGCATGGCACTT, product: 496bp (Full) or 411bp (Skip); XPNPEP1-F2: GTTGGTGTGGACCCCTTGAT, XPNPEP1-R2: GACCCACACCTTCTCCCTTG, product: 522bp (Full) or 426bp (Skip); MYBBP1A-F1: AAAGTCTGGGAGAGAAGCCC, MYBBP1A-R1: CGCAGAGCCTTCTCCTTCTG, product: 586bp (Full) or 499bp (Skip). The following primers were used for single band PCR in human cells and brain samples: STM-F1: CTGCACATCCCTACAATGGC, STM-R1: CACAAGCCGCATTCACATTCA, product: 167bp; HYOU1-F1: CCGTATGCACCATTGTGACC, HYOU1-S5: CTGCTCAATCTCATCCTGTGC, product: 290bp; NUP93-S2: GGTCATATTGATAGAGCTTTTGATATCAGG, NUP93-R1: TGTACTGACAGTGTGCCGAC, product: 304bp; XPNPEP1-S3: GCCTGGATTACACAGGGCTATTT, XPNPEP1-R1: ATTCGGCTTCCAGACCCAAG, product: 177bp; MYBBP1A-S2: CTGCAGCTAATTCTGGATGACAAG, MYBBP1A-R1: CGCAGAGCCTTCTCCTTCTG, product: 320bp. Differences in RT-PCR product sizes were resolved on 4% agarose gels.

### Percent of Splicing Analysis

FASTQ files were aligned using STAR (*66*) and a filtered splice junction file was processed to calculate percent of splicing (PSI) values following methodology described in ASCOT (*67*). Briefly, BAM files were converted into bigWig files using Megadepth (*68*). We calculated the 5’PSI or 3’as the ratio between the number of mapped splice junctions from one exon to another over the total amount of splice junctions including that exon the 5’or 3’splicing site. PSI values between controls and TDP-43 overexpression were compared and visualized using UCSC genome browser (*72*), examples of human skipping events are shown in the Supplementary Data file.

### Analysis of exon skipping in GTEx human samples

Splice junctions from the GTEx archive were queried using the Snaptron Web Services Interface and only brain, nerve and pituitary samples were collected (*55*). PSI values for skipped junctions were calculated as previously mentioned. We used the normalized quantification of transcripts per million (TPM) from the GTEx consortium (GTEx_Analysis_2017-06-05_v8_RSEMv1.3.0) to correlate PSI values and TDP-43 expression level (*55*). Plots from different brain regions were made using RStudio.

### Protein structure modeling

Wildtype mRNA sequences for XPNPEP1 (ENST00000502935.6), NUP93 (ENST00000308159.10), MYBBP1A (ENST00000254718.9) and HYOU1 (ENST00000617285.5), were obtained from the GENCODE database and protein sequences were identified by translating the mRNA sequences with or without the exon of interest (*69*). Wildtype structures for NUP93 and HYOU1were downloaded from the AlphaFold Protein Structure Database. All other structures were generated using the Alpha Fold Monomer v2.0 pipeline (version 2.2.0) from the amino acid sequences on the Rockfish cluster at Johns Hopkins University(*70, 71*). PyMOL was used to visualize the structures (Schrödinger, LLC. The PyMOL Molecular Graphics System, Version 2.5.4, 2015)

### Statistical Analysis

Mouse data was analyzed using Microsoft Excel and GraphPad Prism software, with P values calculated using unpaired Student’s t tests. P values less than 0.05 were considered significant. All quantitative values were reported as mean ± standard deviation (SD).

### Data availability

RNA-Seq FASTQ files have been deposited in the NCBI Sequence Read Archive under accession number SRP######.

## Acknowledgements

We thank X. Zhang and the Johns Hopkins Ross Flow Cytometry Core Facility. We also thank L. Orzolek, T. Creamer, and the Johns Hopkins The Single Cell & Transcriptomics Core for sequencing services. We would like to thank Silvia Porta and Virginia Lee for kindly providing QBI-293 stable cell lines. Part of this work was carried out at the Advanced Research Computing at Hopkins (ARCH) core facility (rockfish.jhu.edu), supported by the National Science Foundation (NSF) grant number OAC 1920103. This work was supported by grants from the Alzheimer’s Association (J.P.L.), the Packard Center for ALS Research at Johns Hopkins to P.C.W., the ALS Association to P.C.W., the NIH (RF1MH123237 to S.B., P30AG066507 to J.C.T., U19AG033655 to J.C.T., R01NS123538 to L.R.H., R01NS107347 to S.S., RF1NS113820 to S.S., R21AG072078 to S.S., R01NS095969 to P.C.W.) and the National Science Foundation Graduate Research Fellowship Program to I.R.S. under Grant No. DGE2139757.

## Contributions

J.P.L. and R.P.C.O. conceived and oversaw all aspects of the study. R.P.C.O and J.P.L conceived analysis of exon repression and validation in human cells and mouse lines. W.T. and P.C.W. oversaw TDP-43^WT^ and TDP-43^G298S^ mouse line development and behavioral and pathological analyses. Y.Y. and S.S. oversaw human iPSC neuronal culture studies. L.R.H. assisted with Dox inducible QBI-293 cell culture. K.C. and J.C.T. assisted with pathological characterization of human brain tissues. V.T., W.C, and K.B., assisted with data analysis and human RT-PCR. I.R.S. assisted with protein structure predictions. S.B., L.R.H., S.S., and P.C.W. provided valuable feedback on experimental design and manuscript writing. R.P.C.O. and J.P.L. drafted the manuscript. All authors approved the final manuscript.

