## Supplementary Figures for "Elevated nuclear TDP-43 induces constitutive exon skipping"

**Affiliations:**
Departments of [1]Pathology, [2]Neuroscience, [3]Physiology, [4]Ophthalmology and [5]Neurology, Johns Hopkins School of Medicine, Baltimore, MD 21205

* indicates equal contribution
[#] indicates corresponding author

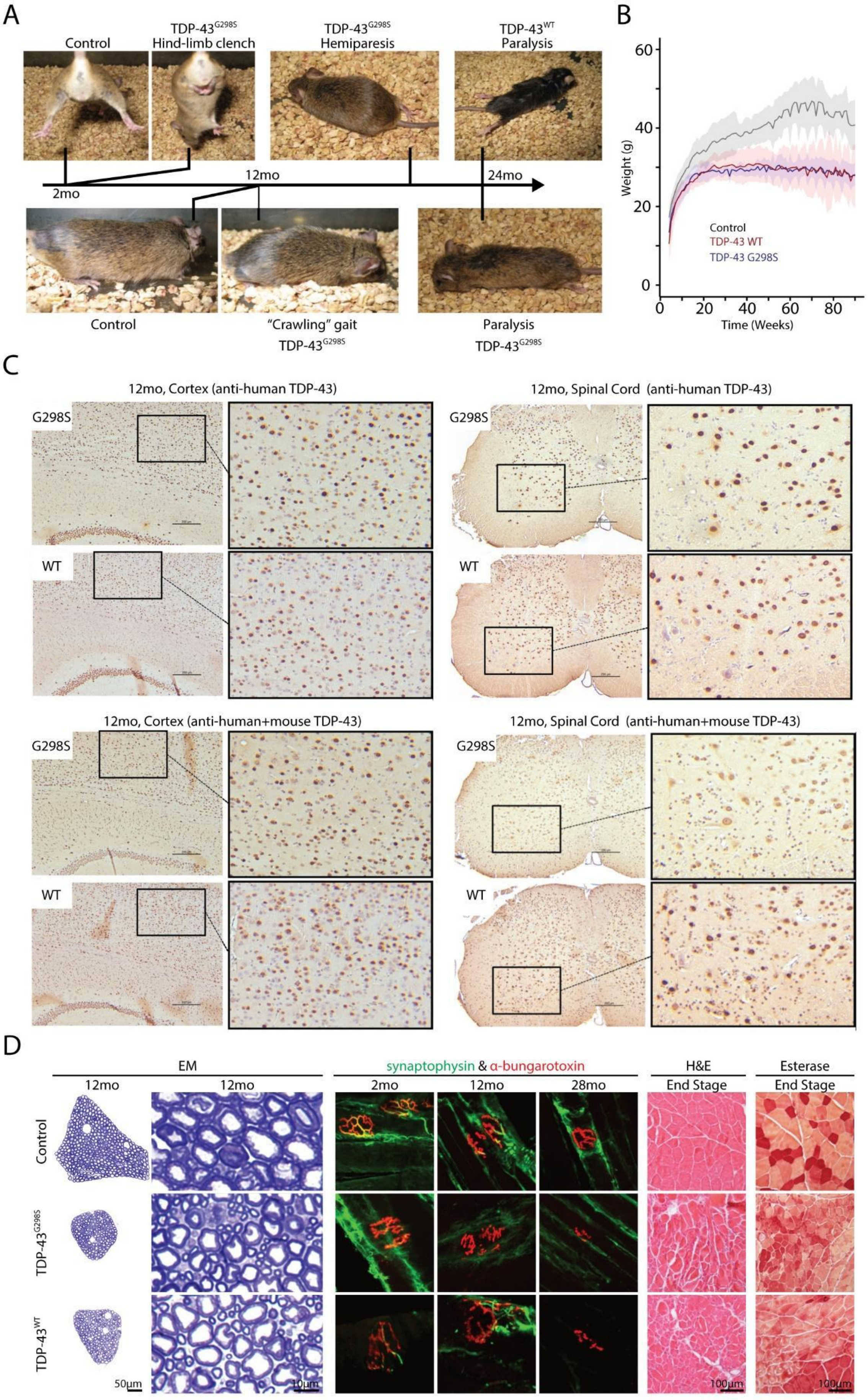

A
Control
TDP-43G298S Hind-limb clench
TDP-43G298S Hemiparesis
TDP-43WT Paralysis
2mo
12mo
24mo
Control
"Crawling" gait TDP-43G298S
Paralysis TDP-43G298S
B
Weight (g)
Time (Weeks)
Control
TDP-43 WT
TDP-43 G298S
C
12mo, Cortex (anti-human TDP-43)
12mo, Spinal Cord (anti-human TDP-43)
G298S
WT
12mo, Cortex (anti-human+mouse TDP-43)
12mo, Spinal Cord (anti-human+mouse TDP-43)
D
EM
synaptophysin & α-bungarotoxin
H&E End Stage
Esterase End Stage
12mo
2mo
28mo
Control
TDP-43G298S
TDP-43WT
50µm
10µm
100µm

**Supplementary Figure 1. Pathological findings in TDP-43$^{WT}$ and TDP-43$^{G298S}$ transgenic models.** (**A**) Animals exhibited marked motor disabilities starting with hind-limb clench at 2 months of age, progressing through a crawling gait at 12 months and continue with hemiparesis at 18 months, finally leading to end stage paralysis at 24 months. (**B**) Reduction in motor capabilities was accompanied by a decrease in weight compared to controls. Immunohistochemical images showed that TDP-43 was confined to the nucleus of cortical and spinal cord cells. No cytoplasmic inclusions or TDP-43 clearance was observed. (**C**) Immunohistochemistry using an antibody against human-specific TDP-43 (top panel) and immunohistochemistry using an antibody against both human and mouse TDP-43 (bottom panel). (**D**) In order to assess the lumbar L3 ventral roots of both wild-type (WT) and G298S transgenic mice at 12 months, Toluene-stained Epon thick sections were utilized. The L3 roots from transgenic mice exhibited smaller ventral roots than the non-transgenic animals. Further examination of the electron microscopy thick sections of the ventral roots in G298S and WT transgenic mice revealed a reduced number of large myelinated axons, resulting in the increased visibility of smaller axons. Staining of synaptophysin (green) and $\alpha$-bungarotoxin (red) on gastrocnemius muscles revealed a progressive denervation of neuromuscular junctions in both transgenic mice. While a significant number of junctions appeared well-innervated in the 2-month-old transgenic mice, denervated junctions were notably observed in the 12-month-old transgenic mice. At the end-stage of symptoms, transgenic mice experienced muscle degeneration to the extent that it was difficult to detect end plates by α-bungarotoxin-staining. We analyzed quadriceps muscle at their end-stage because the motor neurons innervating quadriceps muscle are in the L1 to L3 region of the spinal cord in mice. As the transgenic mice aged, H&E staining showed a progressively increased disorganization and angulation. Additionally, esterase staining indicated denervated fibers (darker staining) with the fiber muscle architecture completely altered in transgenic animals.

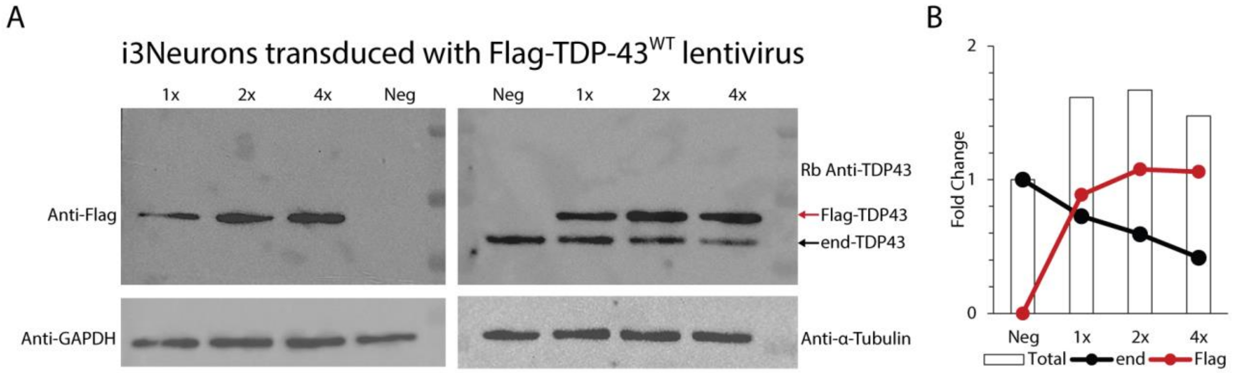


**Supplementary Figure 2. i3Neuron transduction with human TDP-43.** (**A**) i3Neurons were transduced with lentivirus containing an N-terminal Flag-tagged wild-type TDP-43 at 1MOI, 2MOI, and 4MOI, respectively. Consistent increase of exogenous TDP-43 (left) is seen accompanied with endogenous TDP-43 autoregulation (right). (**B**) Quantification of the right immunoblot, shows ~1.6 fold increase of total TDP-43 (endogenous TDP43 + transduced Flag-TDP43$^{WT}$).

A

### Mouse

| Gene | Control_1 | Control_2 | G298S_1 | G298S_2 | TDP43WT_1 | TDP43WT_2 |
|---|---|---|---|---|---|---|
| Arhgap44 | 89 | 78 | 31 | 28 | 13 | 8 |
| Ddi2 | 100 | 96 | 52 | 48 | 64 | 42 |
| Psme3 | 90 | 92 | 51 | 46 | 45 | 39 |
| Dgkq | 100 | 100 | 63 | 55 | 55 | 49 |
| Mrpl45 | 99 | 99 | 62 | 61 | 49 | 49 |
| Psmd14 | 100 | 100 | 58 | 62 | 53 | 60 |
| Hjurp | 96 | 95 | 49 | 44 | 75 | 64 |
| Cops4 | 90 | 85 | 54 | 56 | 52 | 49 |
| Tiam1 | 71 | 49 | 30 | 21 | 15 | 44 |
| Creld1 | 100 | 100 | 72 | 66 | 70 | 61 |
| Inpp4a | 67 | 77 | 45 | 41 | 37 | 38 |
| Shank1 | 59 | 57 | 27 | 27 | 31 | 27 |
| Pakap | 66 | 61 | 28 | 27 | 36 | 43 |
| Arhgef11 | 94 | 76 | 71 | 70 | 40 | 43 |
| Osbpl6 | 64 | 81 | 36 | 51 | 61 | 33 |
| Pik3cb | 93 | 100 | 72 | 80 | 71 | 54 |
| Pdzd4 | 78 | 82 | 59 | 43 | 51 | 62 |
| Dst | 37 | 28 | 4 | 8 | 9 | 3 |
| Herc2 | 100 | 91 | 76 | 80 | 64 | 58 |
| Golga4 | 40 | 33 | 14 | 10 | 11 | 7 |
| Stk24 | 56 | 50 | 42 | 23 | 21 | 21 |
| Ube3c | 100 | 97 | 75 | 76 | 75 | 64 |
| C530008M17Rik | 64 | 57 | 39 | 33 | 31 | 36 |
| Tub | 100 | 80 | 66 | 72 | 56 | 71 |
| Atp2b1 | 67 | 65 | 41 | 46 | 45 | 38 |

B

### Human

| Gene | CONTROL1 | CONTROL2 | CONTROL3 | TDP43WT1 | TDP43WT2 | TDP43WT3 |
|---|---|---|---|---|---|---|
| ADAM23 | 83 | 63 | 64 | 7 | 2 | 23 |
| AMT | 100 | 95 | 89 | 11 | 5 | 29 |
| ATP9B | 100 | 96 | 100 | -1 | 44 | 13 |
| BMPR1A | 100 | 100 | 100 | 20 | 19 | 26 |
| BRI3BP | 100 | 100 | 100 | 42 | 20 | 60 |
| CANX | 99 | 100 | 99 | 46 | 37 | 49 |
| CCDC126 | 100 | 79 | 100 | 26 | 36 | 44 |
| CERT1 | 70 | 73 | 72 | 17 | 10 | 20 |
| CLASP2 | 94 | 92 | 96 | 28 | 27 | 31 |
| COQ5 | 100 | 99 | 99 | 27 | 30 | 19 |
| CSDE1 | 98 | 96 | 97 | 10 | 8 | 11 |
| CSPP1 | 100 | 94 | 100 | 22 | 17 | 38 |
| DDI2 | 86 | 96 | 100 | 29 | 13 | 60 |
| DPY19L1P1 | -1 | 100 | 100 | 20 | -1 | 40 |
| DRGX | 93 | 87 | 100 | 42 | 29 | 45 |
| ELP1 | 88 | 88 | 98 | 21 | 31 | 56 |
| ELP2 | 96 | 96 | 97 | 27 | 29 | 41 |
| ENSG00000256591 | 100 | 100 | 100 | 40 | 40 | 42 |
| ERMAD | 100 | 100 | 100 | 27 | 31 | 47 |
| FAM102A | 100 | 100 | 95 | -1 | -1 | -1 |
| FAM66C | 73 | 53 | 76 | 0 | 22 | -1 |
| FANCG | 83 | 57 | 88 | 14 | 14 | 13 |
| FAT1 | 88 | 91 | 79 | 11 | 11 | 20 |
| FBRS | 100 | 98 | 98 | 34 | 25 | 39 |
| FBXO22 | 95 | 89 | 89 | 25 | 28 | 26 |
| FRAS1 | 100 | 100 | 100 | 29 | 38 | 30 |
| GART | 90 | 94 | 93 | 15 | 10 | 18 |
| HARS2 | 98 | 97 | 92 | 27 | 22 | 32 |
| HCFC2 | 100 | 88 | 100 | 31 | 17 | 18 |
| HECW1 | 59 | 73 | 75 | 24 | 8 | 9 |
| HIRA | 100 | 99 | 100 | 45 | 41 | 45 |
| HYOU1 | 100 | 100 | 100 | 18 | 16 | 21 |
| IL17RB | -1 | 91 | 71 | 0 | 2 | 0 |
| INSYN1 INSYN1-AS1 | -1 | 75 | 86 | 17 | 20 | 29 |
| INTS10 | 84 | 91 | 87 | 26 | 17 | 25 |
| ITGB1 | 100 | 100 | 100 | 7 | 5 | 14 |
| KCNMA1 | 74 | 59 | 66 | -1 | 0 | 7 |
| LGI4 | 88 | 85 | 97 | 27 | 20 | 50 |
| LINC00680-GUSBP4 | 100 | 92 | 69 | 4 | 0 | 0 |
| MYBBP1A | 91 | 86 | 95 | 14 | 14 | 10 |
| NEBL | -1 | 100 | 100 | 40 | -1 | 33 |
| NHLRC3 | -1 | 100 | 100 | -1 | -1 | -1 |
| NICN1 | 85 | 83 | 94 | 29 | 25 | 31 |
| NIFK | 90 | 97 | 98 | 22 | 44 | 52 |
| NIPAL3 | 59 | 86 | 91 | 2 | 0 | 6 |
| NQO2 | 100 | 100 | 100 | 9 | 10 | 44 |
| NRCAM | 91 | 86 | 93 | 32 | 24 | 27 |
| NUP88 | 100 | 93 | 91 | 35 | 36 | 25 |
| NUP93 | 100 | 99 | 100 | 32 | 28 | 34 |
| PLXNB1 | 95 | 97 | 97 | 27 | 21 | 21 |
| PPA1 | 100 | 96 | 99 | 29 | 29 | 37 |
| PTS | 91 | 86 | 92 | 16 | 11 | 17 |
| RABGGTB | 98 | 89 | 96 | 10 | 12 | 23 |
| RCHY1 | 94 | 91 | 93 | 24 | 17 | 13 |
| RHOT2 | 100 | 97 | 91 | 50 | 30 | 38 |
| RNASET2 | 100 | 71 | 89 | 22 | 50 | 20 |
| RNF114 | 97 | 94 | 95 | 33 | 17 | 31 |
| SBF1 | 81 | 81 | 76 | 15 | 8 | 14 |
| SCN3A | 100 | 99 | 100 | 51 | 29 | 31 |
| SCN9A | 100 | 100 | 99 | 33 | 32 | 55 |
| SEC61A2 | 100 | 100 | 99 | 41 | 39 | 37 |
| SESN3 | 100 | 100 | 97 | 53 | 25 | 47 |
| SLC17A5 | 100 | 86 | 80 | 11 | 11 | 27 |
| SLC35A5 | 100 | 100 | 100 | 32 | 40 | 26 |
| SPRK2 | 86 | 65 | 84 | 21 | 13 | 29 |
| STC2 | 100 | 100 | -1 | 23 | 29 | -1 |
| SUN1 | 100 | 96 | 95 | 0 | 4 | 0 |
| TESK1 | 100 | 94 | 99 | 34 | 26 | 45 |
| TLCD3A | 100 | 92 | 95 | 38 | 29 | 28 |
| TMEM263 | 71 | 59 | 52 | 0 | 14 | 0 |
| TNR | 100 | 95 | 95 | 24 | 32 | 31 |
| TP53BP2 | 75 | 80 | 80 | 24 | 9 | 35 |
| TRIM16 | 100 | 100 | 100 | -1 | -1 | -1 |
| TSPAN11 | 100 | 94 | 97 | 19 | 16 | 20 |
| TTBK2 | -1 | 100 | 100 | 25 | 33 | 57 |
| VARS2 | 98 | 98 | 99 | 20 | 14 | 19 |
| WDR41 | 88 | 89 | 93 | 4 | 4 | 19 |
| WRAP73 | 100 | 100 | 100 | 38 | 29 | 38 |
| WSCD1 | 100 | 100 | 100 | 45 | 28 | 34 |
| XPNPEP1 | 100 | 99 | 100 | 24 | 23 | 26 |
| ZMYND11 | 78 | 64 | 63 | 2 | 6 | 12 |
| ZNF767P | 82 | 88 | 85 | 21 | 20 | 8 |

C

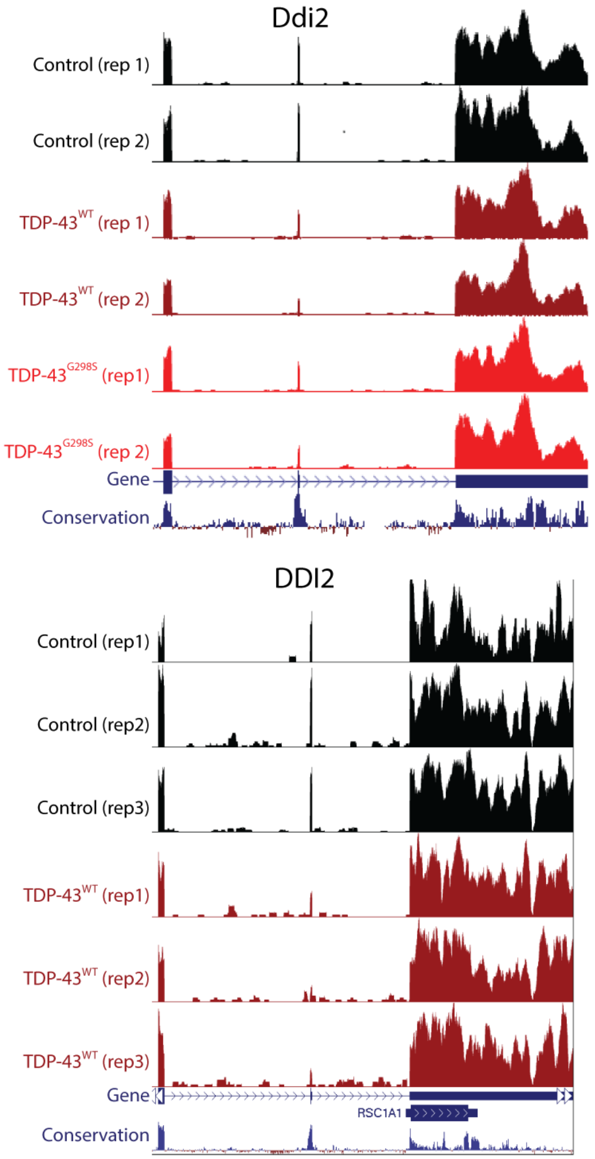

**Supplementary Figure 3. Differences in exon skipping targets between mouse and human.** (**A**) We conducted a comparative analysis of RNA-seq datasets obtained from transgenic lines that overexpressed either TDP-43$^{G298S}$ or TDP-43$^{WT}$ and control littermates. Our analysis revealed that both transgenic lines showed similar profiles of exon repression when compared to the control group. (**B**) Moreover, the comparative analysis of RNA-seq data from i3N human neurons revealed that the set of repressed exons observed in mice differed from those observed in humans. (**C**) Notably, only one target, *Ddi2*, was found to have a repressed exon by TDP-43 overexpression in both mouse and human samples, as shown in the UCSC genome browser plots (Supplemental Data file).

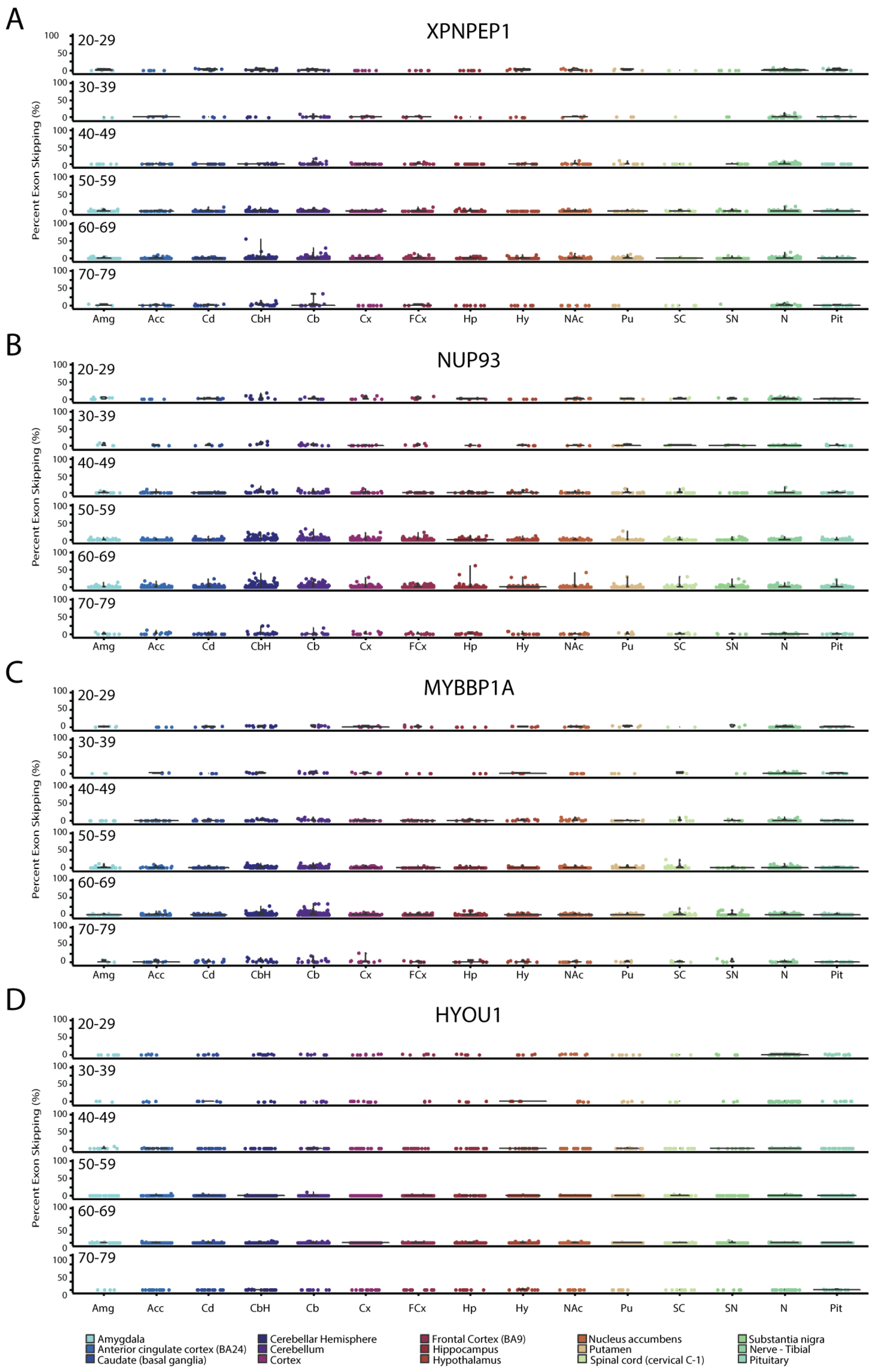
A
XPNPEP1
B
NUP93
C
MYBBP1A
D
HYOU1
Percent Exon Skipping (%)
20-29
30-39
40-49
50-59
60-69
70-79
100
50
0
Amg Acc Cd CbH Cb Cx FCx Hp Hy NAc Pu SC SN N Pit
Amygdala
Anterior cingulate cortex (BA24)
Caudate (basal ganglia)
Cerebellar Hemisphere
Cerebellum
Cortex
Frontal Cortex (BA9)
Hippocampus
Hypothalamus
Nucleus accumbens
Putamen
Spinal cord (cervical C-1)
Substantia nigra
Nerve - Tibial
Pituitary

**Supplementary Figure 4. Exon skipping events are found in human brain samples**. Aging is the main risk factor for developing neurodegenerative diseases. We show that skipping events appear normally in the CNS, but we wanted to explore whether skipping events correlate with aging. We queried the splice junction archive, Snaptron, and analyzed publicly available human RNA-seq datasets from GTEx. We show the percent of exon skipping in four genes at ages ranging from 20-29, 30-39, 40-49, 50-59, 60-69, and 70-79. Skipping events were found in most of the different brain areas analyzed, with a higher frequency in the cerebellum and cortex.
