## Supplementary Data for "Elevated nuclear TDP-43 induces constitutive exon skipping"

A

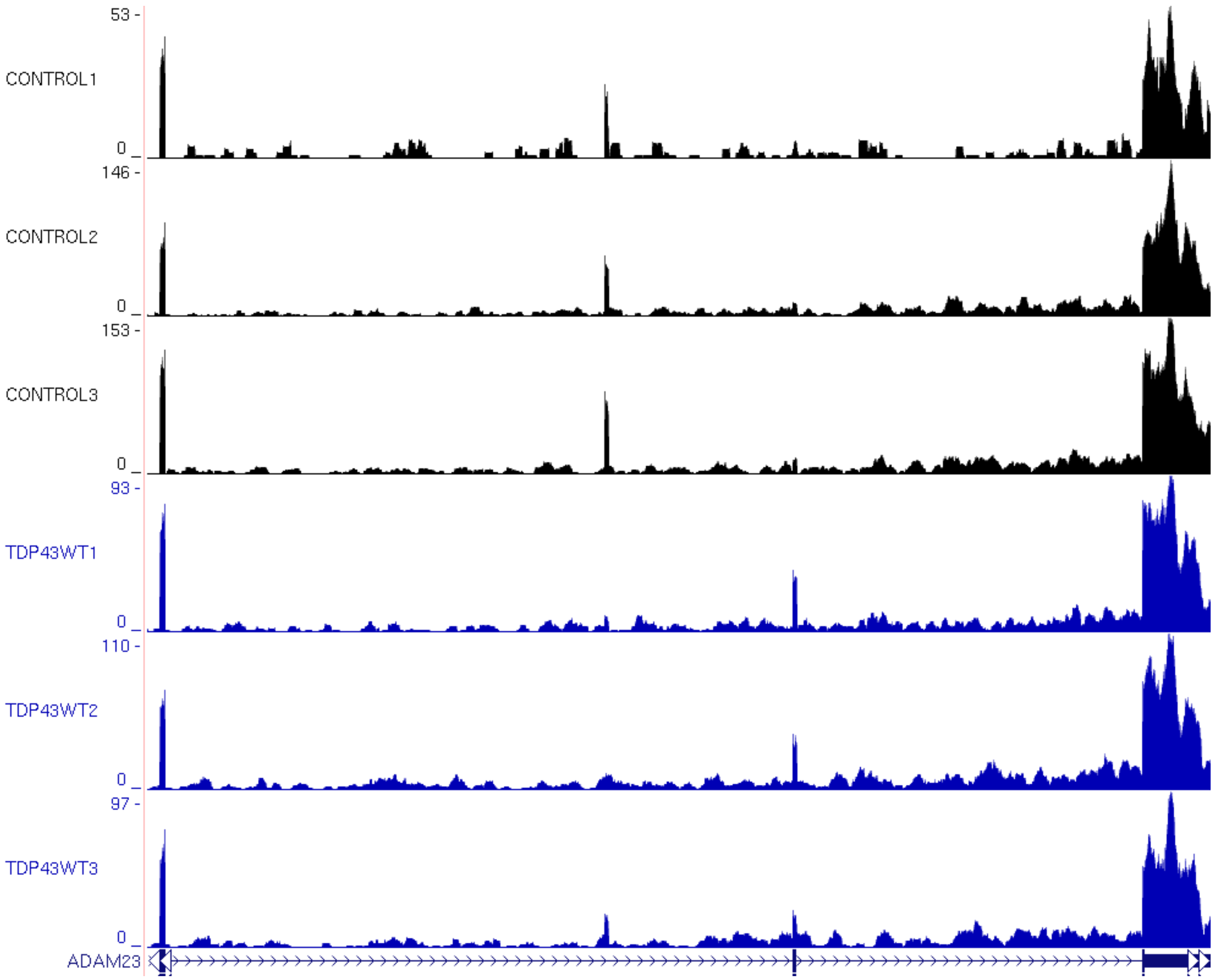

53 -
CONTROL1
0
146 -
CONTROL2
0
153 -
CONTROL3
0
93 -
TDP43WT1
0
110 -
TDP43WT2
0
97 -
TDP43WT3
0
ADAM23

B

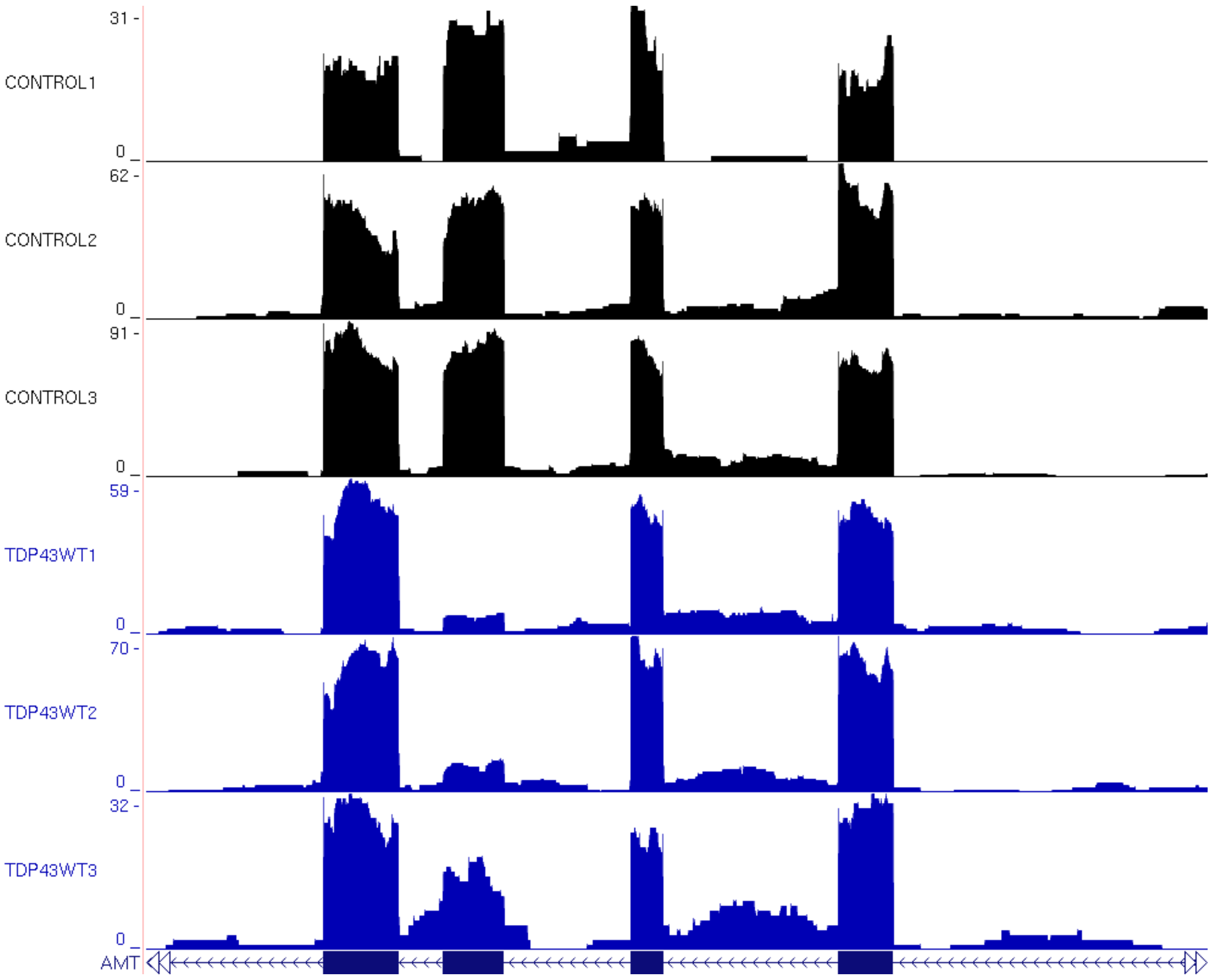
31 -
CONTROL1
0 _
62 -
CONTROL2
0 _
91 -
CONTROL3
0 _
59 -
TDP43WT1
0 _
70 -
TDP43WT2
0 _
32 -
TDP43WT3
0 _
AMT

C

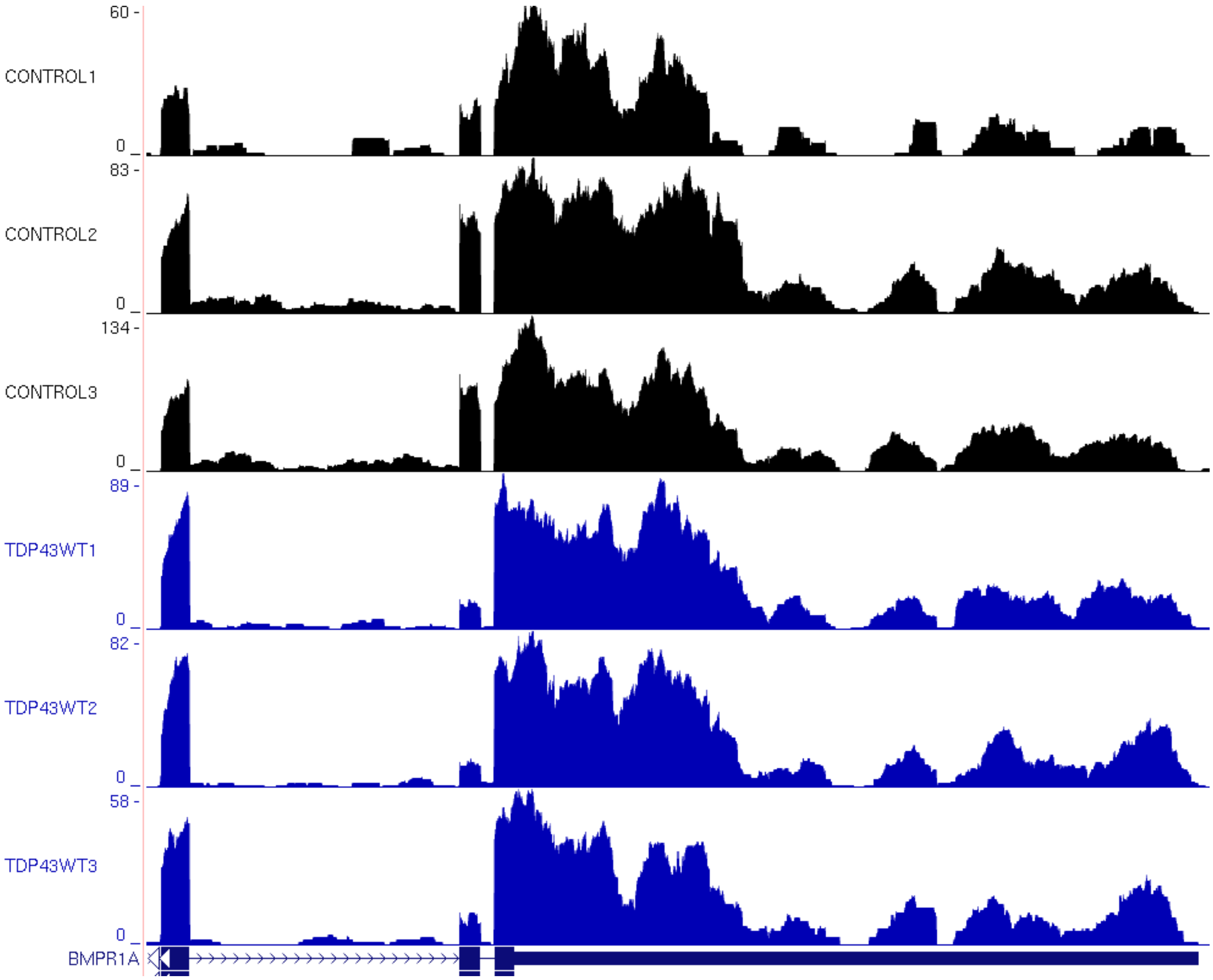
60 -
CONTROL1
0
83 -
CONTROL2
0
134 -
CONTROL3
0
89 -
TDP43WT1
0
82 -
TDP43WT2
0
58 -
TDP43WT3
0
BMPR1A

D

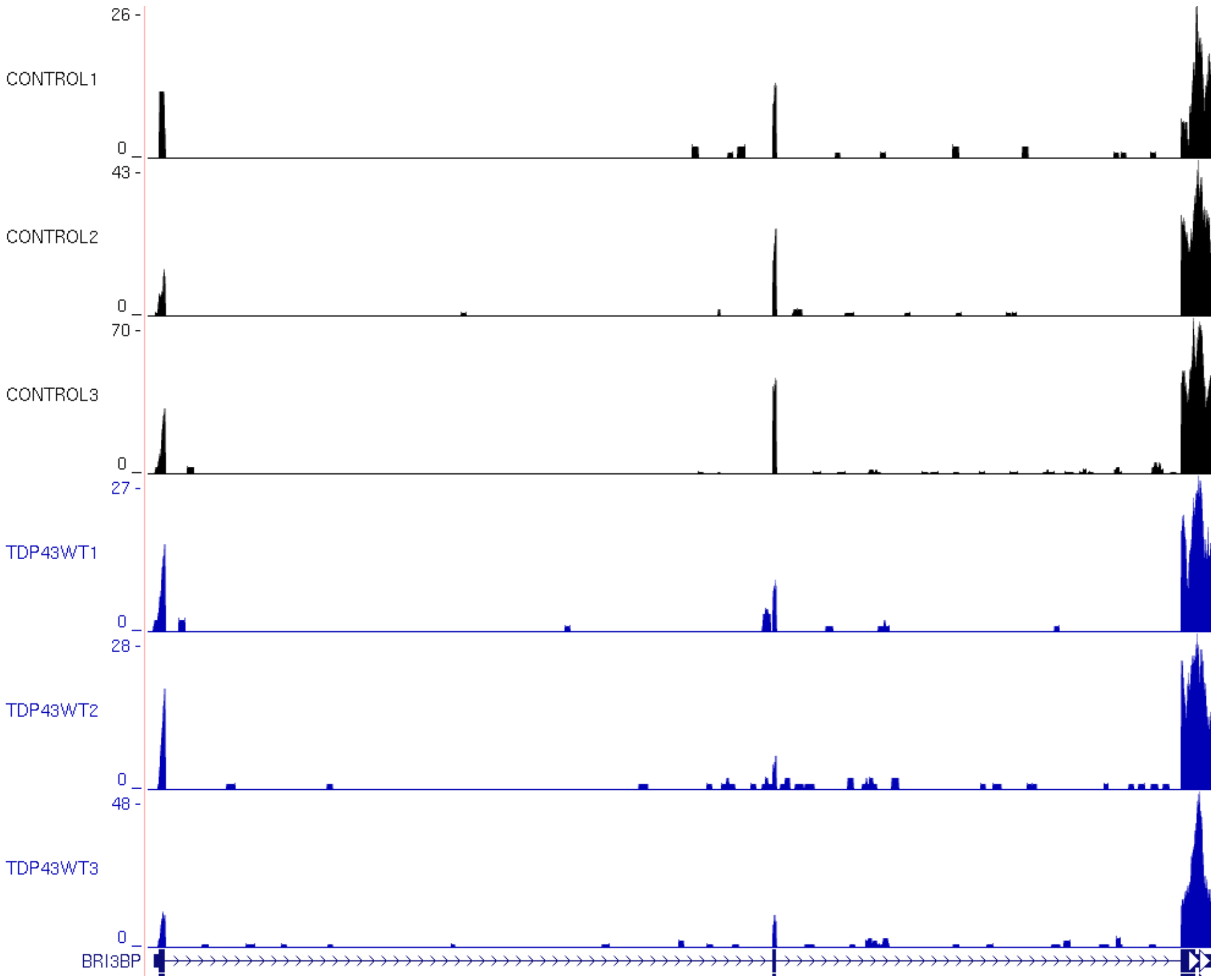
26 -
CONTROL1
0
43 -
CONTROL2
0
70 -
CONTROL3
0
27 -
TDP43WT1
0
28 -
TDP43WT2
0
48 -
TDP43WT3
0
BRI3BP

E

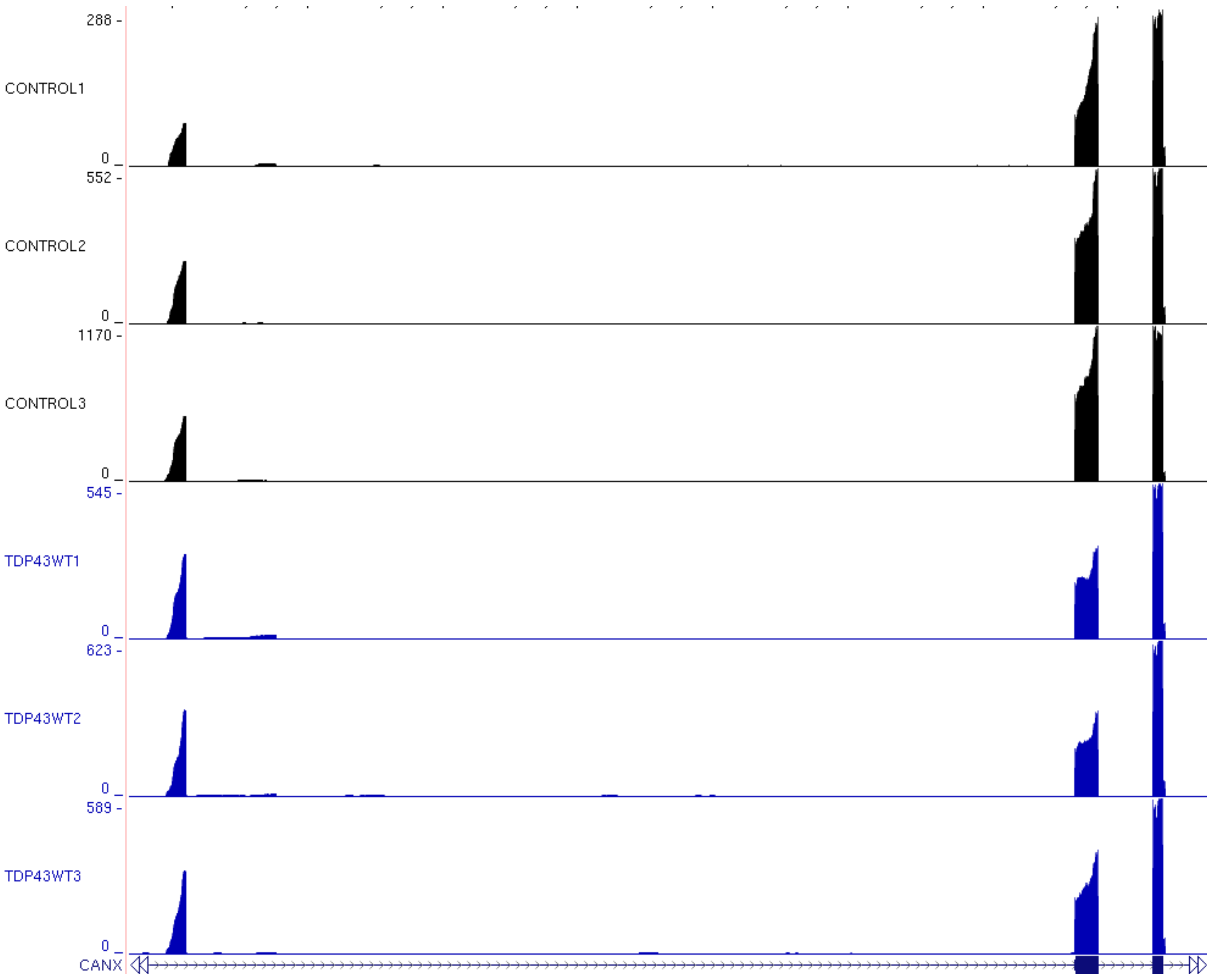
288 -
CONTROL1
0 _
552 -
CONTROL2
0 _
1170 -
CONTROL3
0 _
545 -
TDP43WT1
0 _
623 -
TDP43WT2
0 _
589 -
TDP43WT3
0 _
CANX

F

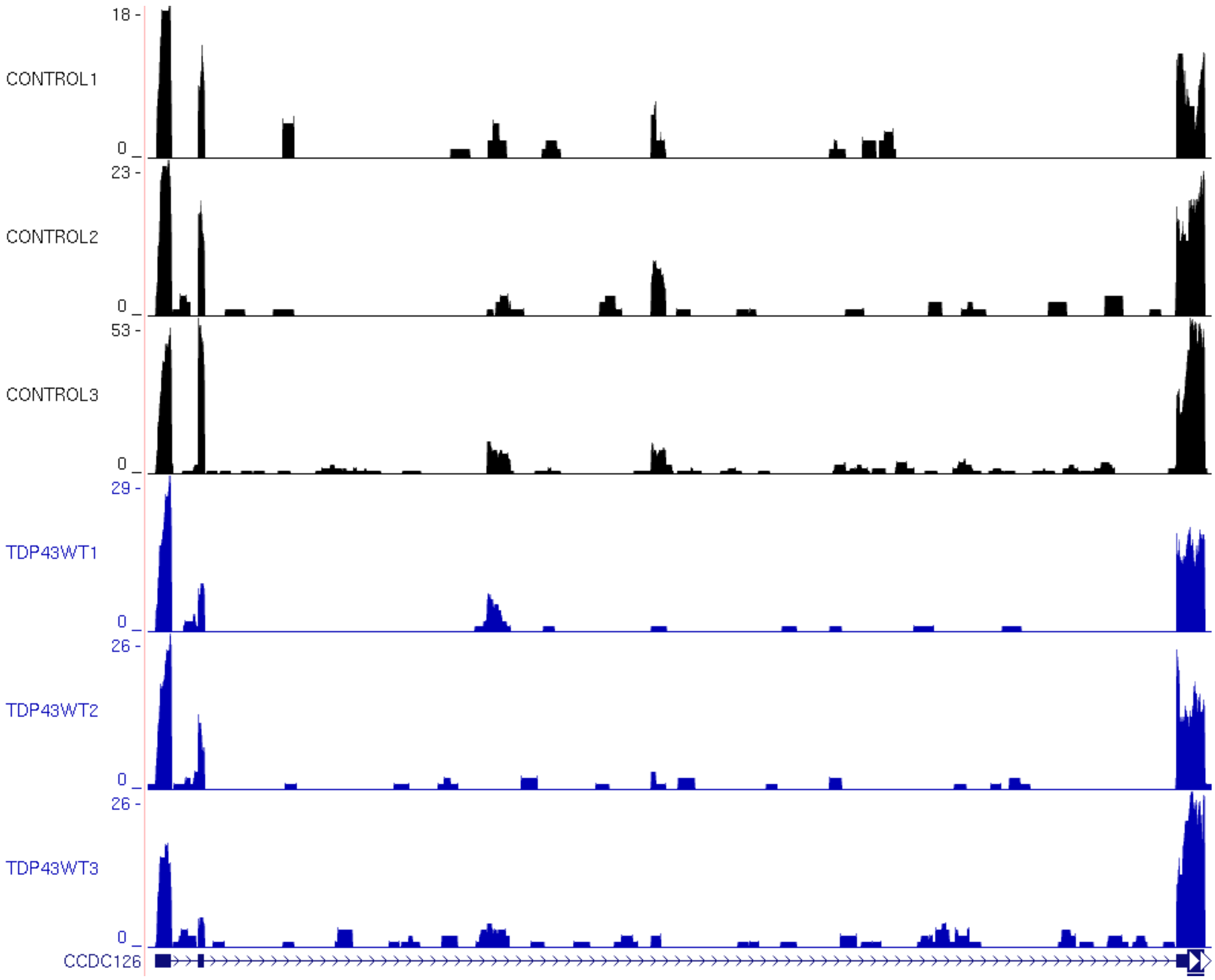
18 -
CONTROL1
0
23 -
CONTROL2
0
53 -
CONTROL3
0
29 -
TDP43WT1
0
26 -
TDP43WT2
0
26 -
TDP43WT3
0
CCDC126

G

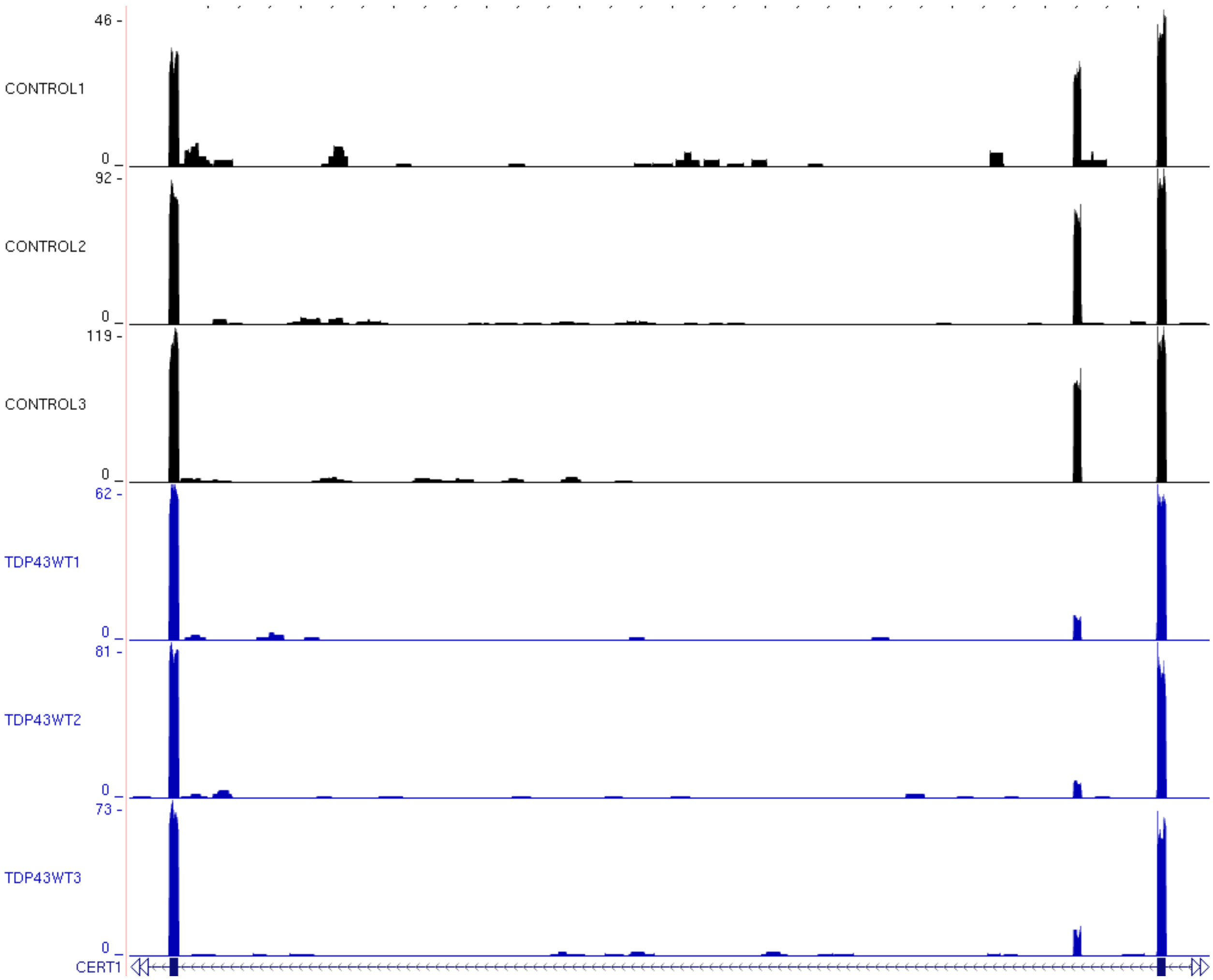

46 -
CONTROL1
0 _
92 -
CONTROL2
0 _
119 -
CONTROL3
0 _
62 -
TDP43WT1
0 _
81 -
TDP43WT2
0 _
73 -
TDP43WT3
0 _
CERT1

H

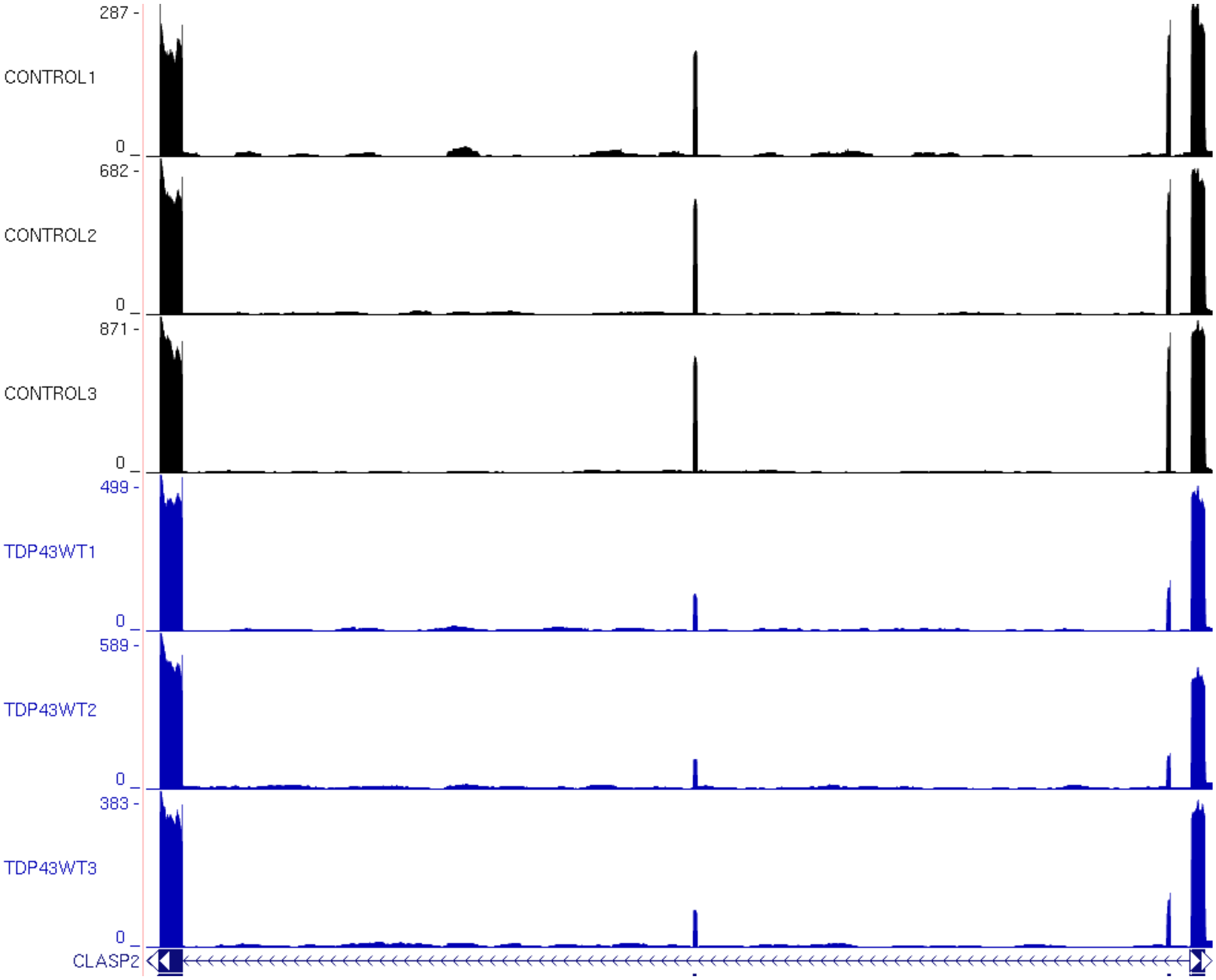

287 -
CONTROL1
0 _
682 -
CONTROL2
0 _
871 -
CONTROL3
0 _
499 -
TDP43WT1
0 _
589 -
TDP43WT2
0 _
383 -
TDP43WT3
0 _
CLASP2

I

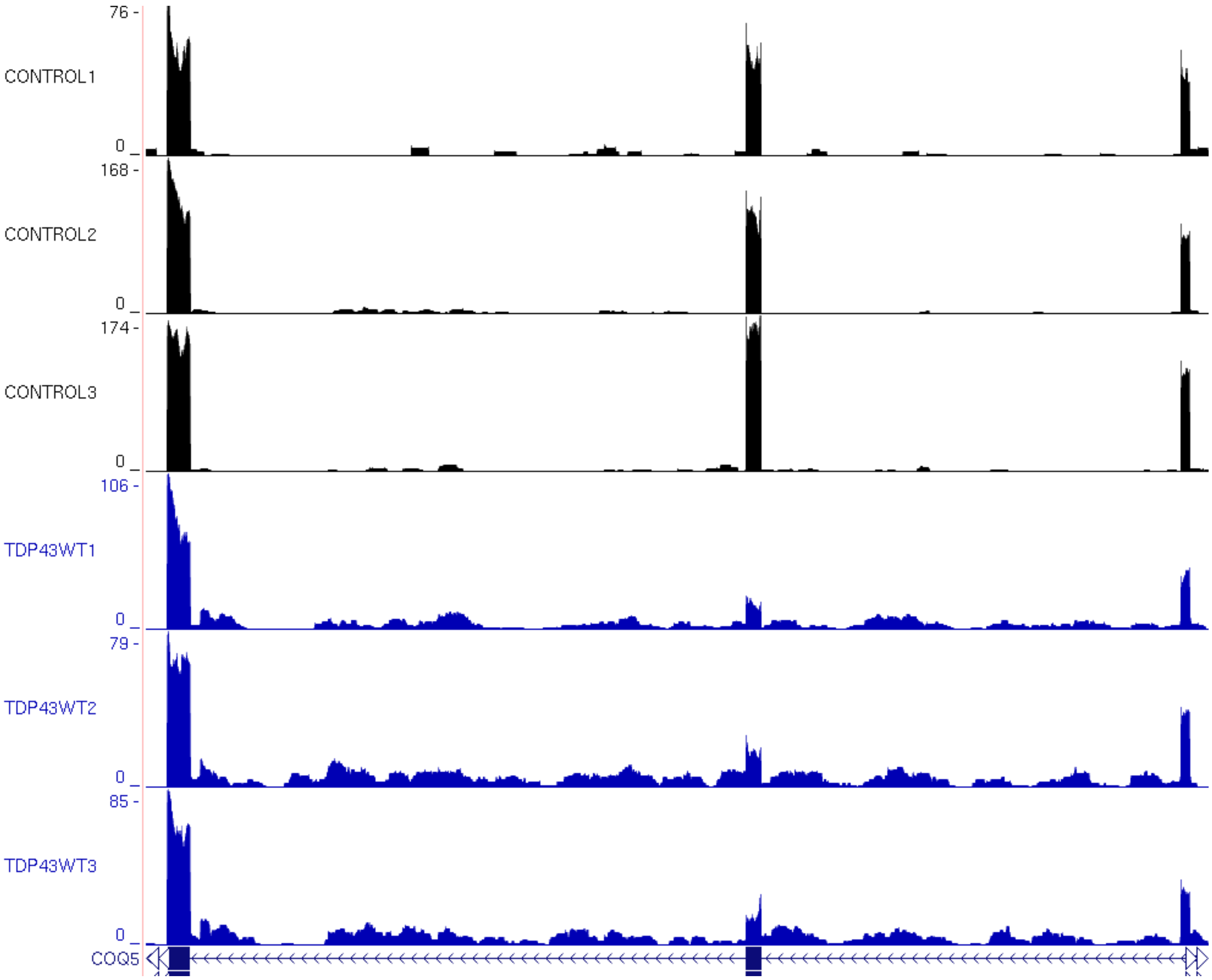

76
0
CONTROL1
168
0
CONTROL2
174
0
CONTROL3
106
0
TDP43WT1
79
0
TDP43WT2
85
0
TDP43WT3
COQ5

J

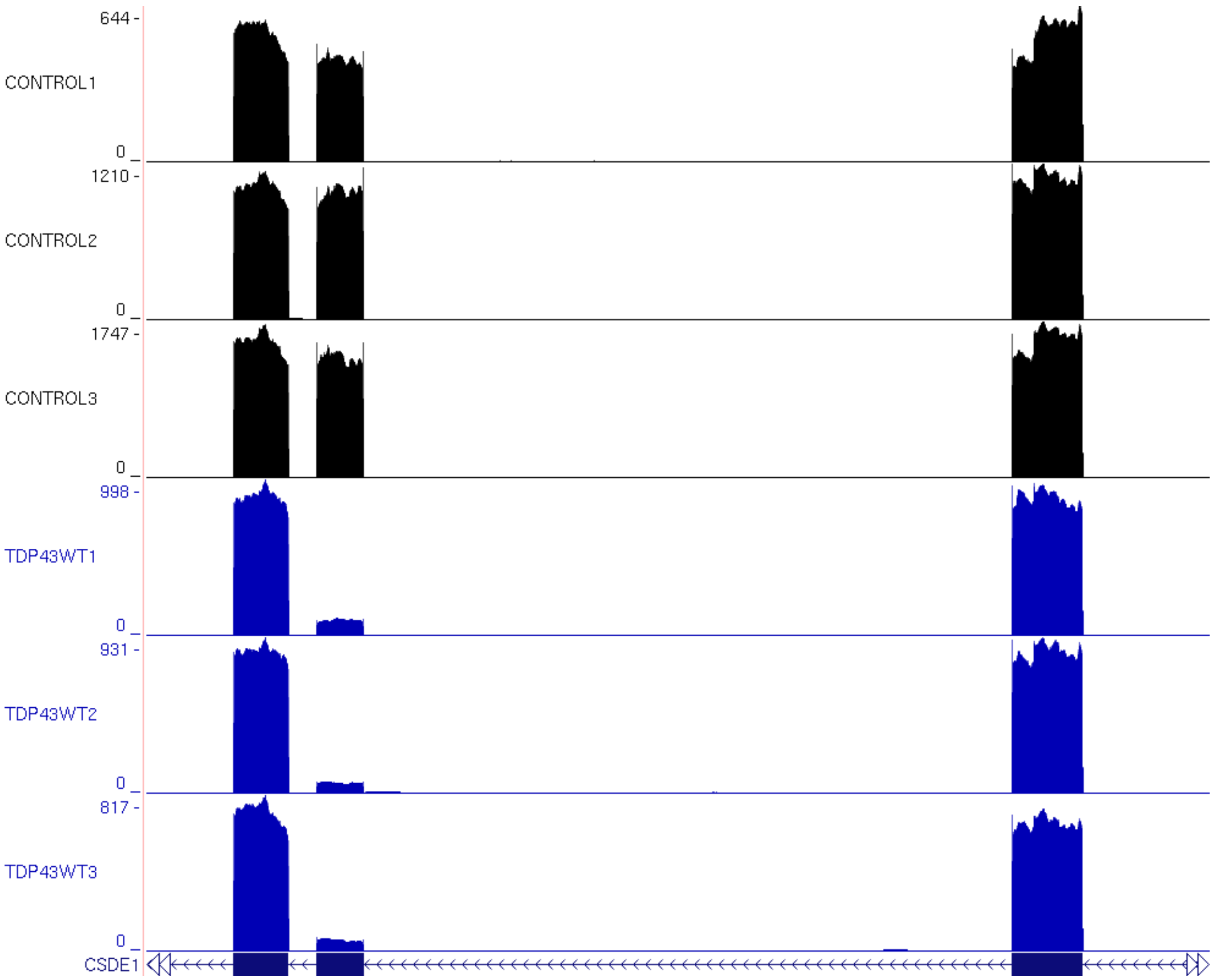
644 -
CONTROL1
0 _
1210 -
CONTROL2
0 _
1747 -
CONTROL3
0 _
998 -
TDP43WT1
0 _
931 -
TDP43WT2
0 _
817 -
TDP43WT3
0 _
CSDE1

K

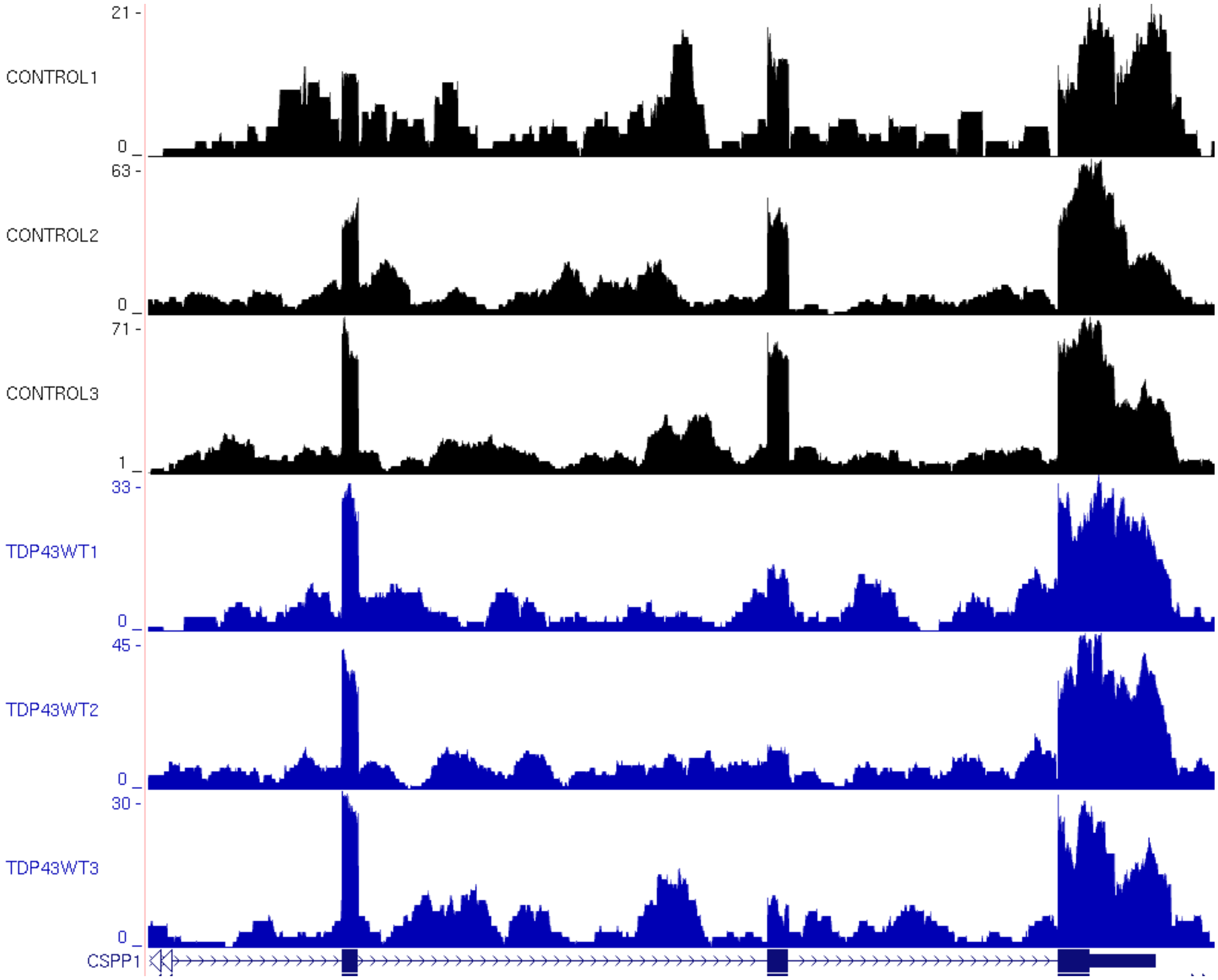

21 -
CONTROL1
0 _
63 -
CONTROL2
0 _
71 -
CONTROL3
1 _
33 -
TDP43WT1
0 _
45 -
TDP43WT2
0 _
30 -
TDP43WT3
0 _
CSPP1

L

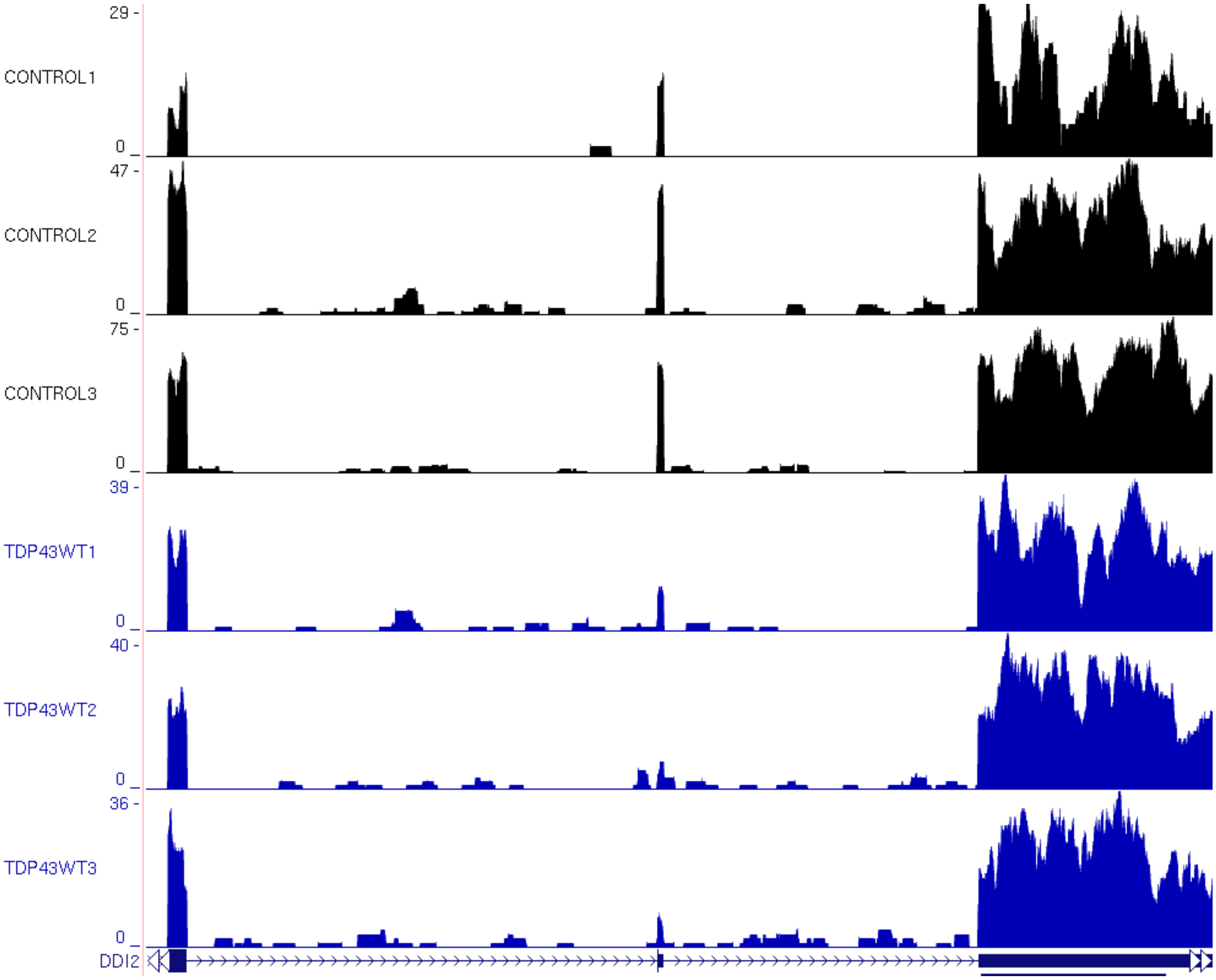
29 -
CONTROL1
0
47 -
CONTROL2
0
75 -
CONTROL3
0
39 -
TDP43WT1
0
40 -
TDP43WT2
0
36 -
TDP43WT3
0
DDI2

M

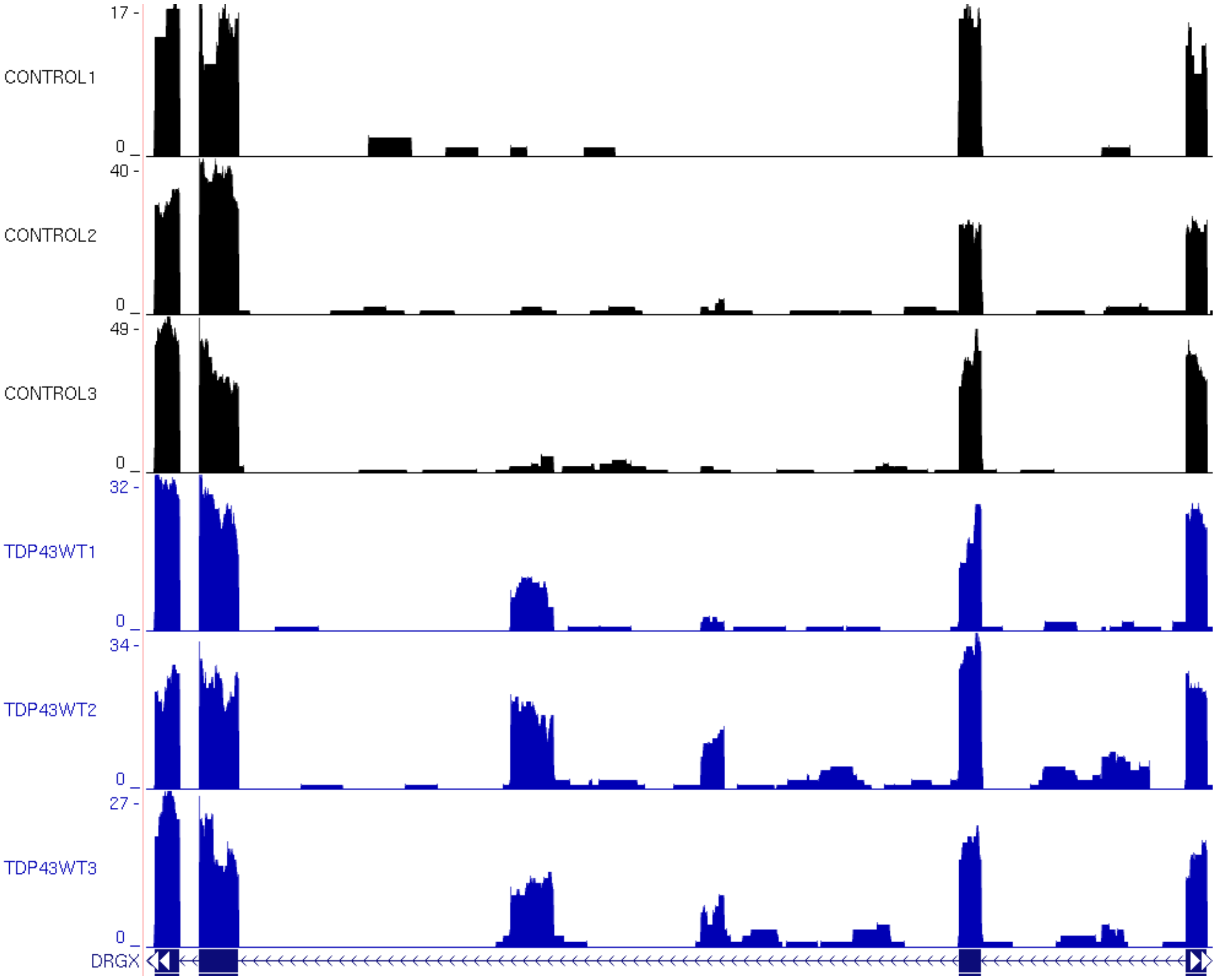

17 -
CONTROL1
0
40 -
CONTROL2
0
49 -
CONTROL3
0
32 -
TDP43WT1
0
34 -
TDP43WT2
0
27 -
TDP43WT3
0
DRGX

N

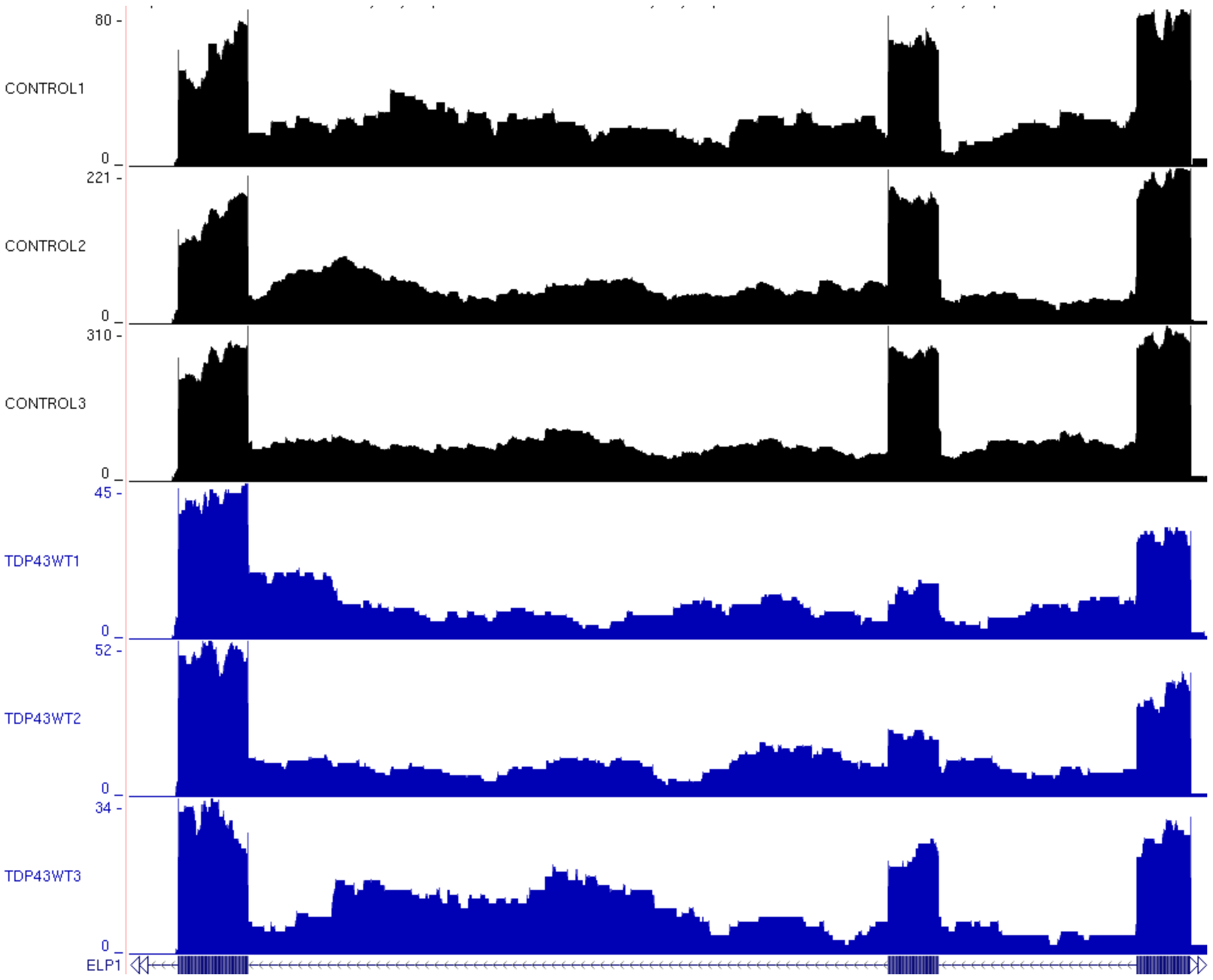
80
CONTROL1
0
221
CONTROL2
0
310
CONTROL3
0
45
TDP43WT1
0
52
TDP43WT2
0
34
TDP43WT3
0
ELP1

O

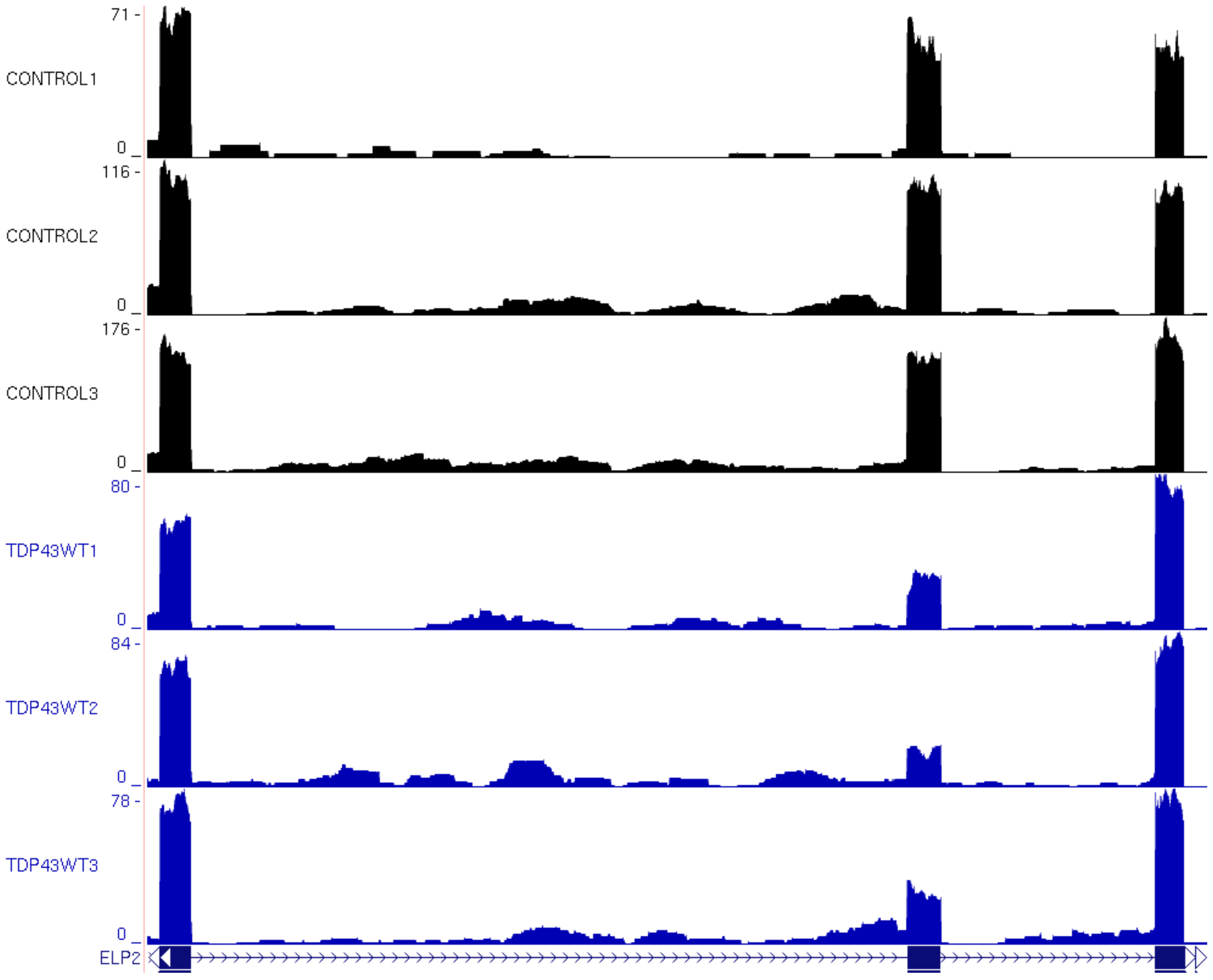
71 -
CONTROL1
0
116 -
CONTROL2
0
176 -
CONTROL3
0
80 -
TDP43WT1
0
84 -
TDP43WT2
0
78 -
TDP43WT3
0
ELP2

P

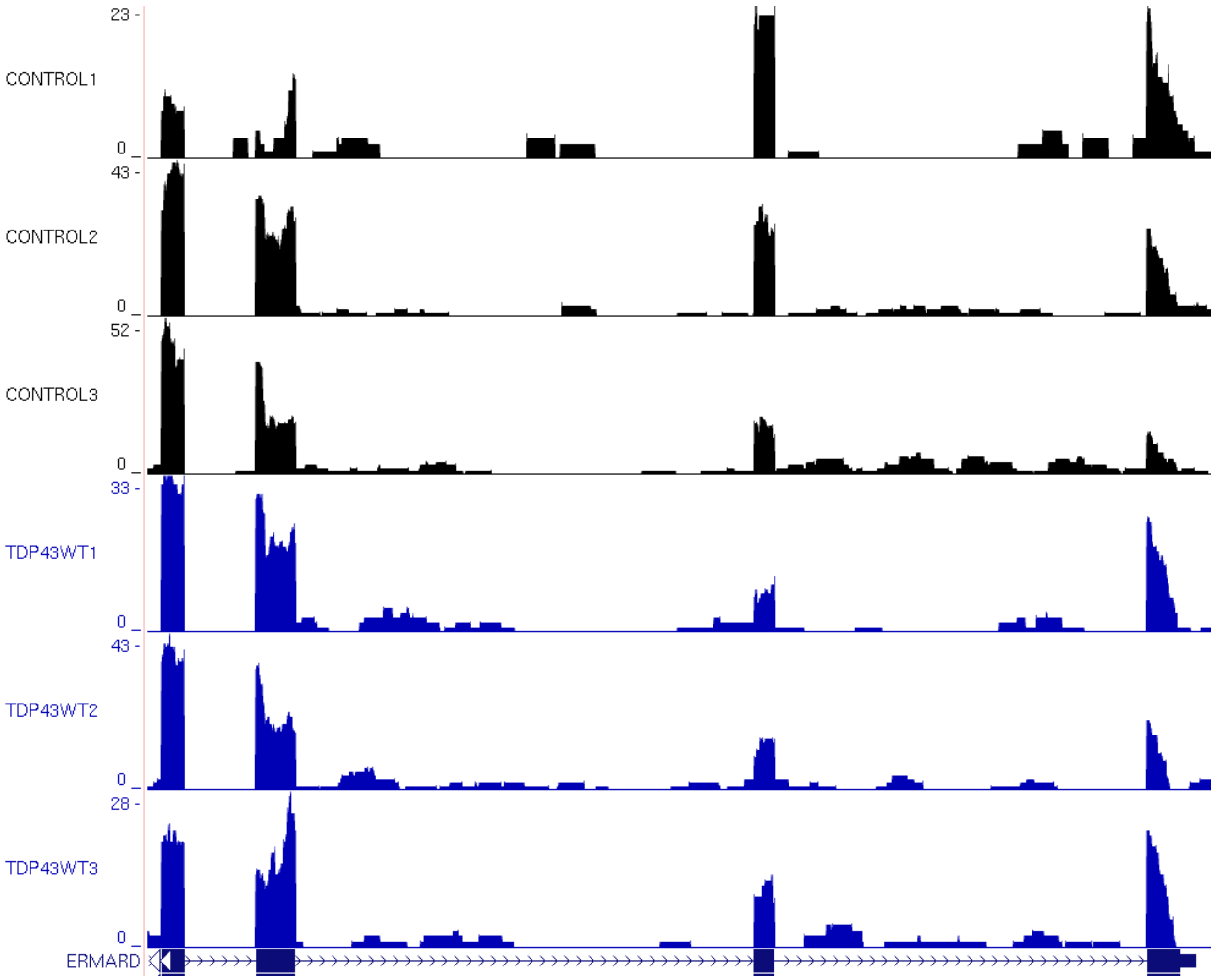
23 -
CONTROL1
0
43 -
CONTROL2
0
52 -
CONTROL3
0
33 -
TDP43WT1
0
43 -
TDP43WT2
0
28 -
TDP43WT3
0
ERMARD

Q

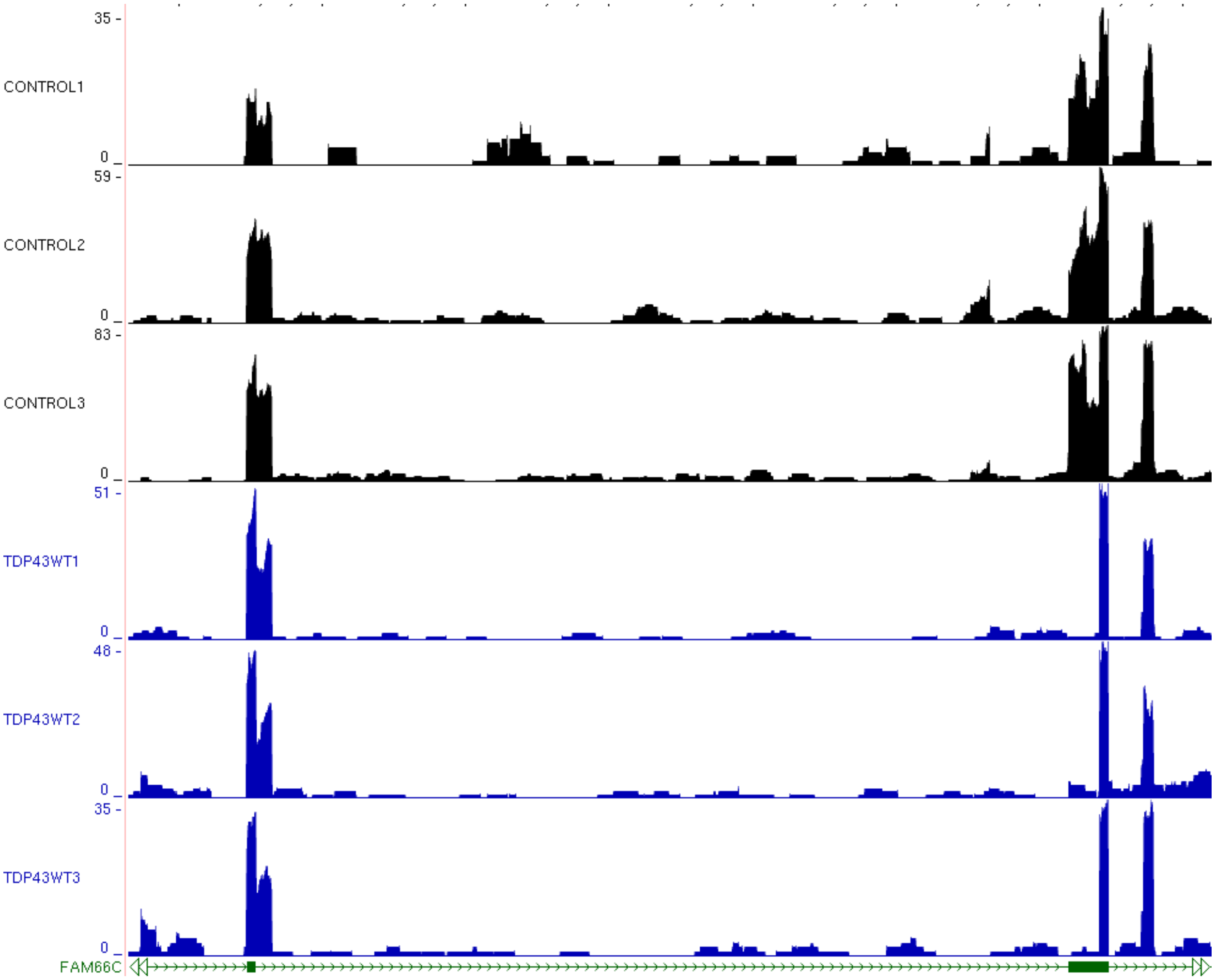
35 -
CONTROL1
0
59 -
CONTROL2
0
83 -
CONTROL3
0
51 -
TDP43WT1
0
48 -
TDP43WT2
0
35 -
TDP43WT3
0
FAM66C

R

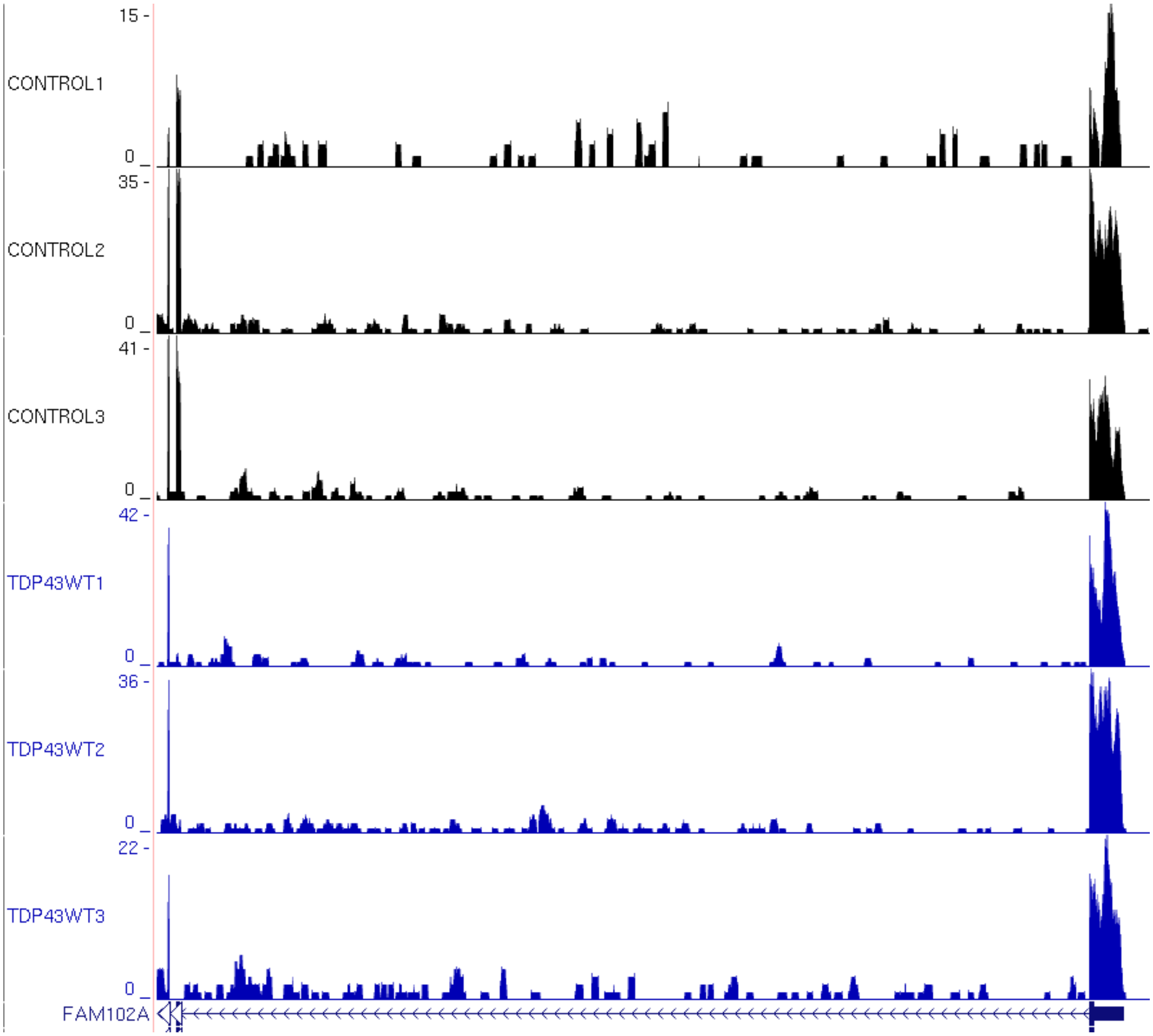
15 -
CONTROL1
0
35 -
CONTROL2
0
41 -
CONTROL3
0
42 -
TDP43WT1
0
36 -
TDP43WT2
0
22 -
TDP43WT3
0
FAM102A

S

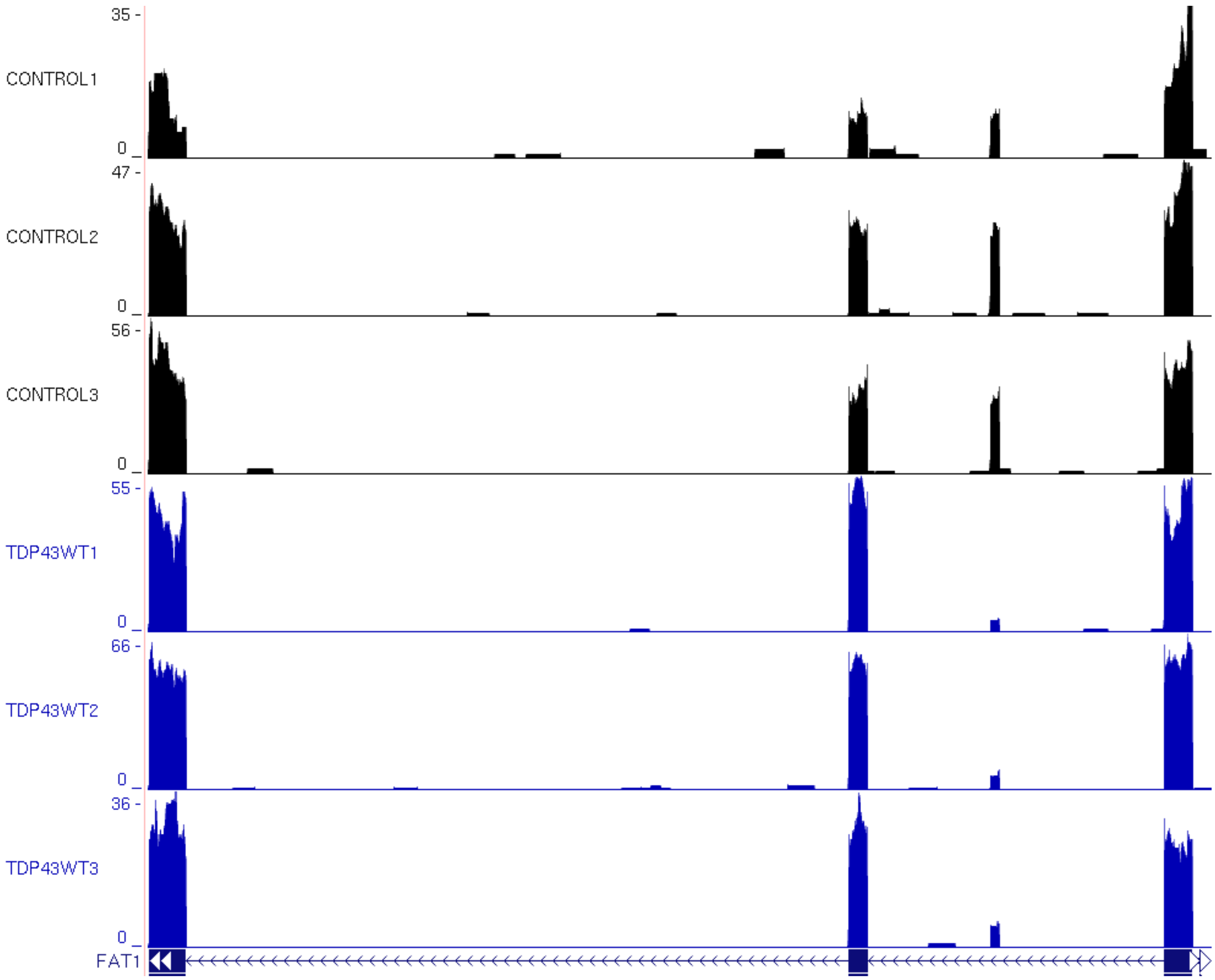
35 -
CONTROL1
0
47 -
CONTROL2
0
56 -
CONTROL3
0
55 -
TDP43WT1
0
66 -
TDP43WT2
0
36 -
TDP43WT3
0
FAT1

T

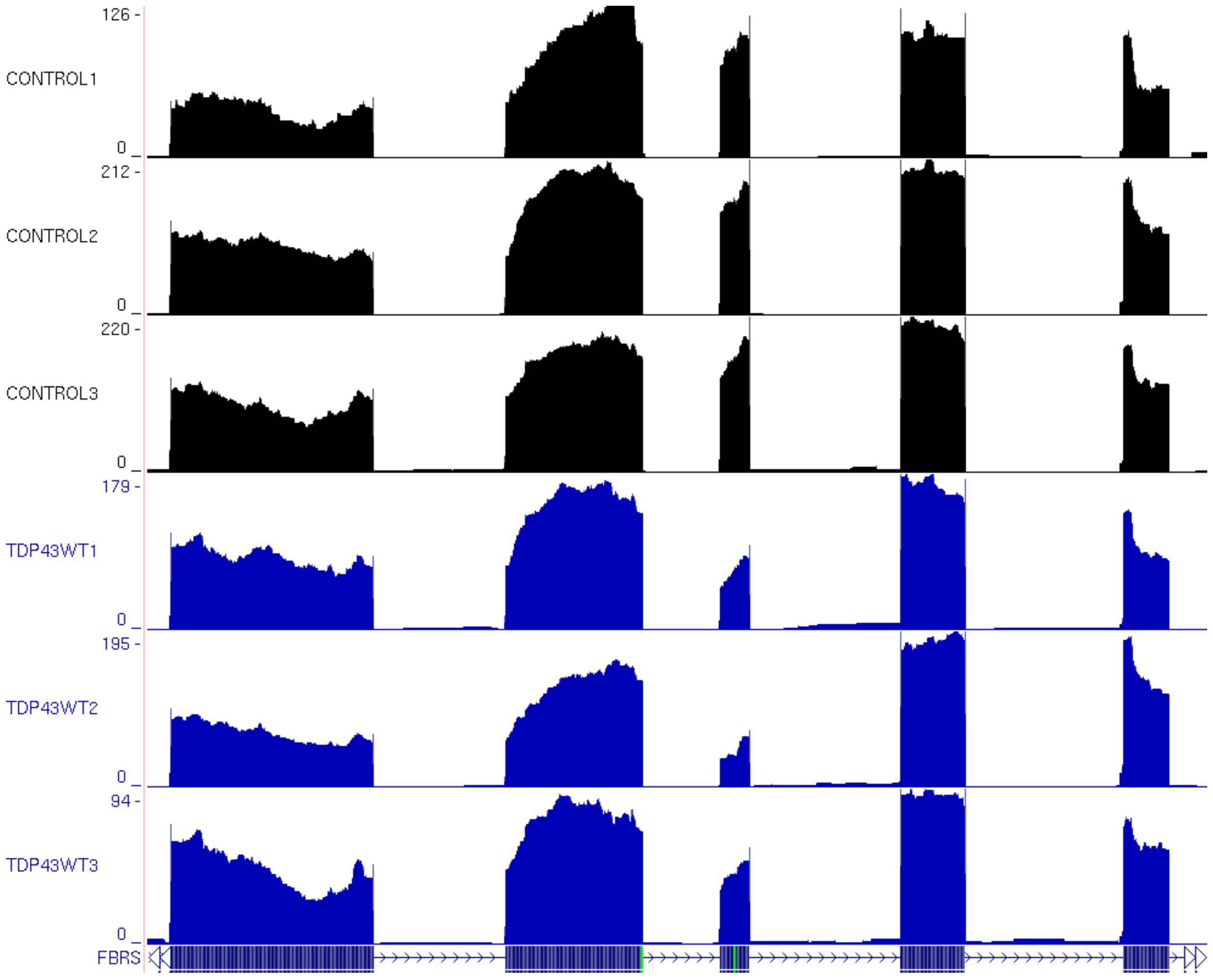
126 -
CONTROL1
0
212 -
CONTROL2
0
220 -
CONTROL3
0
179 -
TDP43WT1
0
195 -
TDP43WT2
0
94 -
TDP43WT3
0
FBRS

U

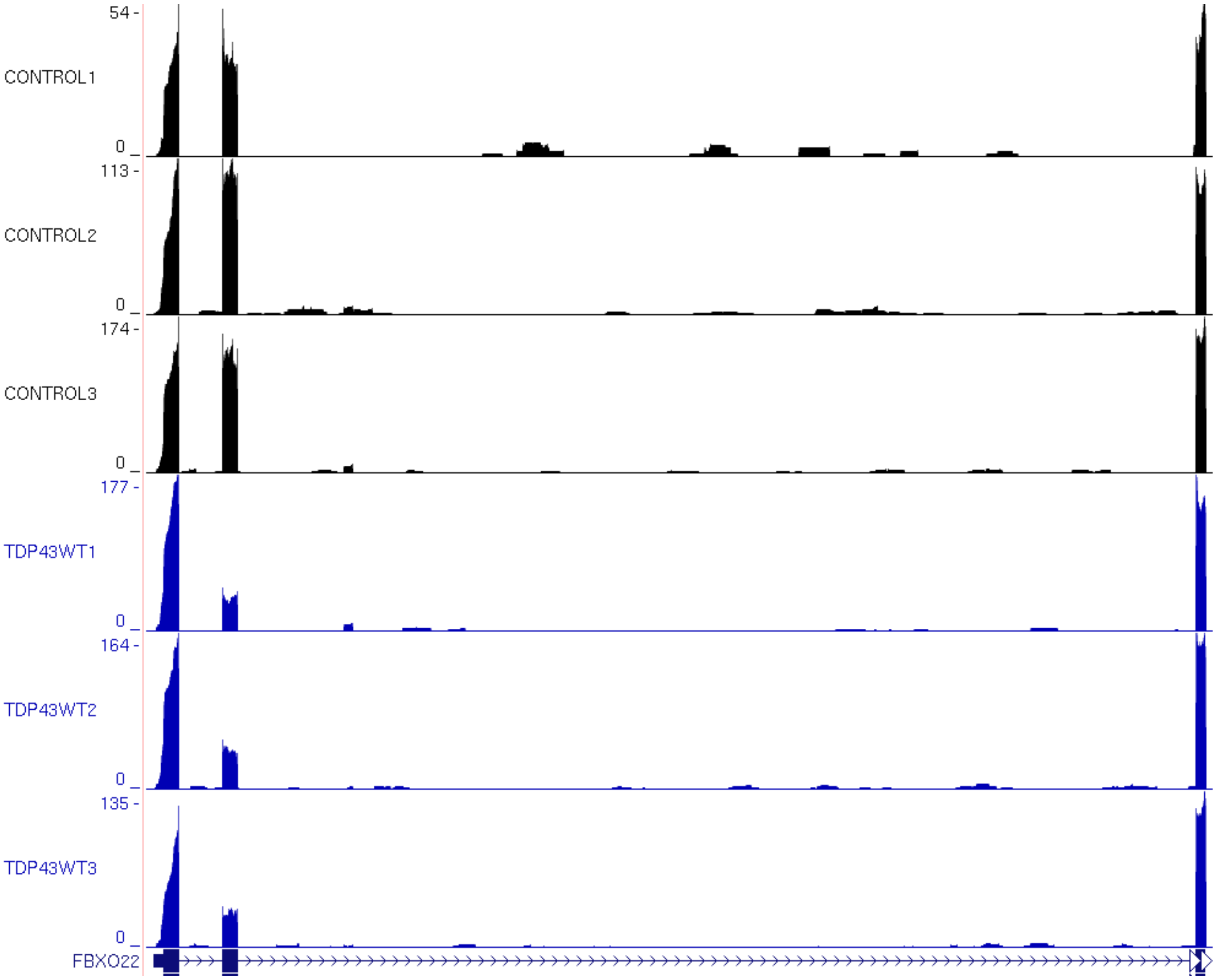
54 -
CONTROL1
0
113 -
CONTROL2
0
174 -
CONTROL3
0
177 -
TDP43WT1
0
164 -
TDP43WT2
0
135 -
TDP43WT3
0
FBXO22

V

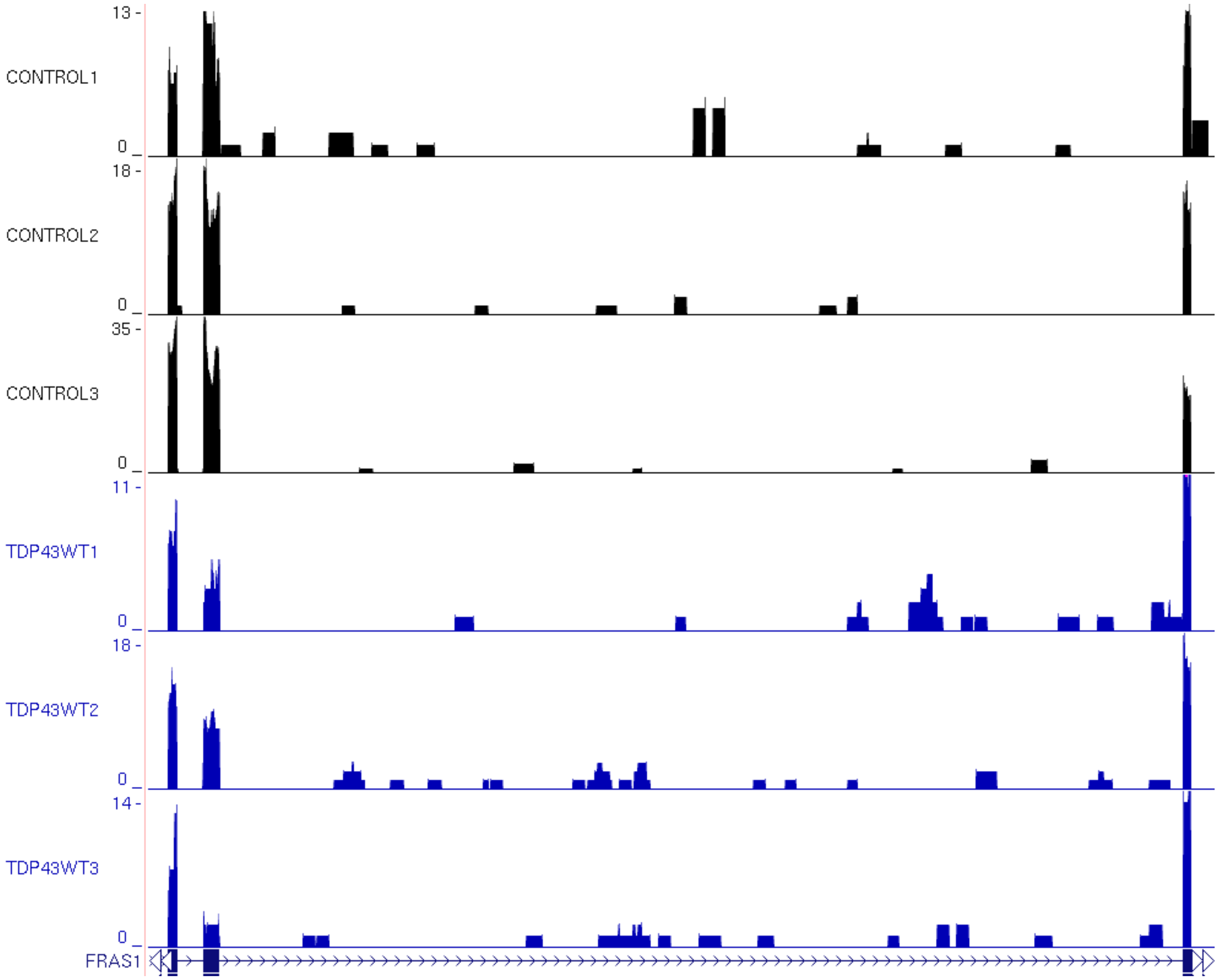
13 -
CONTROL1
0
18 -
CONTROL2
0
35 -
CONTROL3
0
11 -
TDP43WT1
0
18 -
TDP43WT2
0
14 -
TDP43WT3
0
FRAS1

W

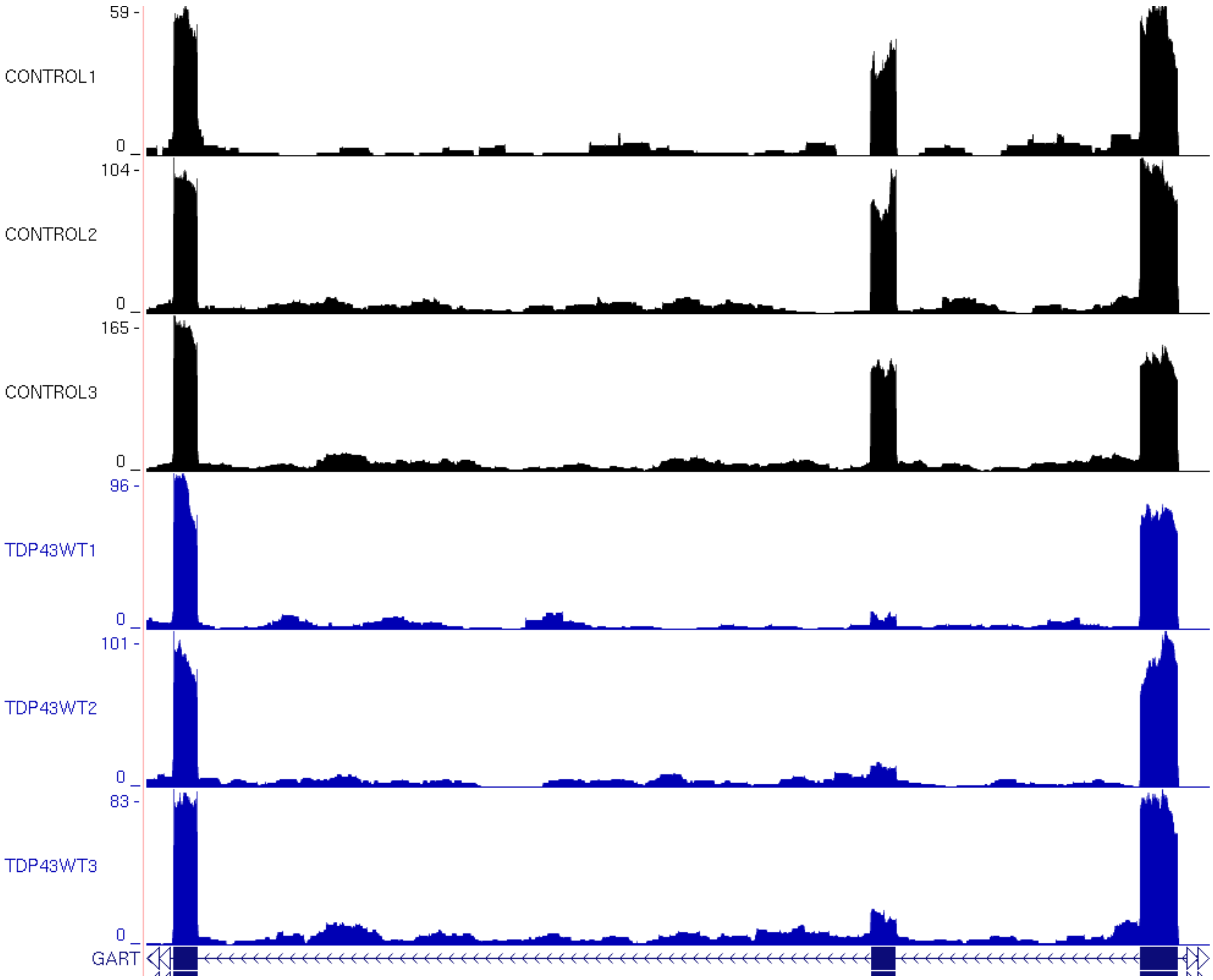

59 -
CONTROL1
0
104 -
CONTROL2
0
165 -
CONTROL3
0
96 -
TDP43WT1
0
101 -
TDP43WT2
0
83 -
TDP43WT3
0
GART

X

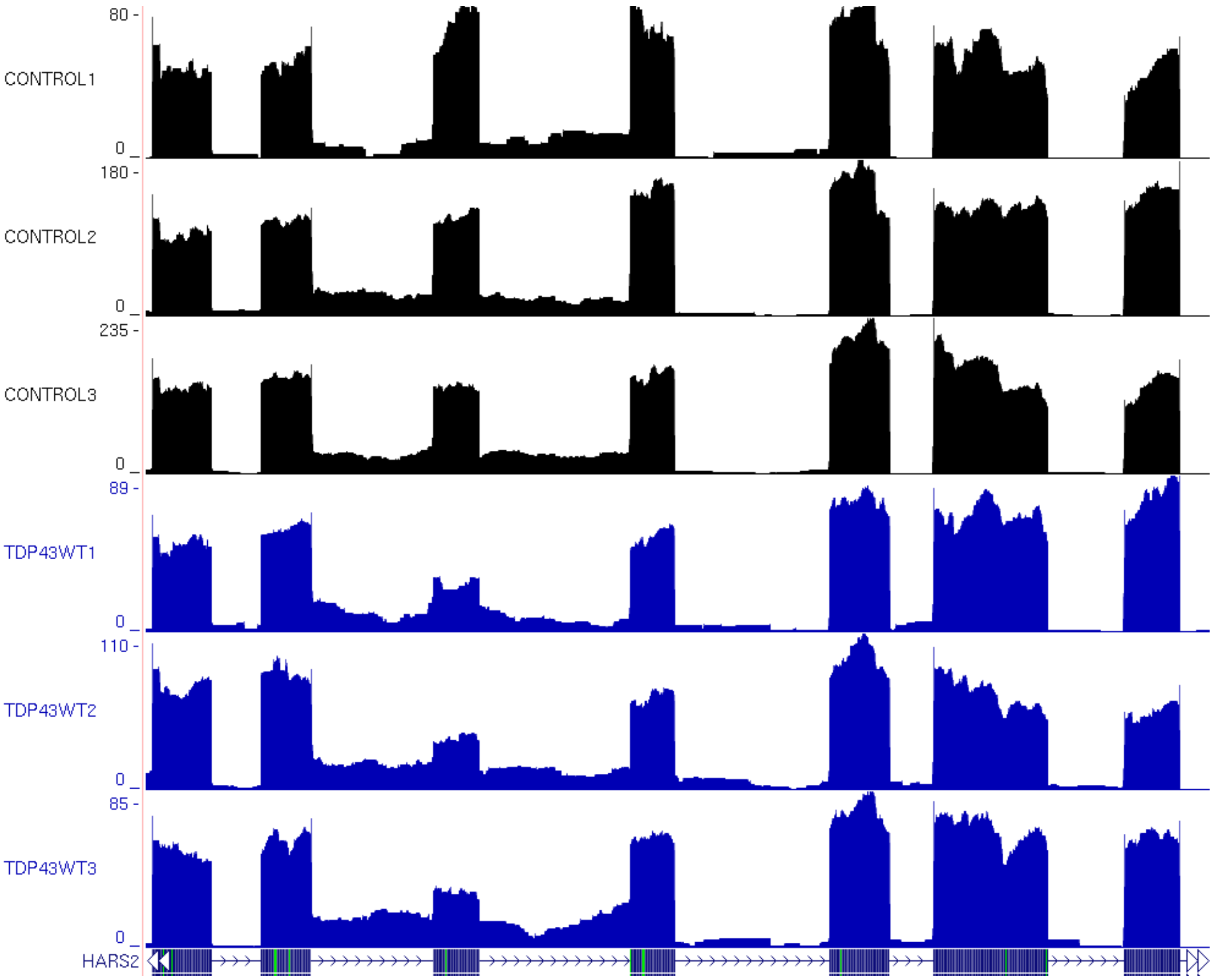
80 -
CONTROL1
0
180 -
CONTROL2
0
235 -
CONTROL3
0
89 -
TDP43WT1
0
110 -
TDP43WT2
0
85 -
TDP43WT3
0
HARS2

Y

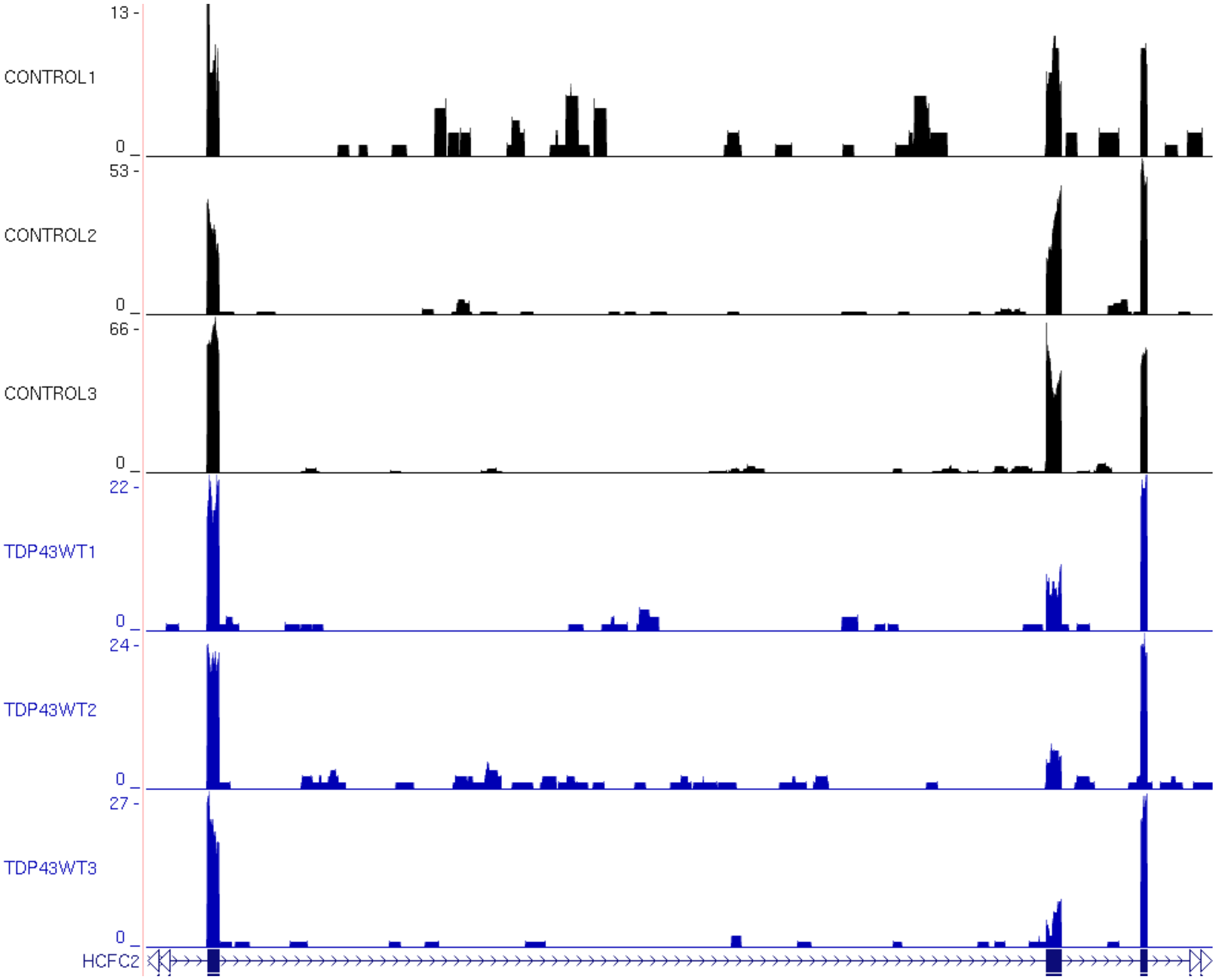

13 -
CONTROL1
0
53 -
CONTROL2
0
66 -
CONTROL3
0
22 -
TDP43WT1
0
24 -
TDP43WT2
0
27 -
TDP43WT3
0
HCFC2

Z

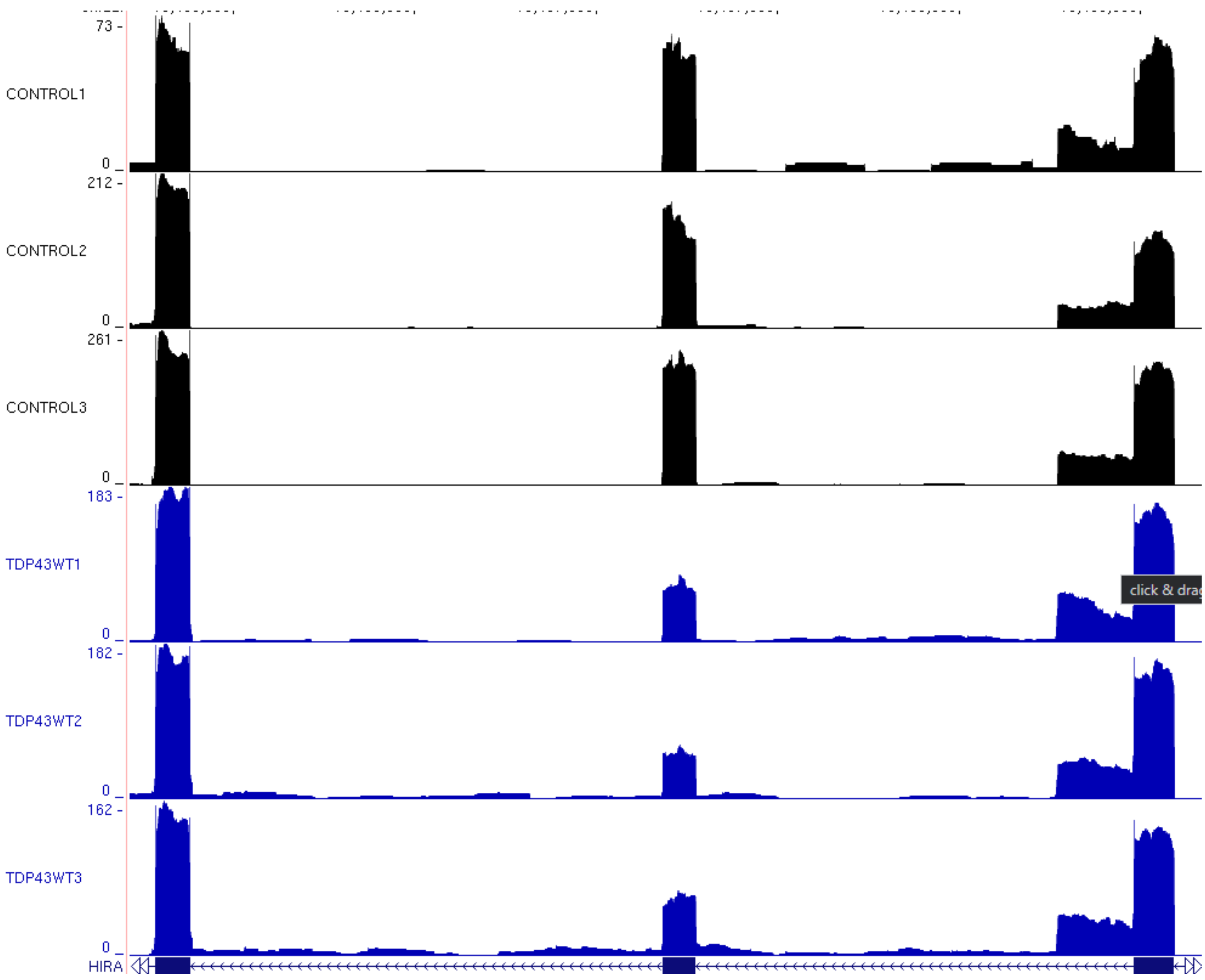
73
CONTROL1
0
212
CONTROL2
0
261
CONTROL3
0
183
TDP43WT1
0
182
TDP43WT2
0
162
TDP43WT3
0
HIRA

AA

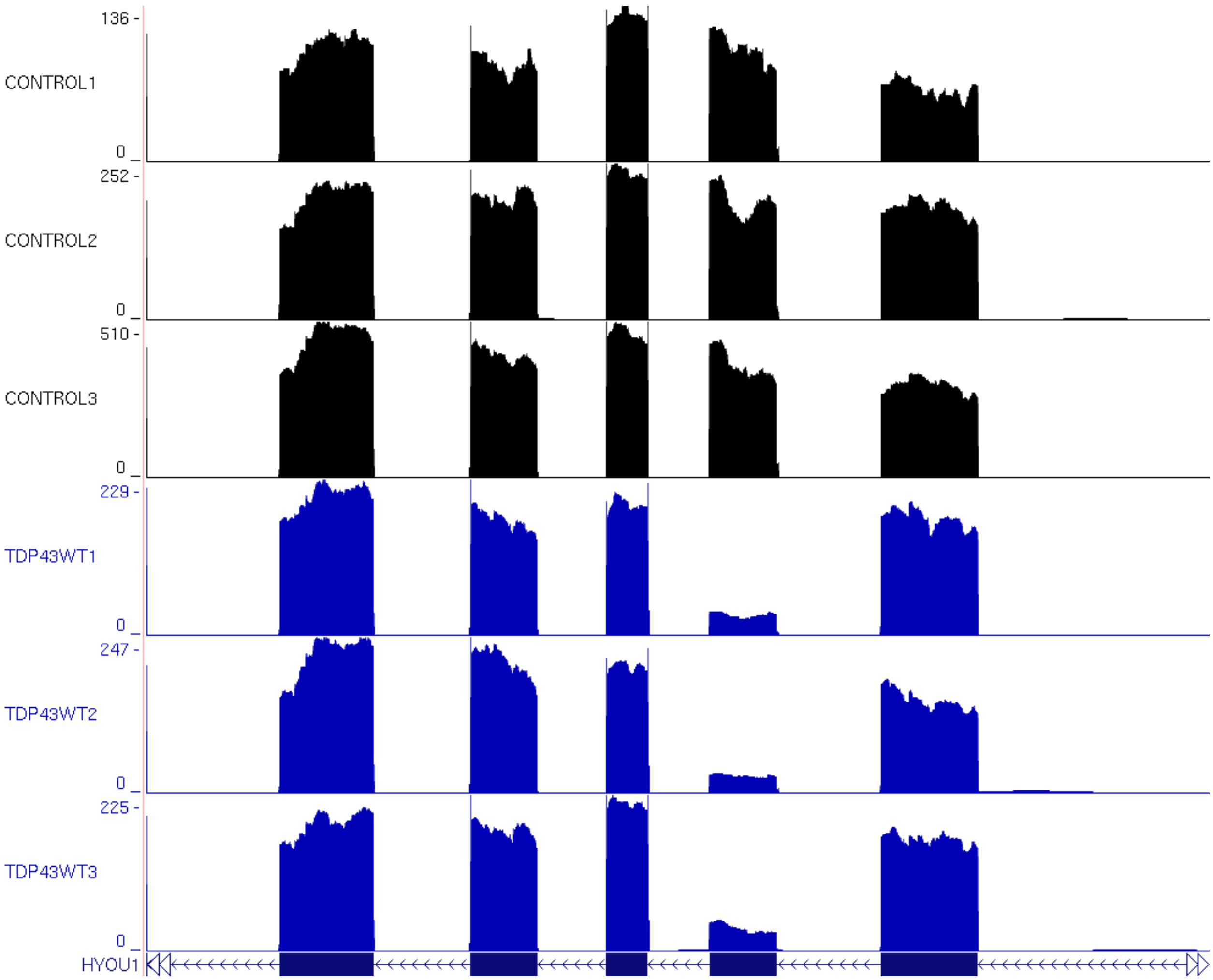

136 -
CONTROL1
0 _
252 -
CONTROL2
0 _
510 -
CONTROL3
0 _
229 -
TDP43WT1
0 _
247 -
TDP43WT2
0 _
225 -
TDP43WT3
0 _
HYOU1

AB

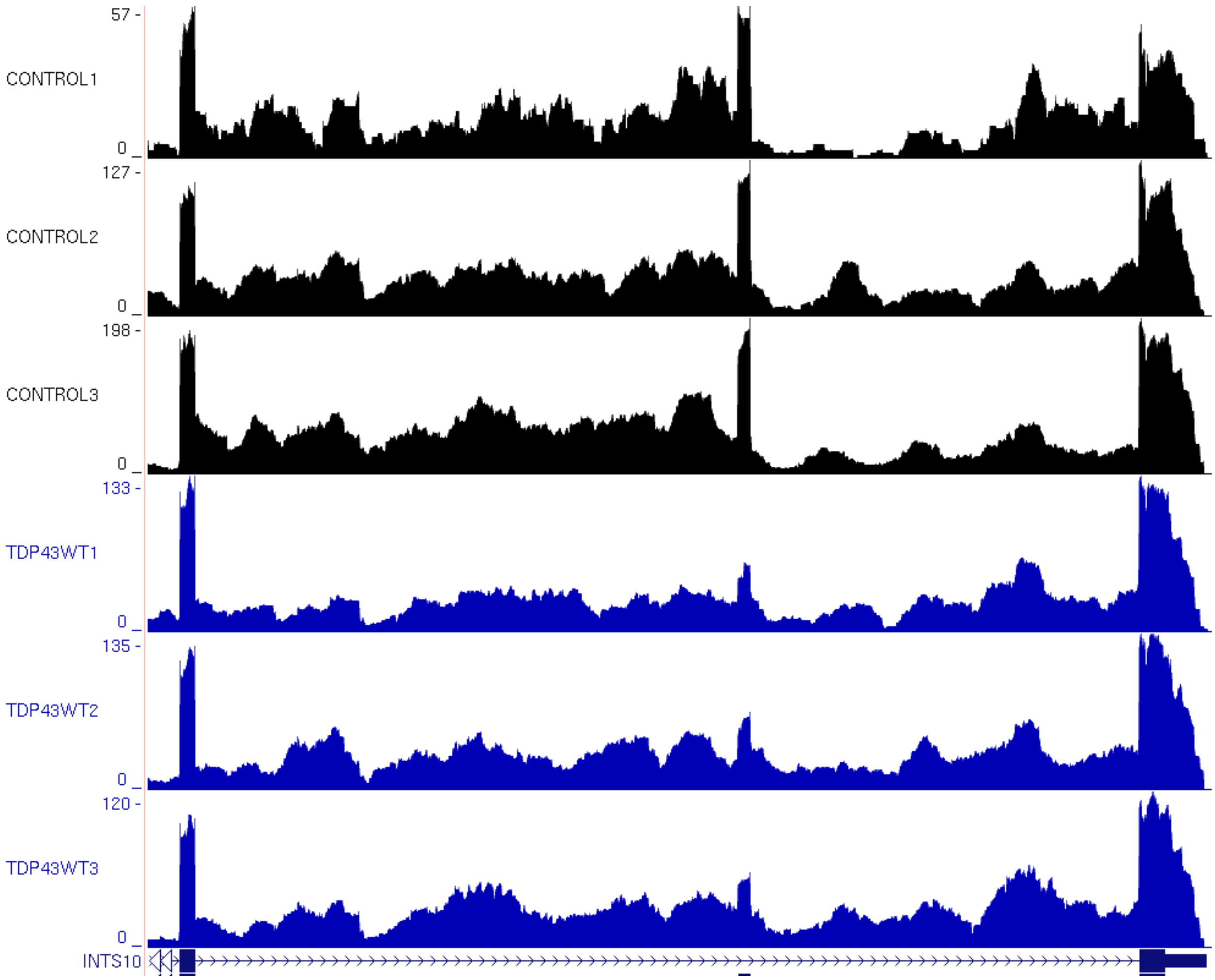

57 -
CONTROL1
0 _
127 -
CONTROL2
0 _
198 -
CONTROL3
0 _
133 -
TDP43WT1
0 _
135 -
TDP43WT2
0 _
120 -
TDP43WT3
0 _
INTS10

AC

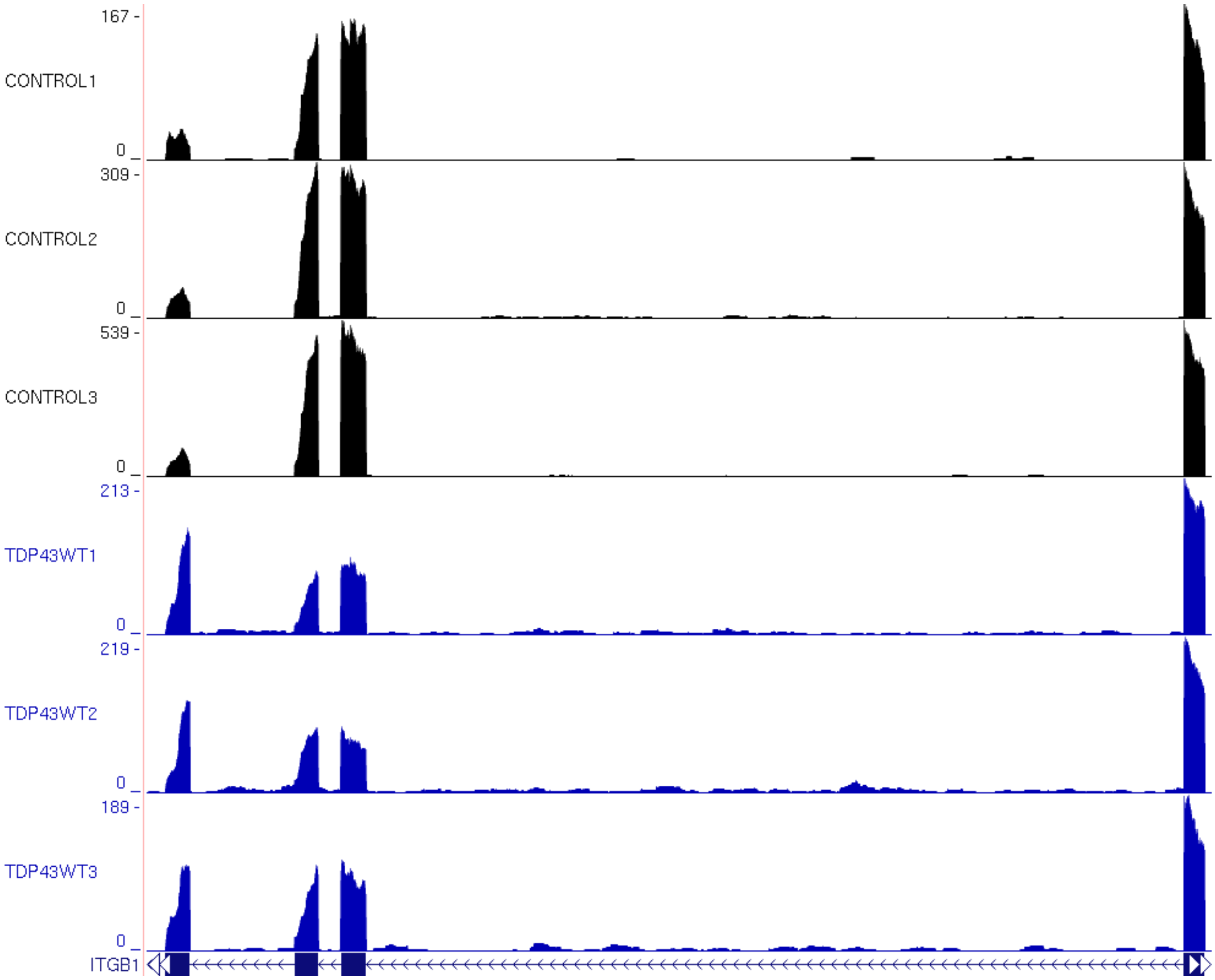
167 -
CONTROL1
0 _
309 -
CONTROL2
0 _
539 -
CONTROL3
0 _
213 -
TDP43WT1
0 _
219 -
TDP43WT2
0 _
189 -
TDP43WT3
0 _
ITGB1

# AD

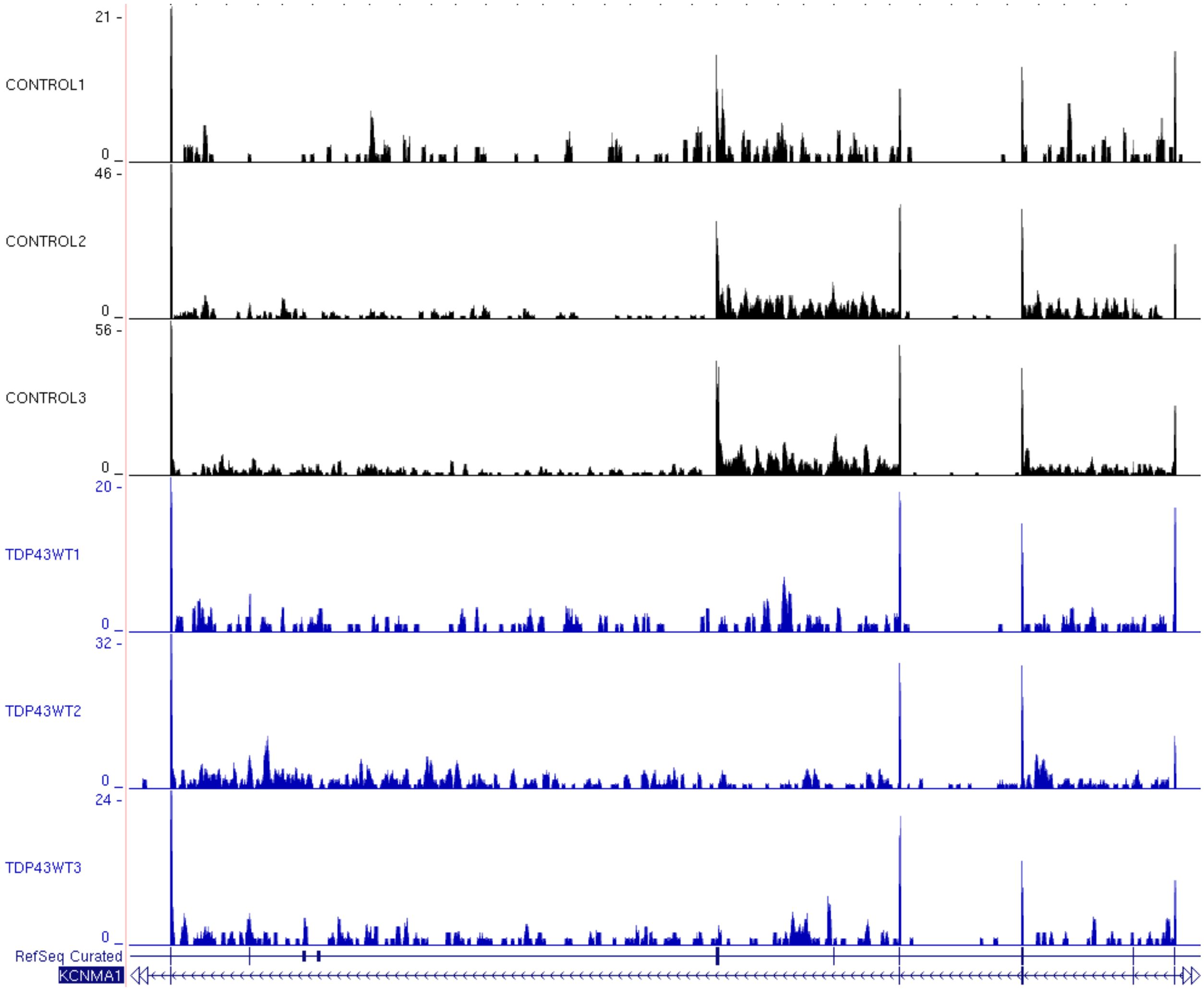

21 -
CONTROL1
0 _
46 -
CONTROL2
0 _
56 -
CONTROL3
0 _
20 -
TDP43WT1
0 _
32 -
TDP43WT2
0 _
24 -
TDP43WT3
0 _
RefSeq Curated
KCNMA1

AE

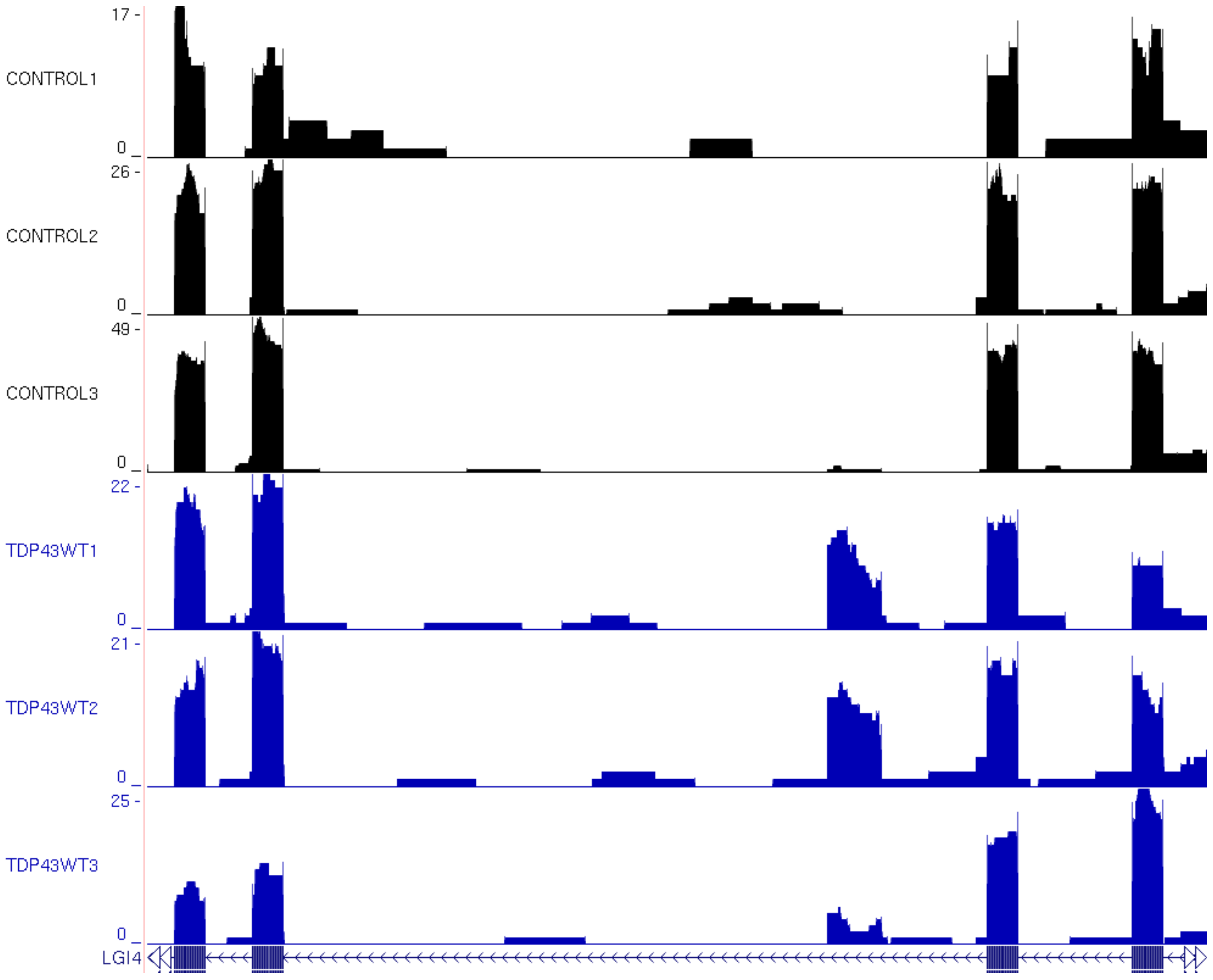

17 -
CONTROL1
0 _
26 -
CONTROL2
0 _
49 -
CONTROL3
0 _
22 -
TDP43WT1
0 _
21 -
TDP43WT2
0 _
25 -
TDP43WT3
0 _
LGI4

AF

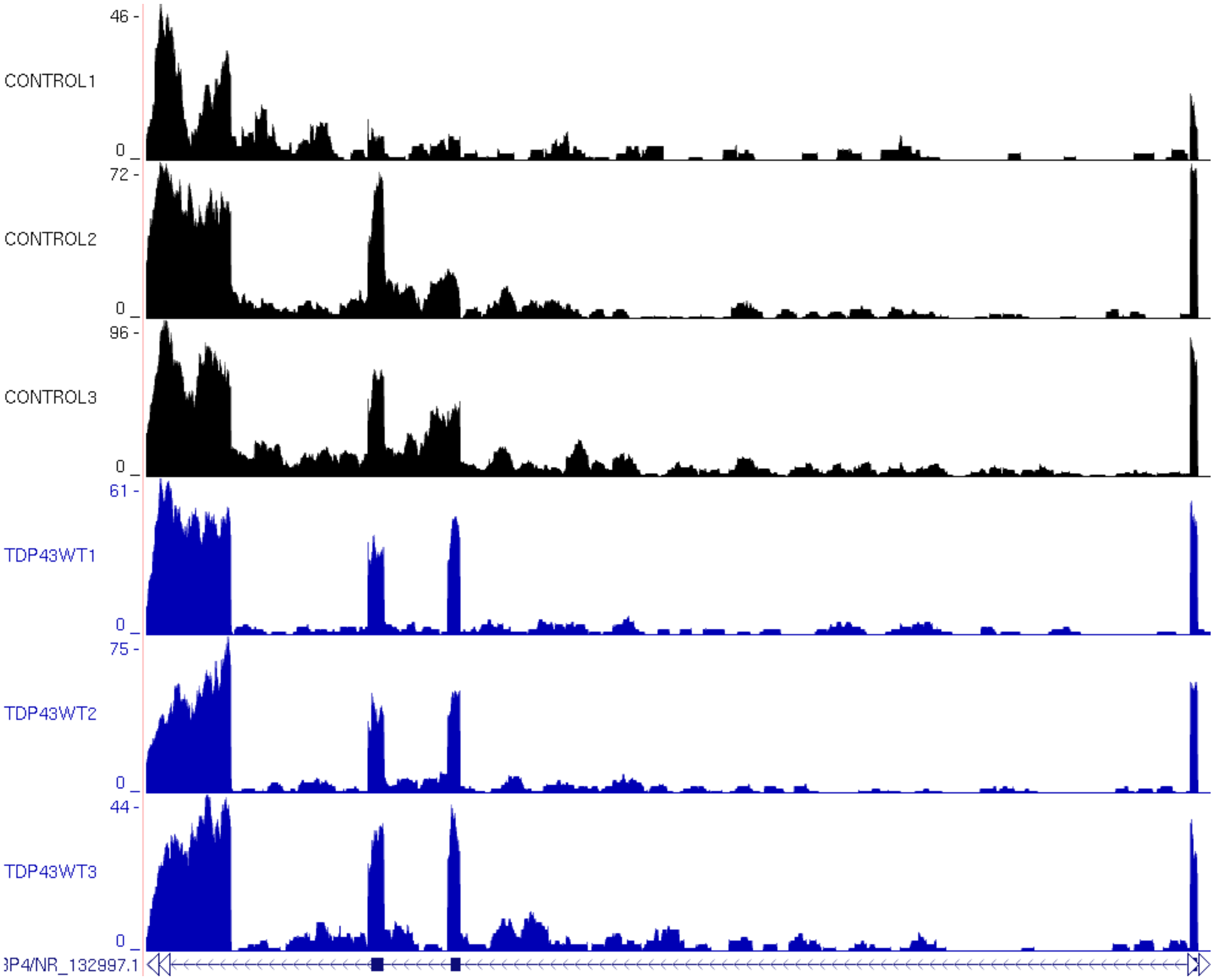

46 -
CONTROL1
0 _
72 -
CONTROL2
0 _
96 -
CONTROL3
0 _
61 -
TDP43WT1
0 _
75 -
TDP43WT2
0 _
44 -
TDP43WT3
0 _
3P4/NR_132997.1

# AG

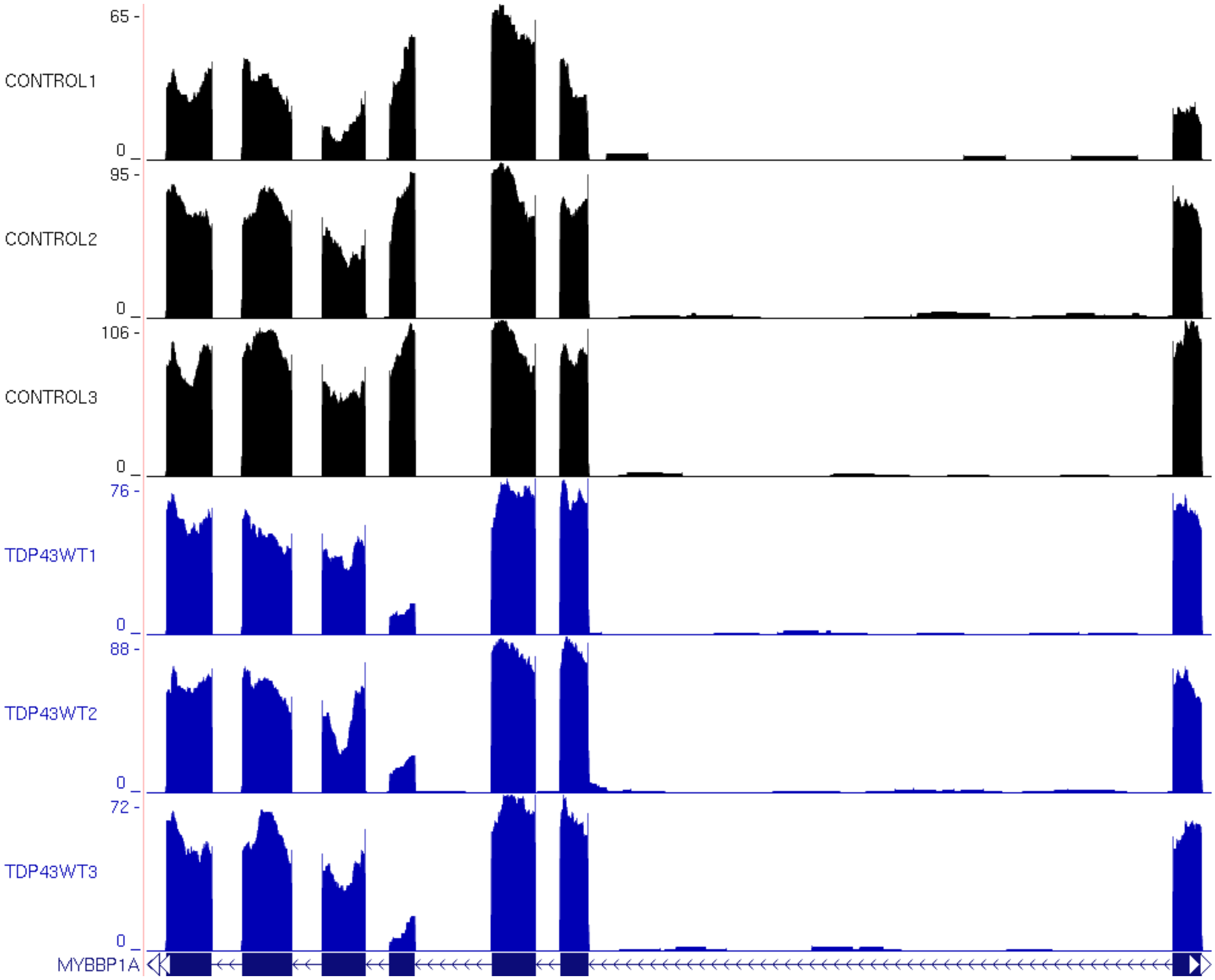
65 -
CONTROL1
0
95 -
CONTROL2
0
106 -
CONTROL3
0
76 -
TDP43WT1
0
88 -
TDP43WT2
0
72 -
TDP43WT3
0
MYBBP1A

AH

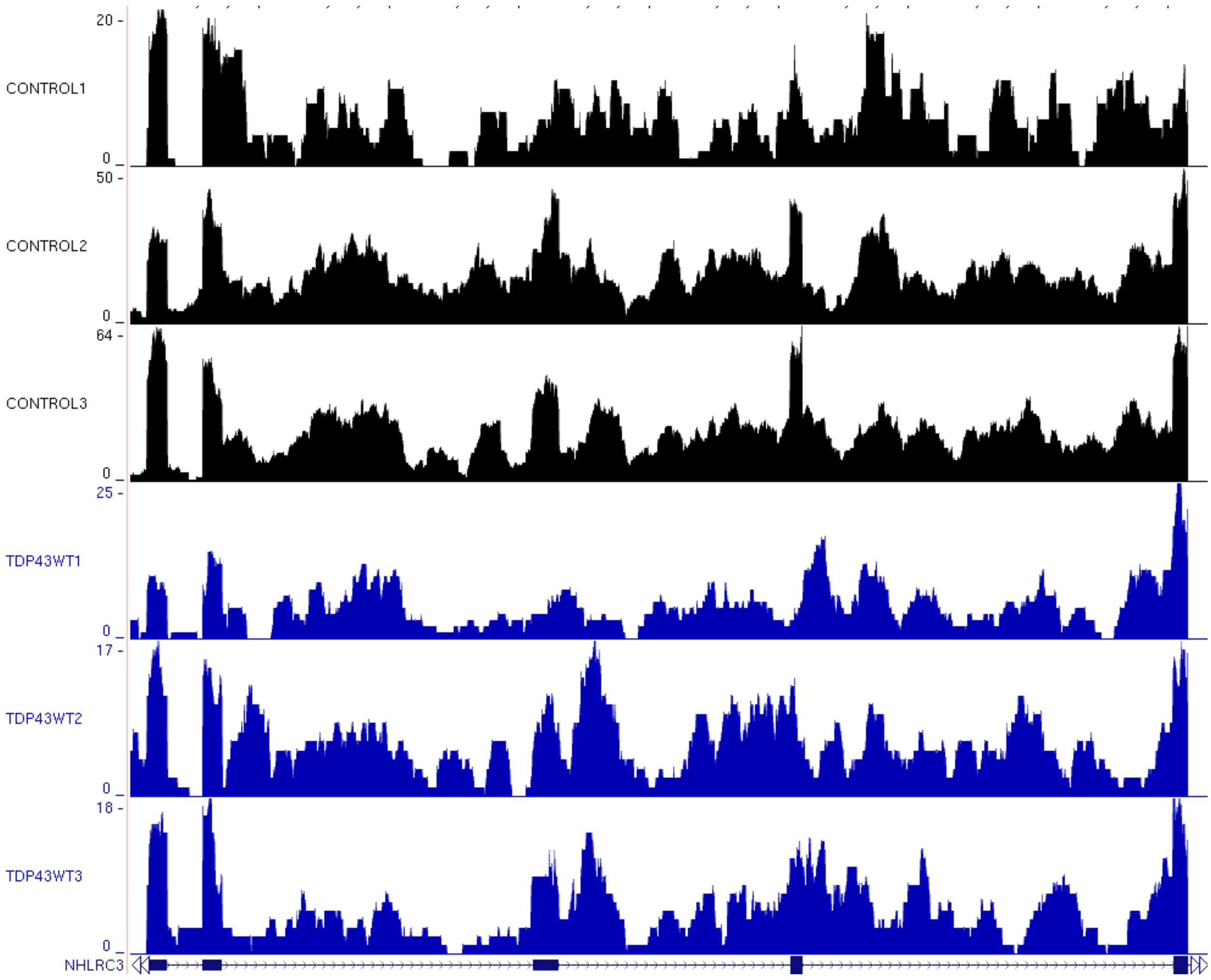
20 -
CONTROL1
0
50 -
CONTROL2
0
64 -
CONTROL3
0
25 -
TDP43WT1
0
17 -
TDP43WT2
0
18 -
TDP43WT3
0
NHLRC3

AI

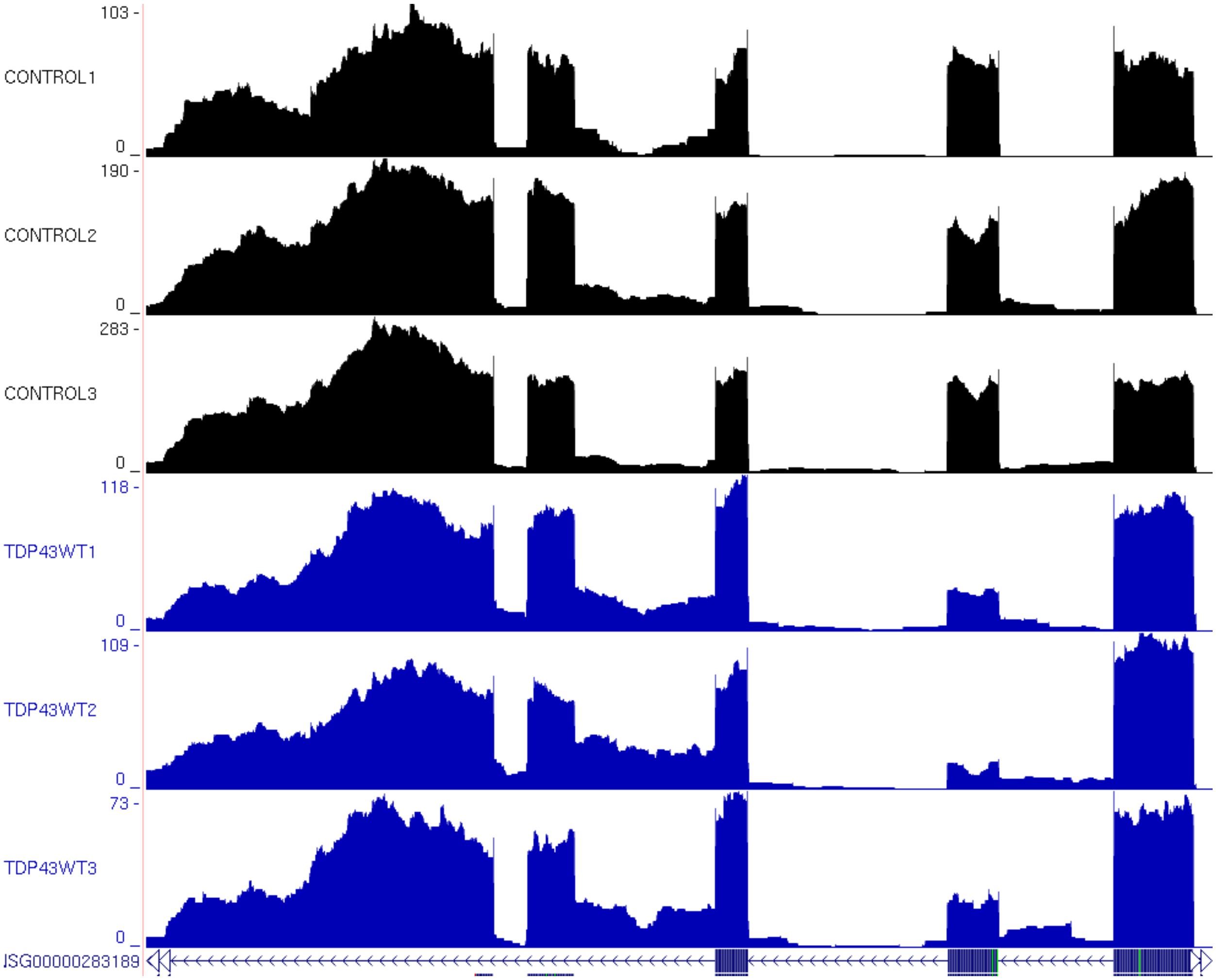

103 -
CONTROL1
0
190 -
CONTROL2
0
283 -
CONTROL3
0
118 -
TDP43WT1
0
109 -
TDP43WT2
0
73 -
TDP43WT3
0
JSG00000283189

AJ

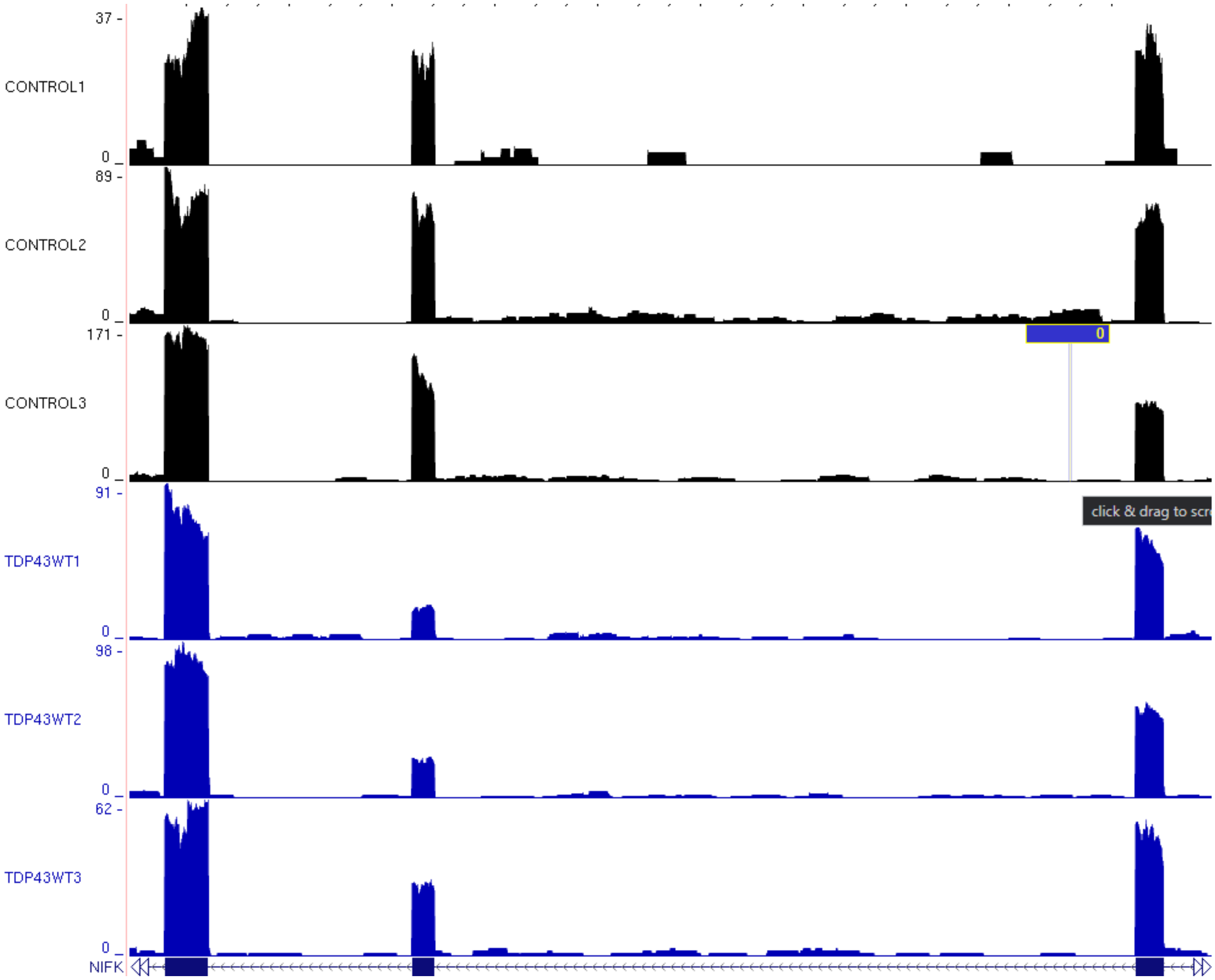
37 -
CONTROL1
0 _
89 -
CONTROL2
0 _
171 -
CONTROL3
0 _
0
91 -
TDP43WT1
0 _
98 -
TDP43WT2
0 _
62 -
TDP43WT3
0 _
NIFK

AK

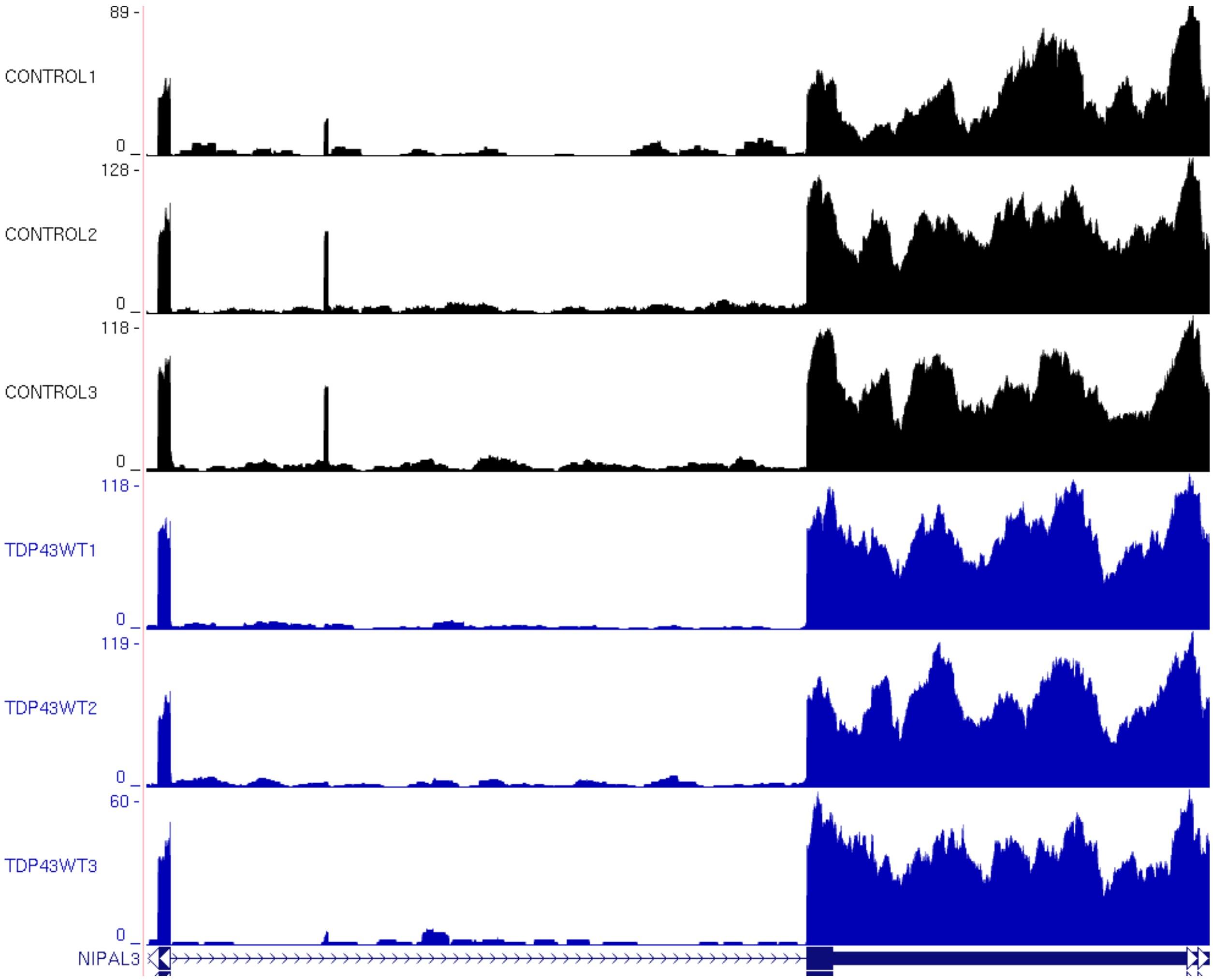
89 -
CONTROL1
0
128 -
CONTROL2
0
118 -
CONTROL3
0
118 -
TDP43WT1
0
119 -
TDP43WT2
0
60 -
TDP43WT3
0
NIPAL3

AL

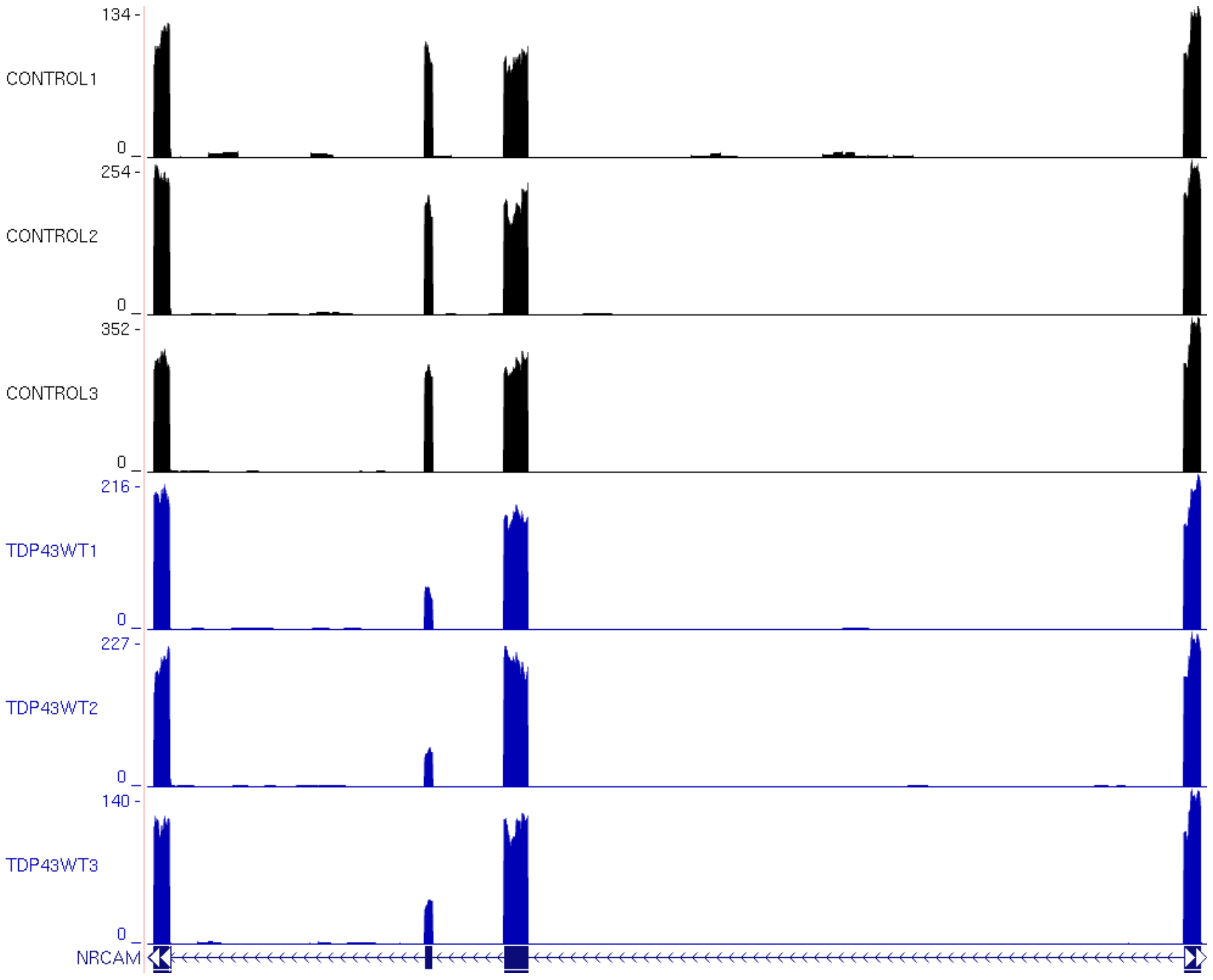
134 -
CONTROL1
0
254 -
CONTROL2
0
352 -
CONTROL3
0
216 -
TDP43WT1
0
227 -
TDP43WT2
0
140 -
TDP43WT3
0
NRCAM

# AM

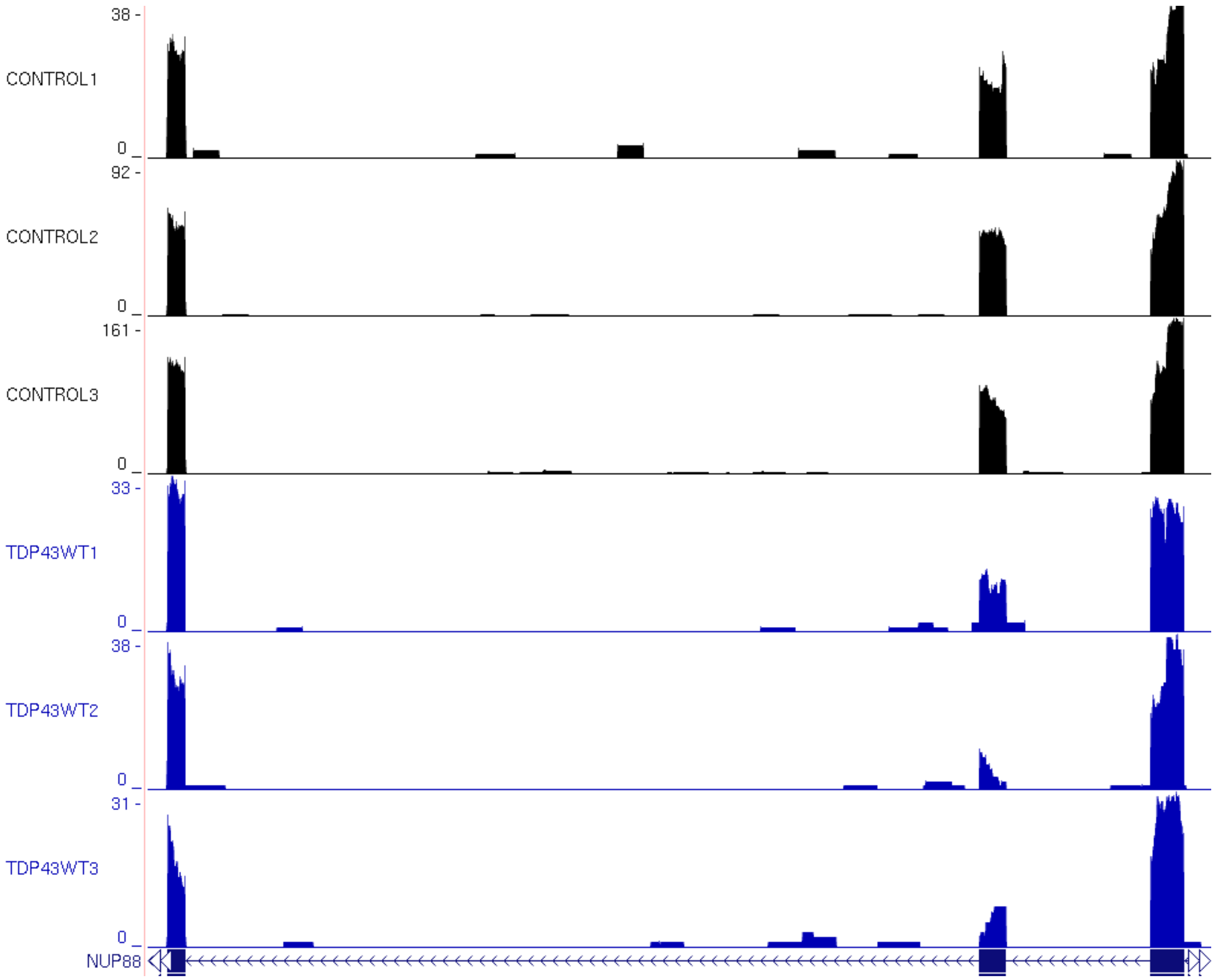

38 -
CONTROL1
0
92 -
CONTROL2
0
161 -
CONTROL3
0
33 -
TDP43WT1
0
38 -
TDP43WT2
0
31 -
TDP43WT3
0
NUP88

AN

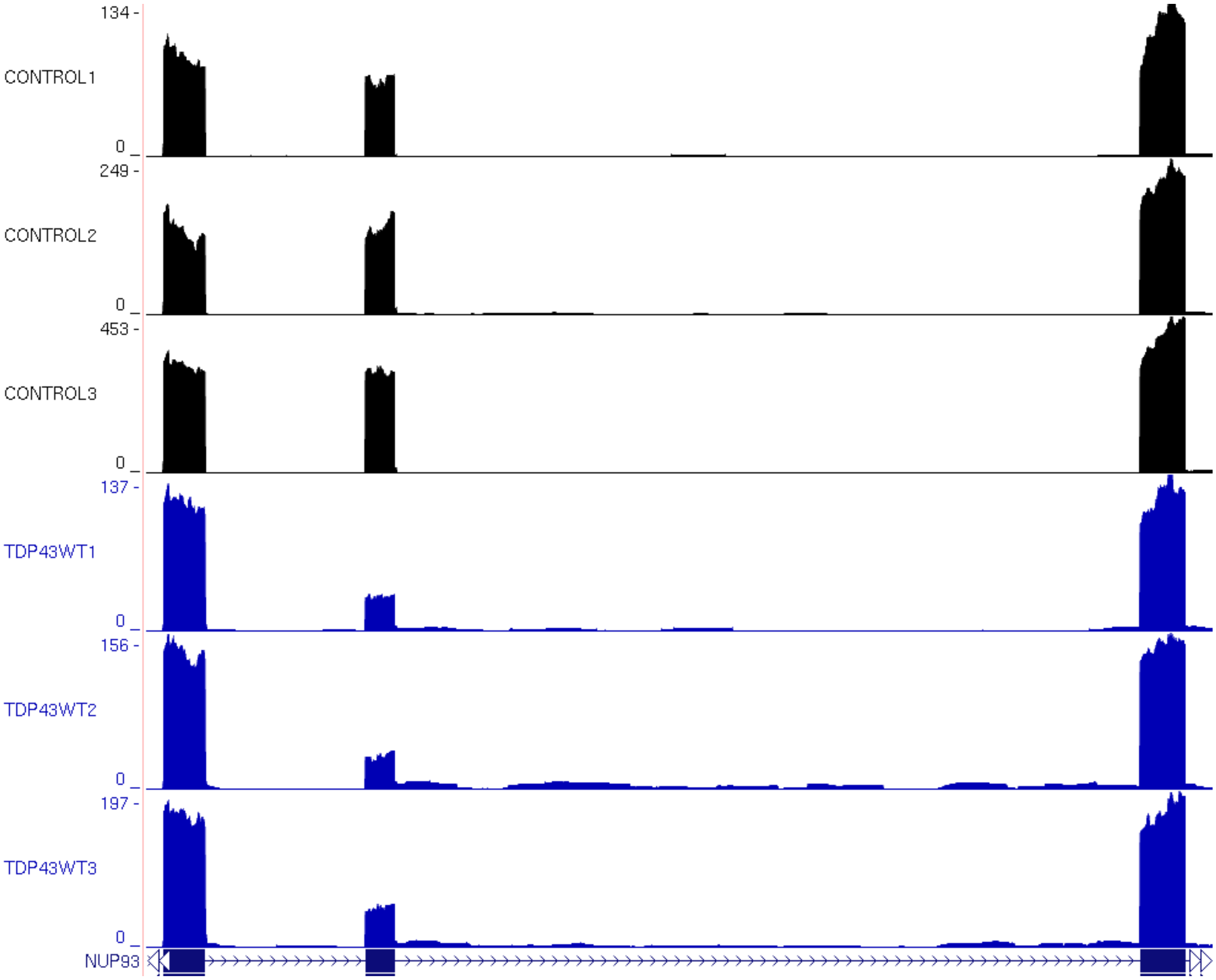
134 -
CONTROL1
0
249 -
CONTROL2
0
453 -
CONTROL3
0
137 -
TDP43WT1
0
156 -
TDP43WT2
0
197 -
TDP43WT3
0
NUP93

AO

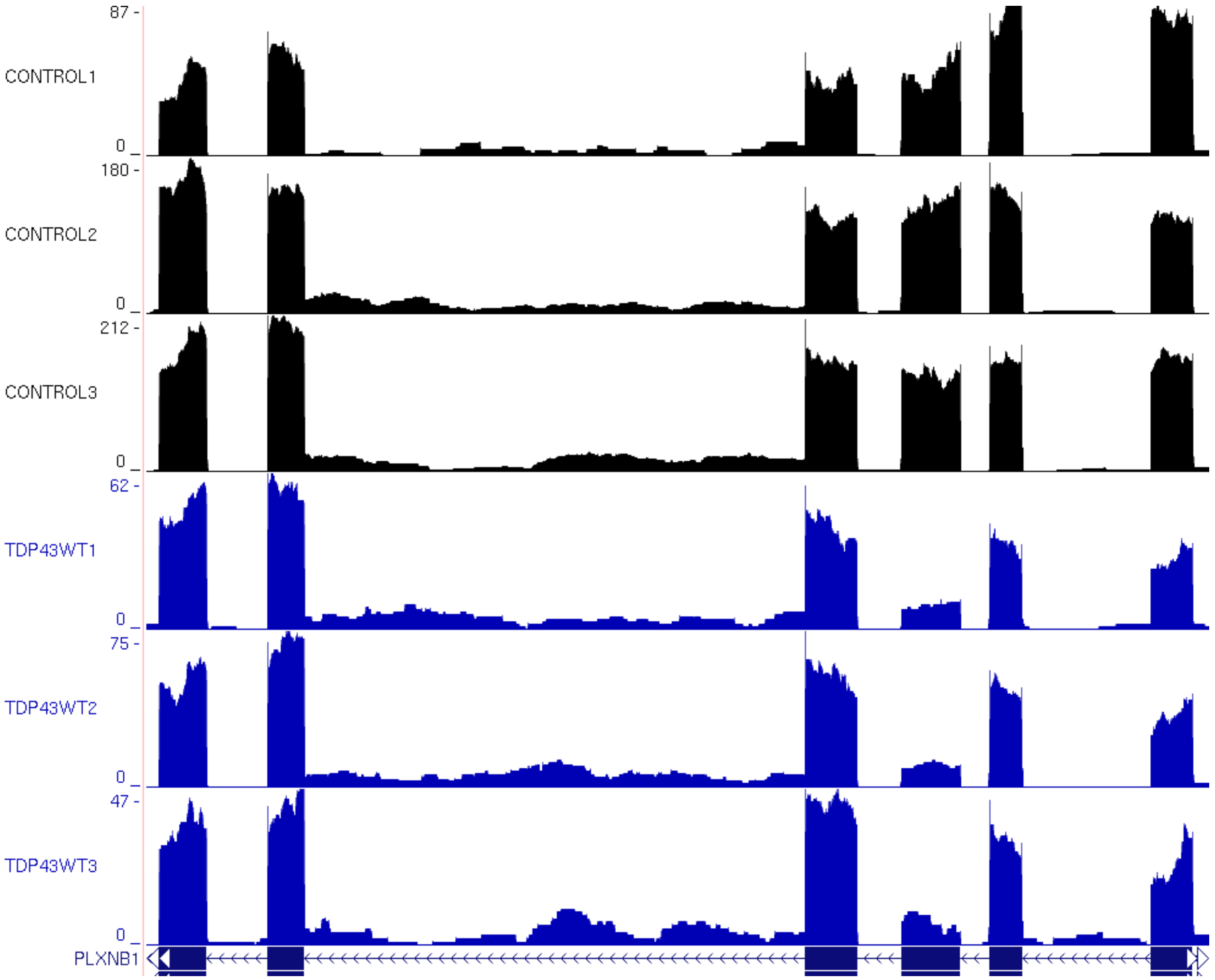

87 -
CONTROL1
0 _
180 -
CONTROL2
0 _
212 -
CONTROL3
0 _
62 -
TDP43WT1
0 _
75 -
TDP43WT2
0 _
47 -
TDP43WT3
0 _
PLXNB1

AP

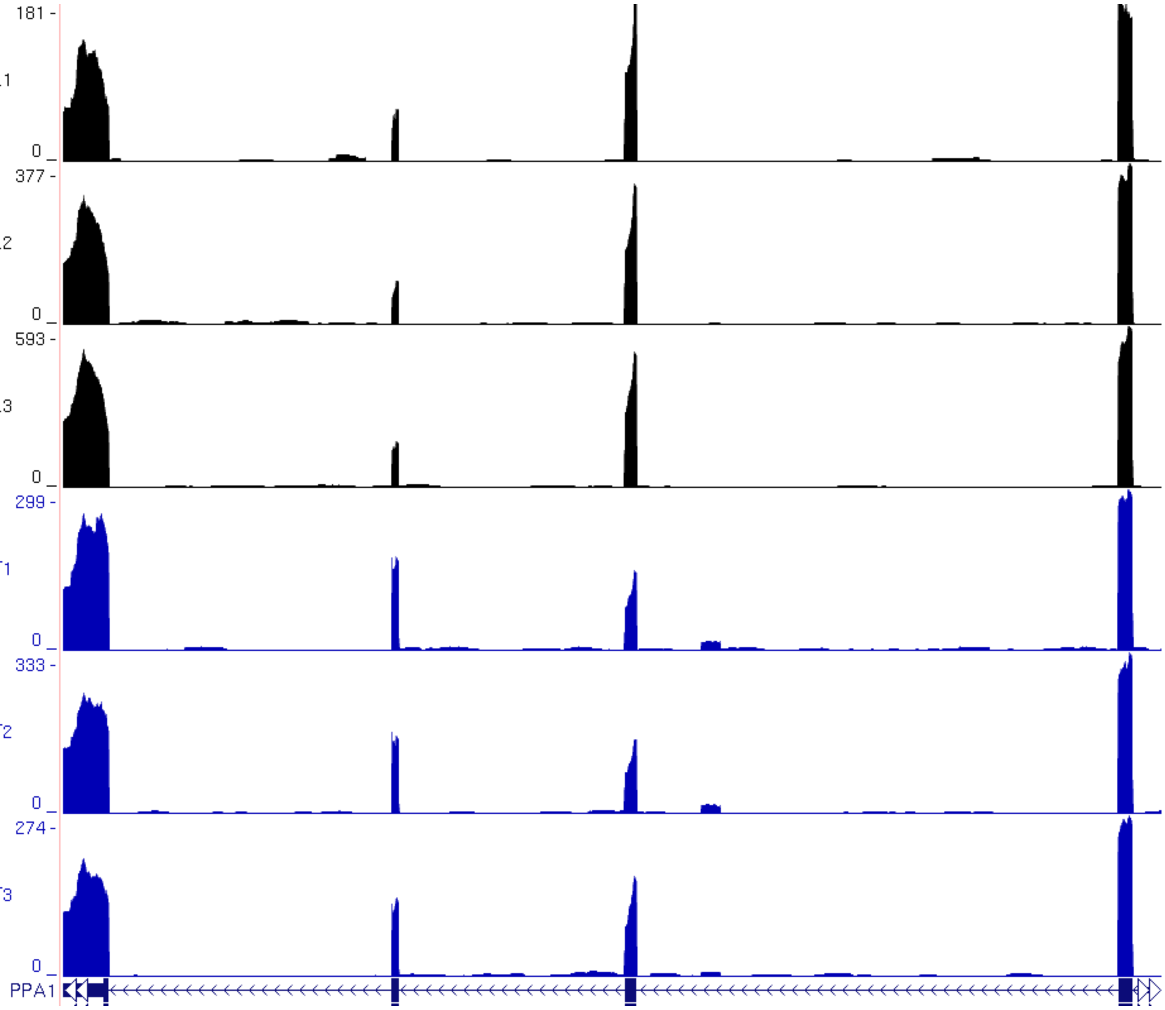
181 -
CONTROL1
0
377 -
CONTROL2
0
593 -
CONTROL3
0
299 -
TDP43WT1
0
333 -
TDP43WT2
0
274 -
TDP43WT3
0
PPA1

AQ

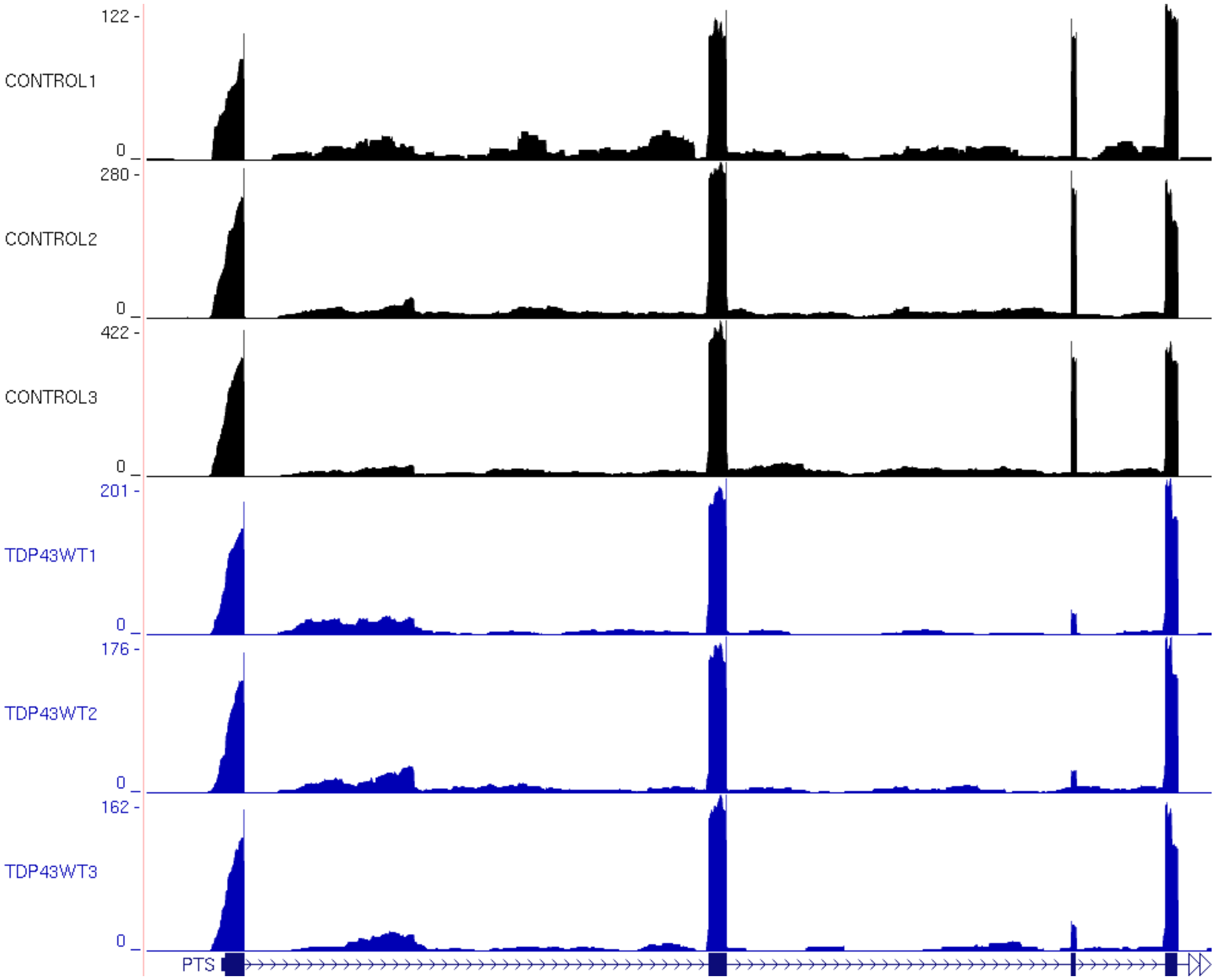

122 -
CONTROL1
0 _
280 -
CONTROL2
0 _
422 -
CONTROL3
0 _
201 -
TDP43WT1
0 _
176 -
TDP43WT2
0 _
162 -
TDP43WT3
0 _
PTS

# AR

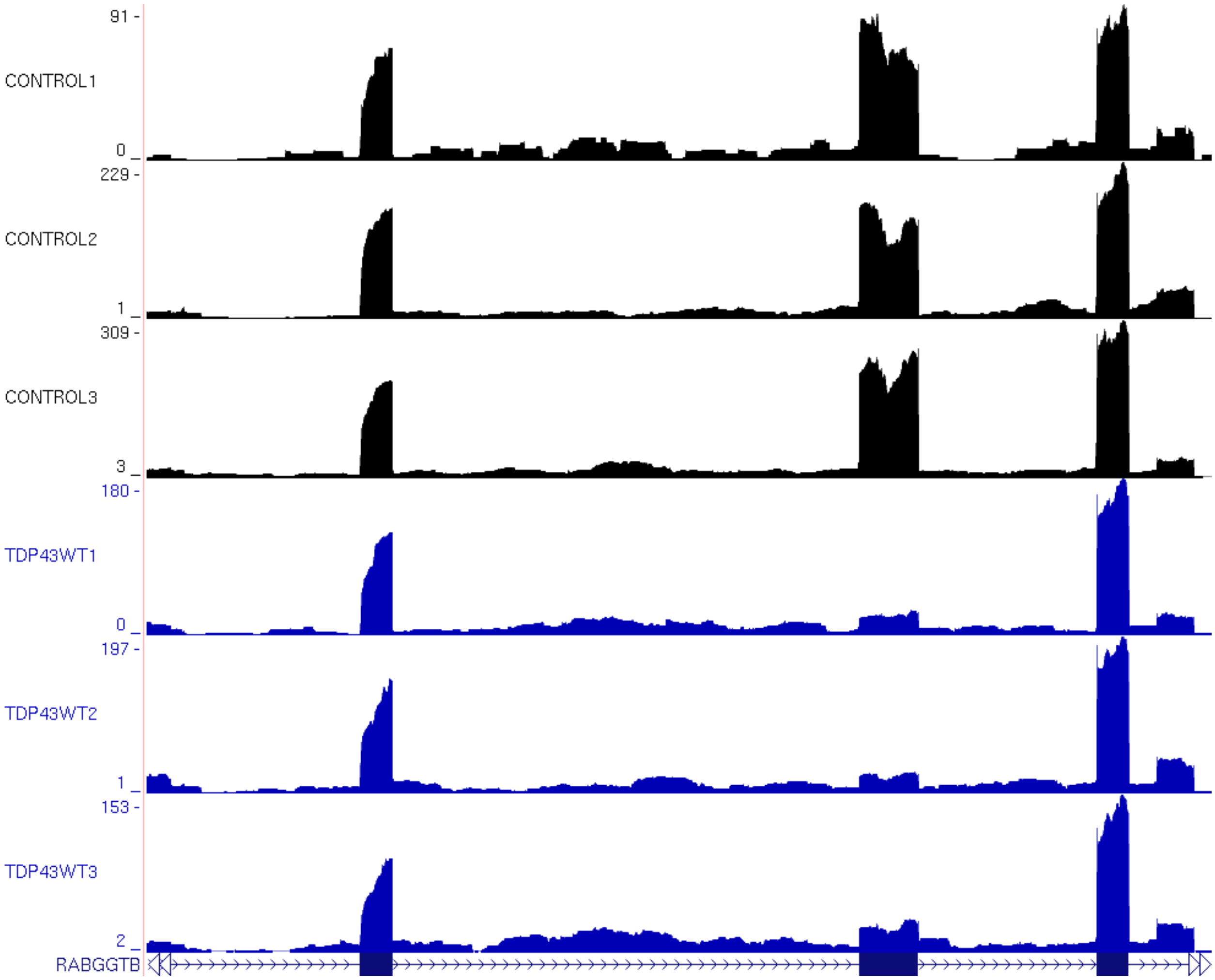

91 -
CONTROL1
0 _
229 -
CONTROL2
1 _
309 -
CONTROL3
3 _
180 -
TDP43WT1
0 _
197 -
TDP43WT2
1 _
153 -
TDP43WT3
2 _
RABGGTB

AS

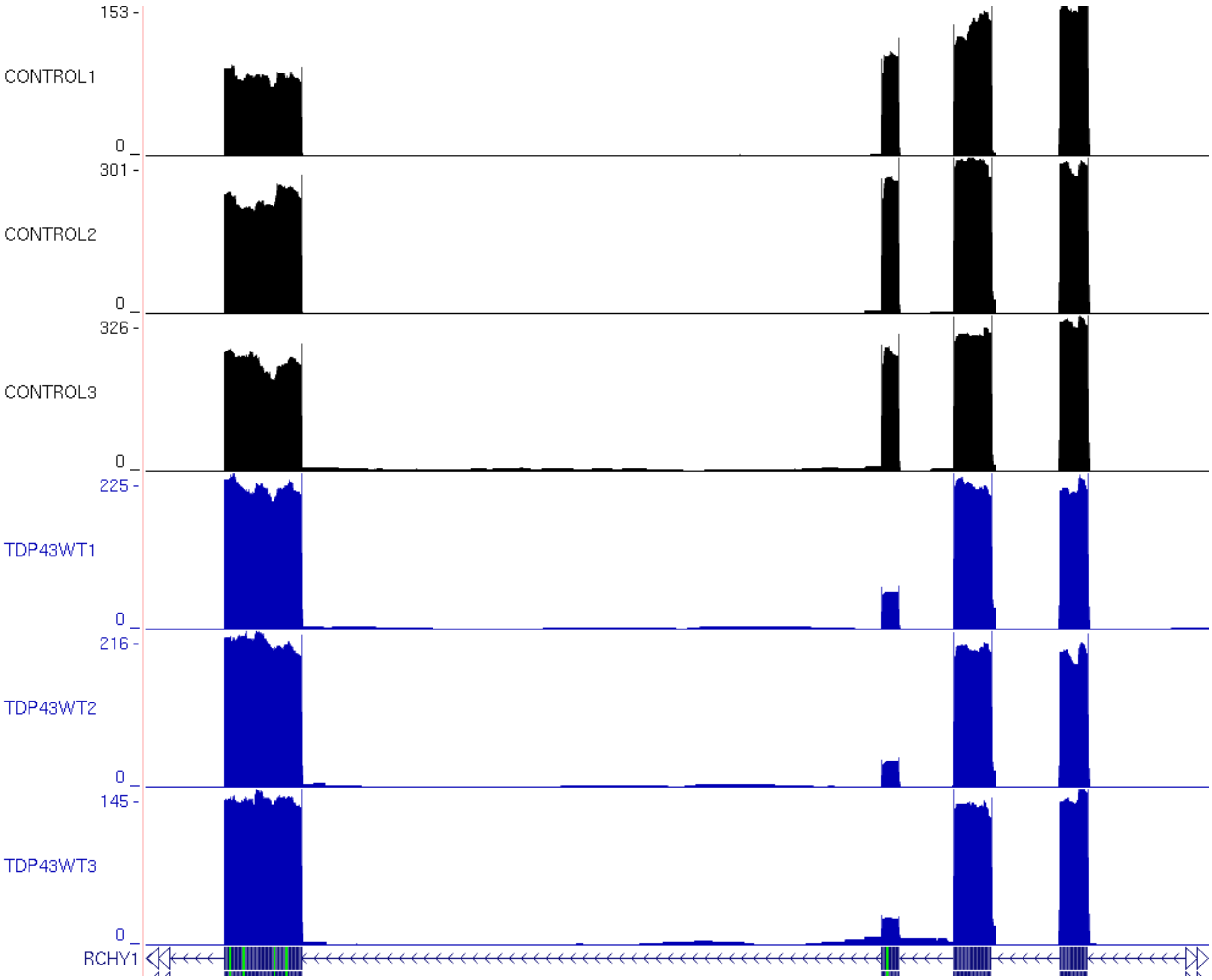

153 -
CONTROL1
0
301 -
CONTROL2
0
326 -
CONTROL3
0
225 -
TDP43WT1
0
216 -
TDP43WT2
0
145 -
TDP43WT3
0
RCHY1

# AT

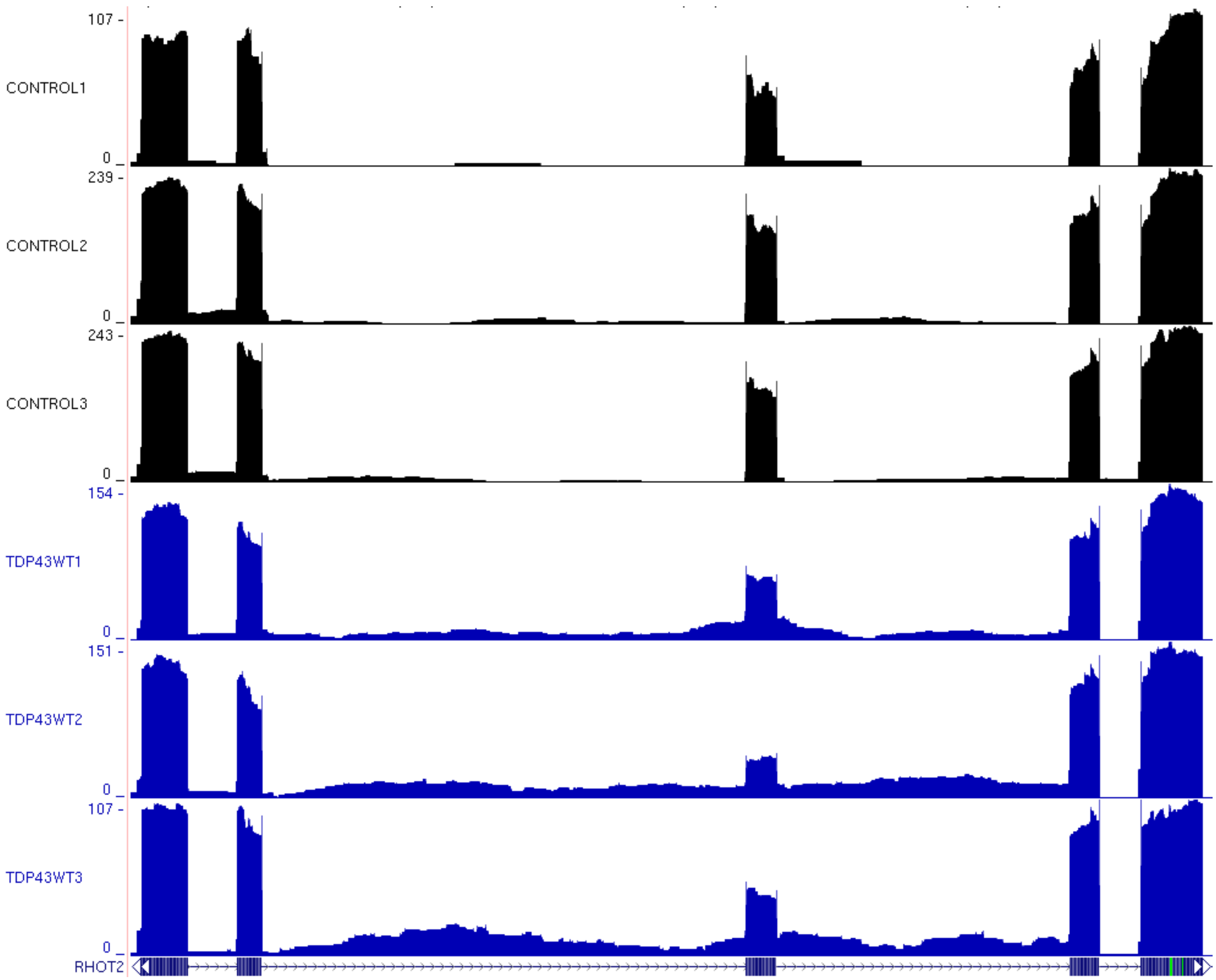
107 -
CONTROL1
0 _
239 -
CONTROL2
0 _
243 -
CONTROL3
0 _
154 -
TDP43WT1
0 _
151 -
TDP43WT2
0 _
107 -
TDP43WT3
0 _
RHOT2

# AU

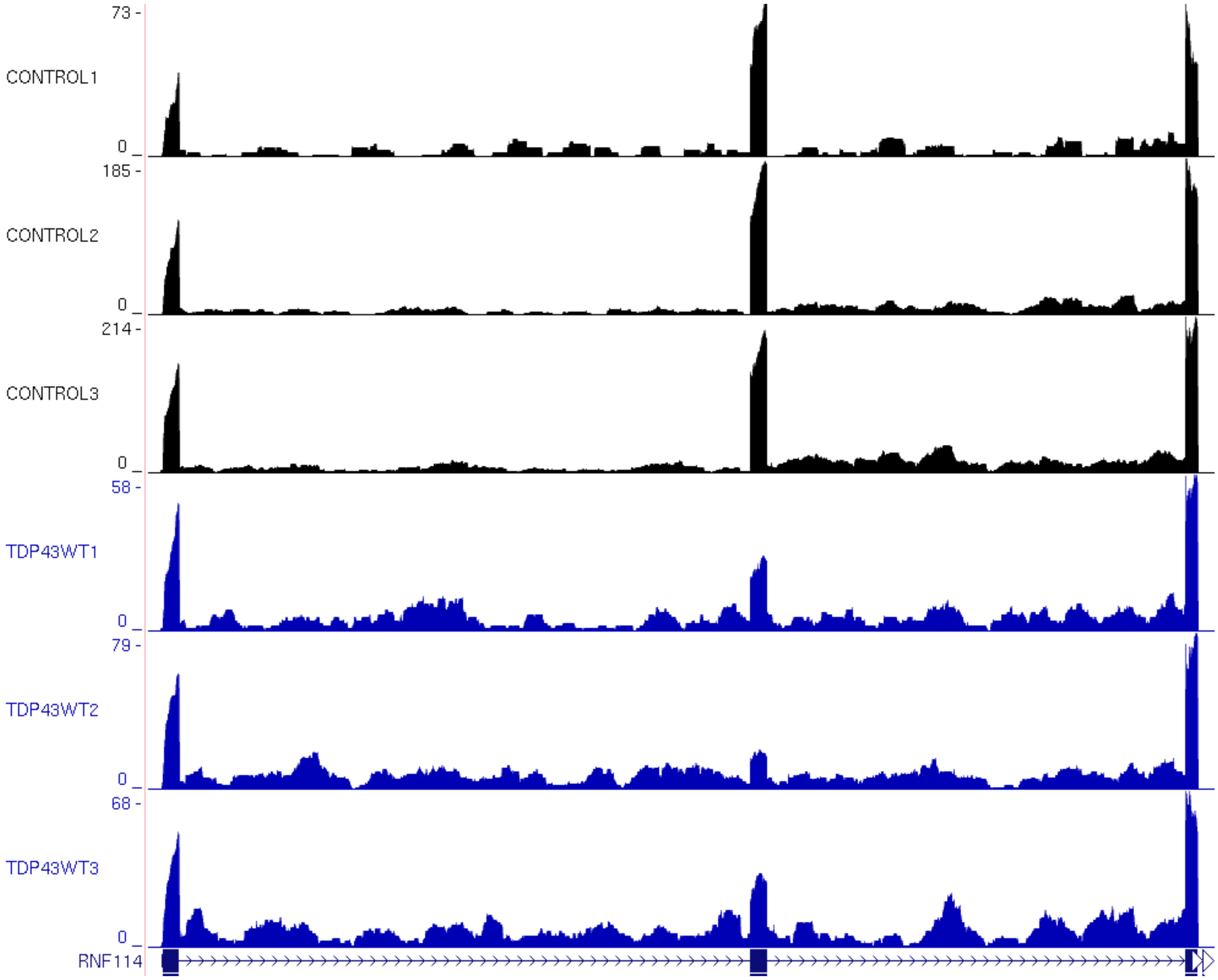
73 -
CONTROL1
0
185 -
CONTROL2
0
214 -
CONTROL3
0
58 -
TDP43WT1
0
79 -
TDP43WT2
0
68 -
TDP43WT3
0
RNF114

# AV

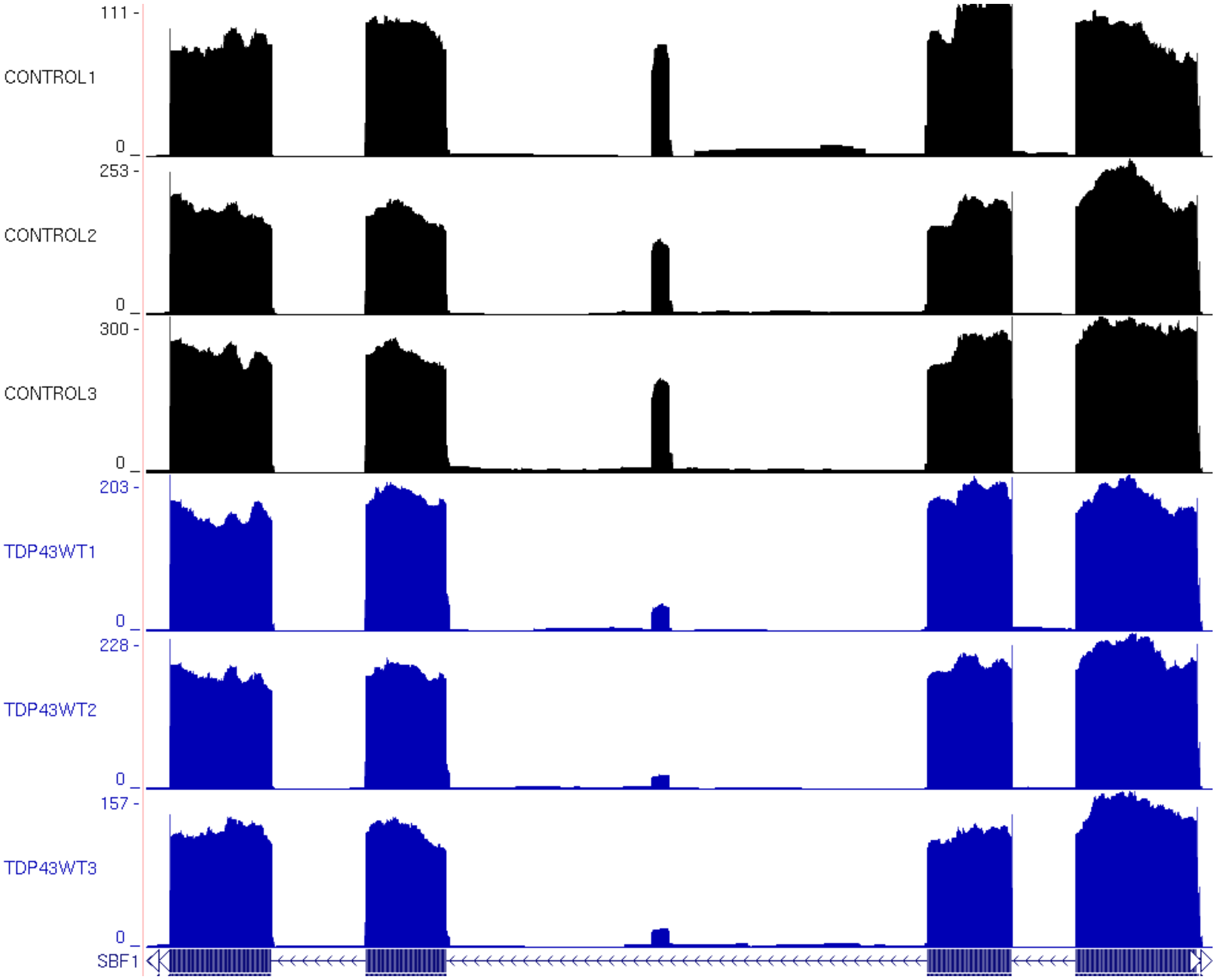

111 -
CONTROL1
0 _
253 -
CONTROL2
0 _
300 -
CONTROL3
0 _
203 -
TDP43WT1
0 _
228 -
TDP43WT2
0 _
157 -
TDP43WT3
0 _
SBF1

AW

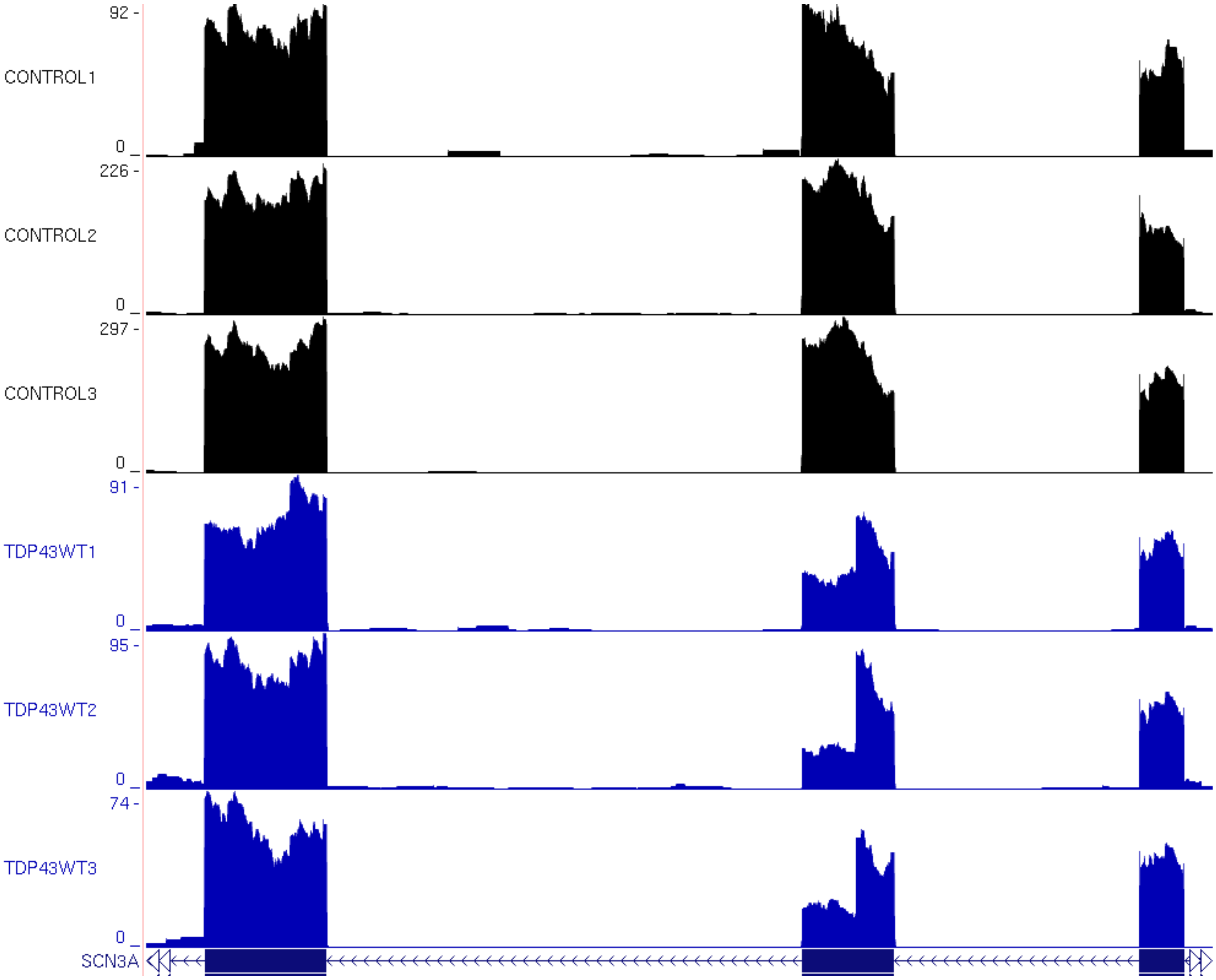
92 -
CONTROL1
0
226 -
CONTROL2
0
297 -
CONTROL3
0
91 -
TDP43WT1
0
95 -
TDP43WT2
0
74 -
TDP43WT3
0
SCN3A

AX

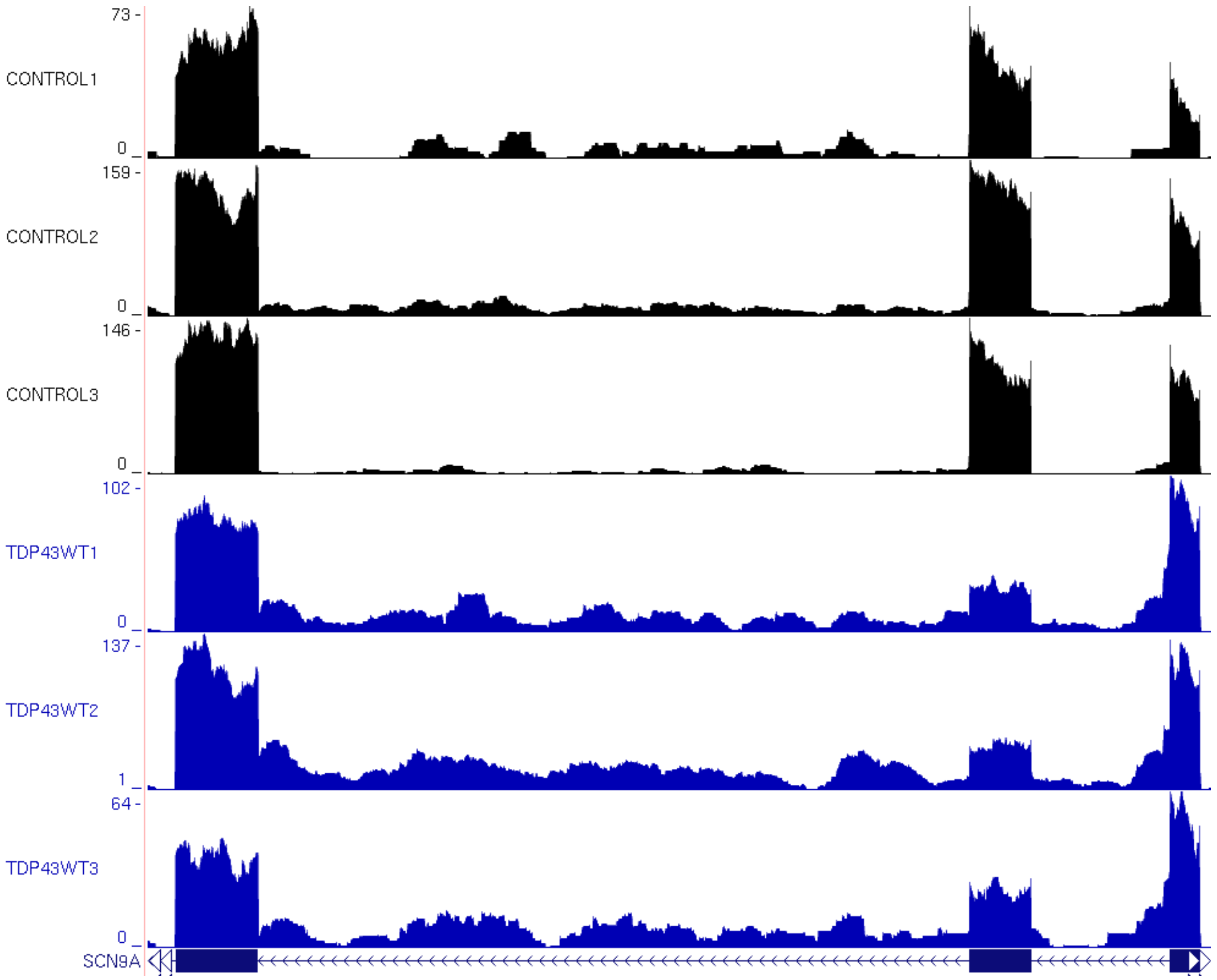

73 -
CONTROL1
0
159 -
CONTROL2
0
146 -
CONTROL3
0
102 -
TDP43WT1
0
137 -
TDP43WT2
1
64 -
TDP43WT3
0
SCN9A

# AY

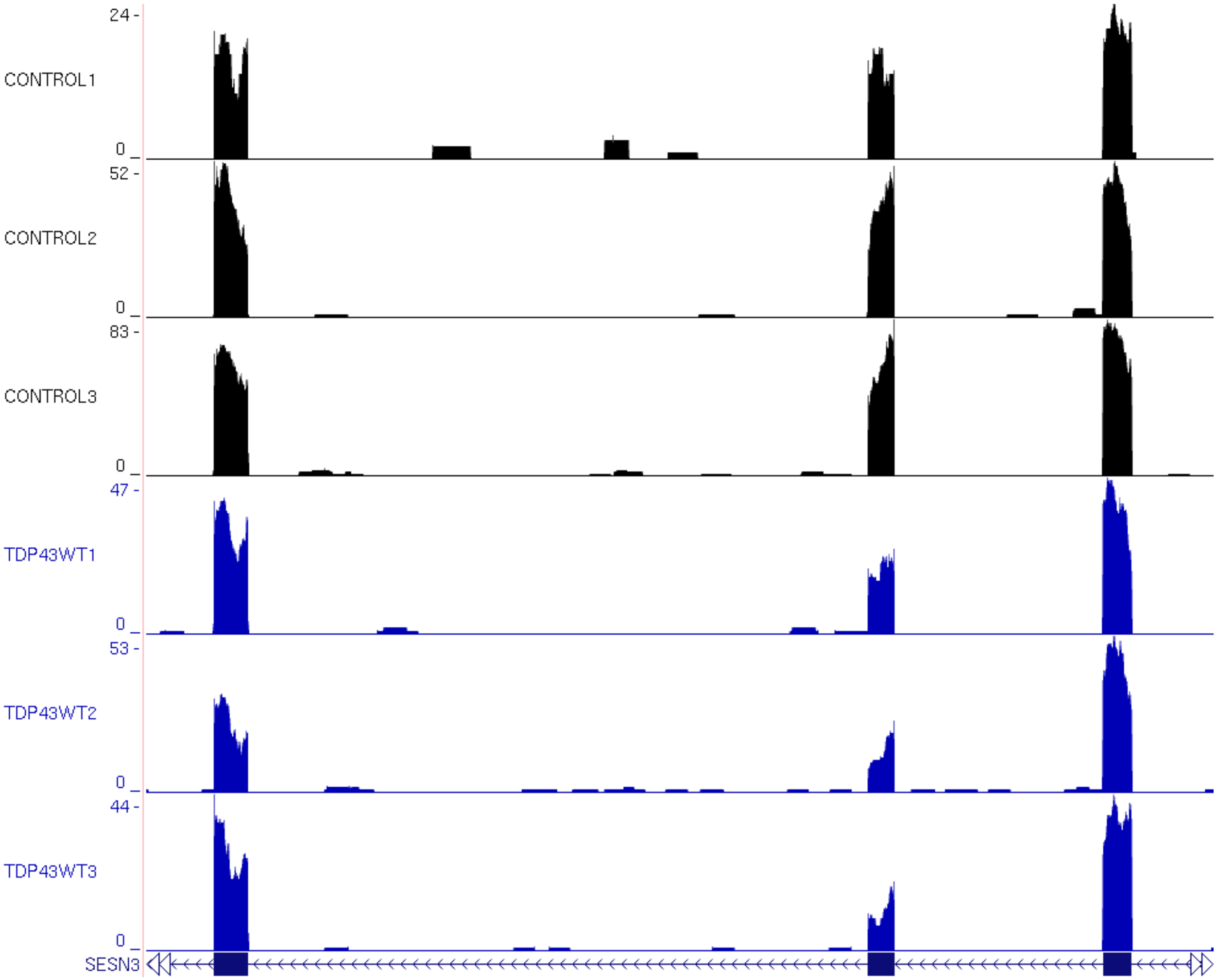

24 -
CONTROL1
0
52 -
CONTROL2
0
83 -
CONTROL3
0
47 -
TDP43WT1
0
53 -
TDP43WT2
0
44 -
TDP43WT3
0
SESN3

AZ

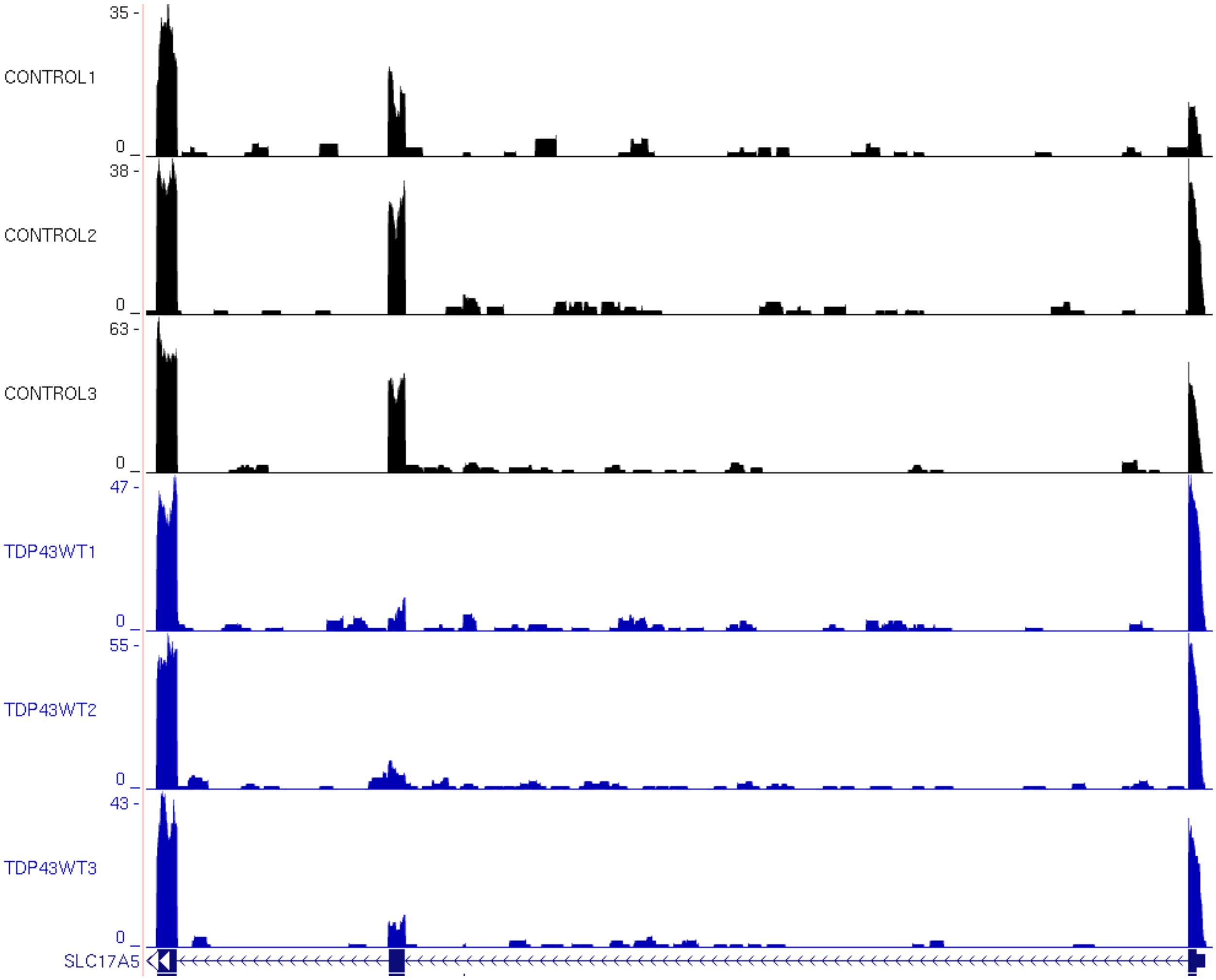
35 -
CONTROL1
0
38 -
CONTROL2
0
63 -
CONTROL3
0
47 -
TDP43WT1
0
55 -
TDP43WT2
0
43 -
TDP43WT3
0
SLC17A5

AAA

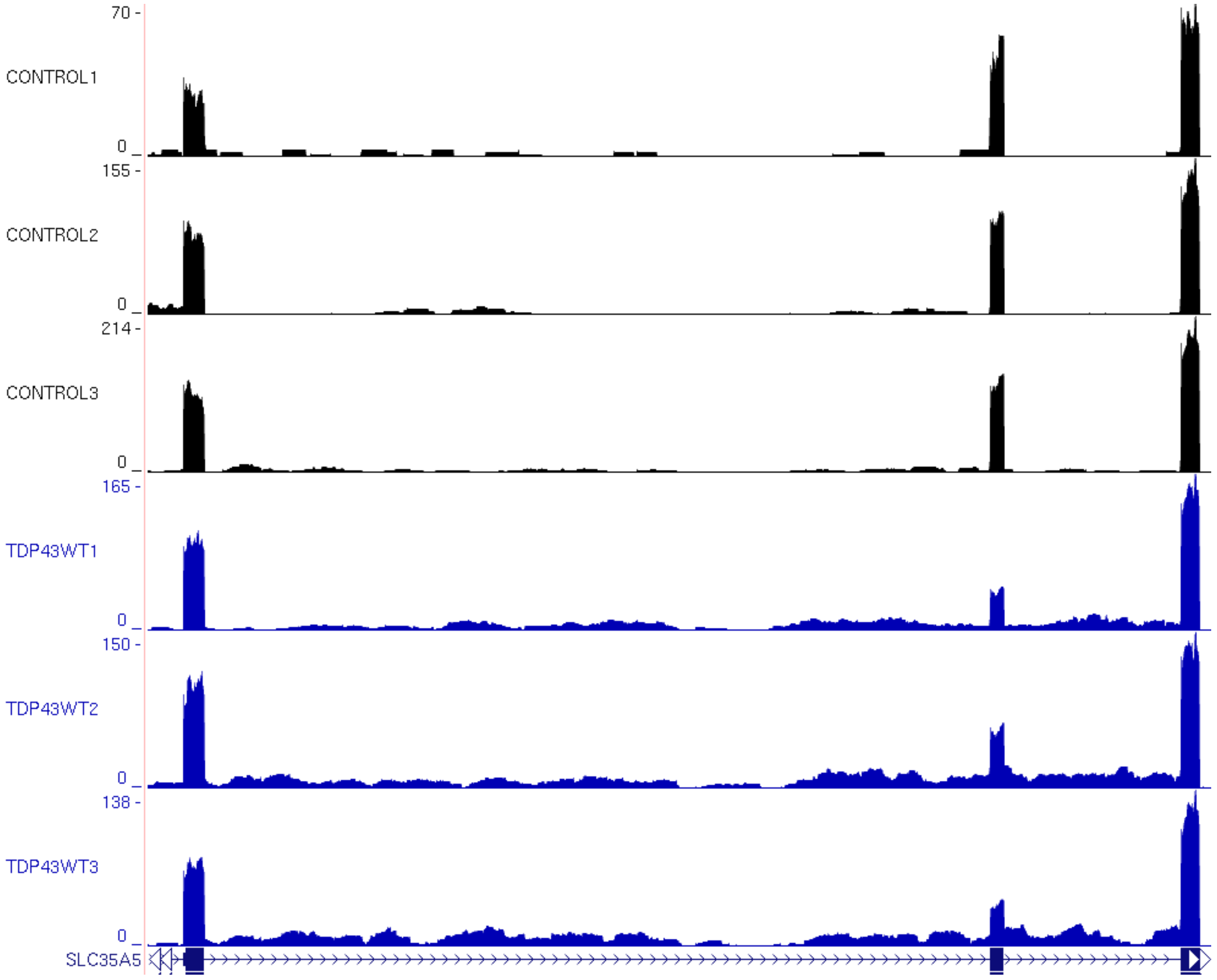

70 -
CONTROL1
0
155 -
CONTROL2
0
214 -
CONTROL3
0
165 -
TDP43WT1
0
150 -
TDP43WT2
0
138 -
TDP43WT3
0
SLC35A5

AAB

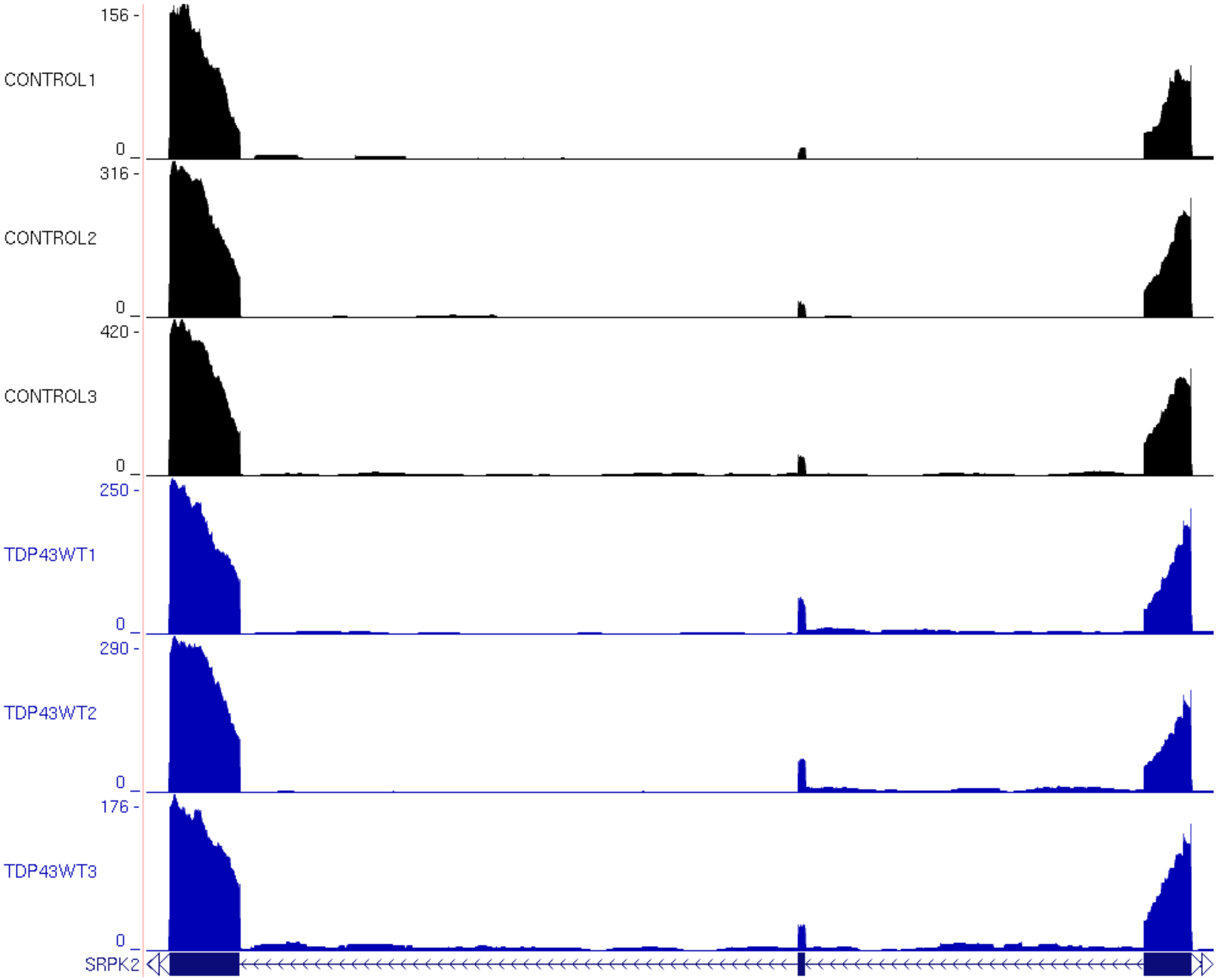
156 -
CONTROL1
0
316 -
CONTROL2
0
420 -
CONTROL3
0
250 -
TDP43WT1
0
290 -
TDP43WT2
0
176 -
TDP43WT3
0
SRPK2

AAC

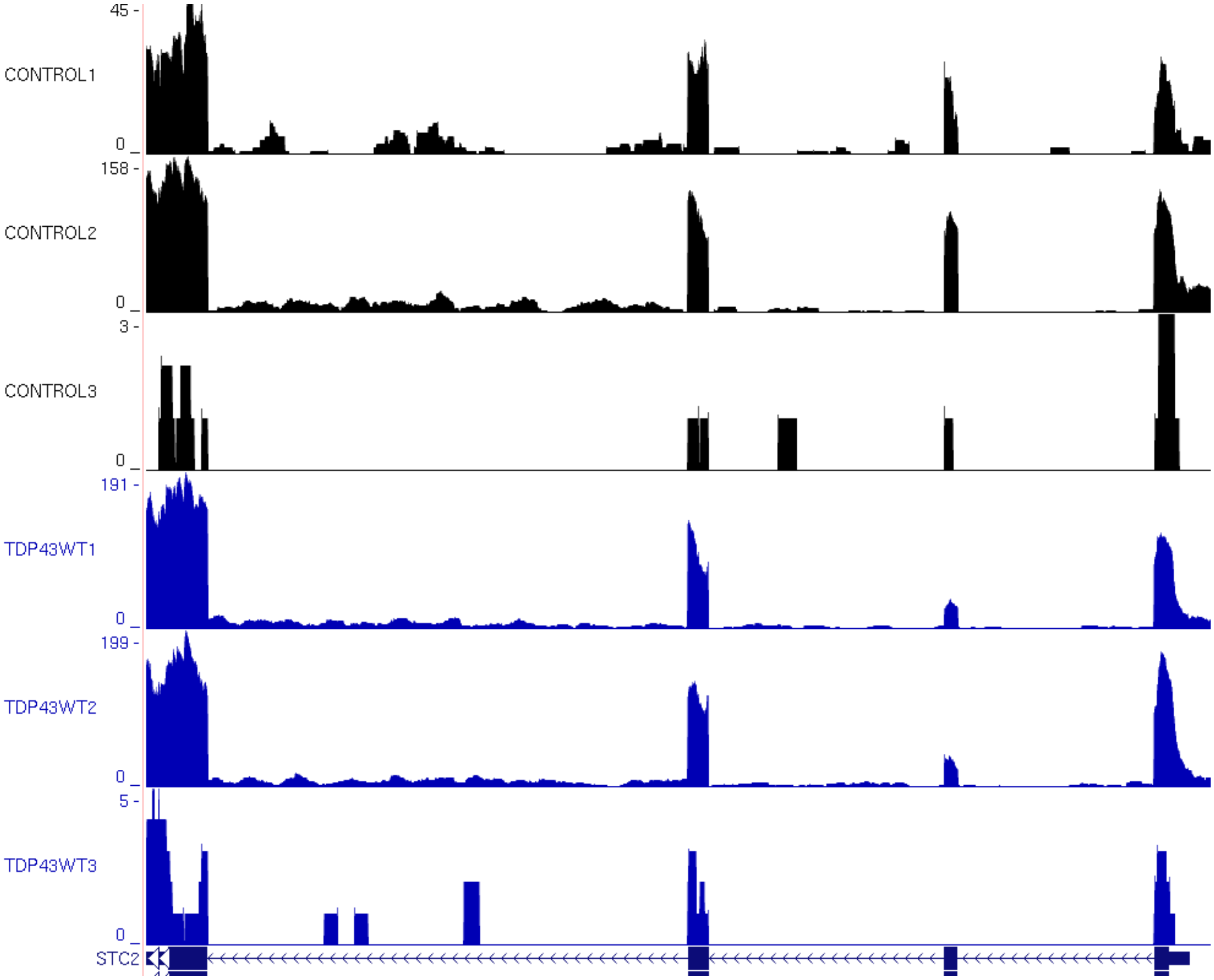
45 -
CONTROL1
0
158 -
CONTROL2
0
3 -
CONTROL3
0
191 -
TDP43WT1
0
199 -
TDP43WT2
0
5 -
TDP43WT3
0
STC2

AAD

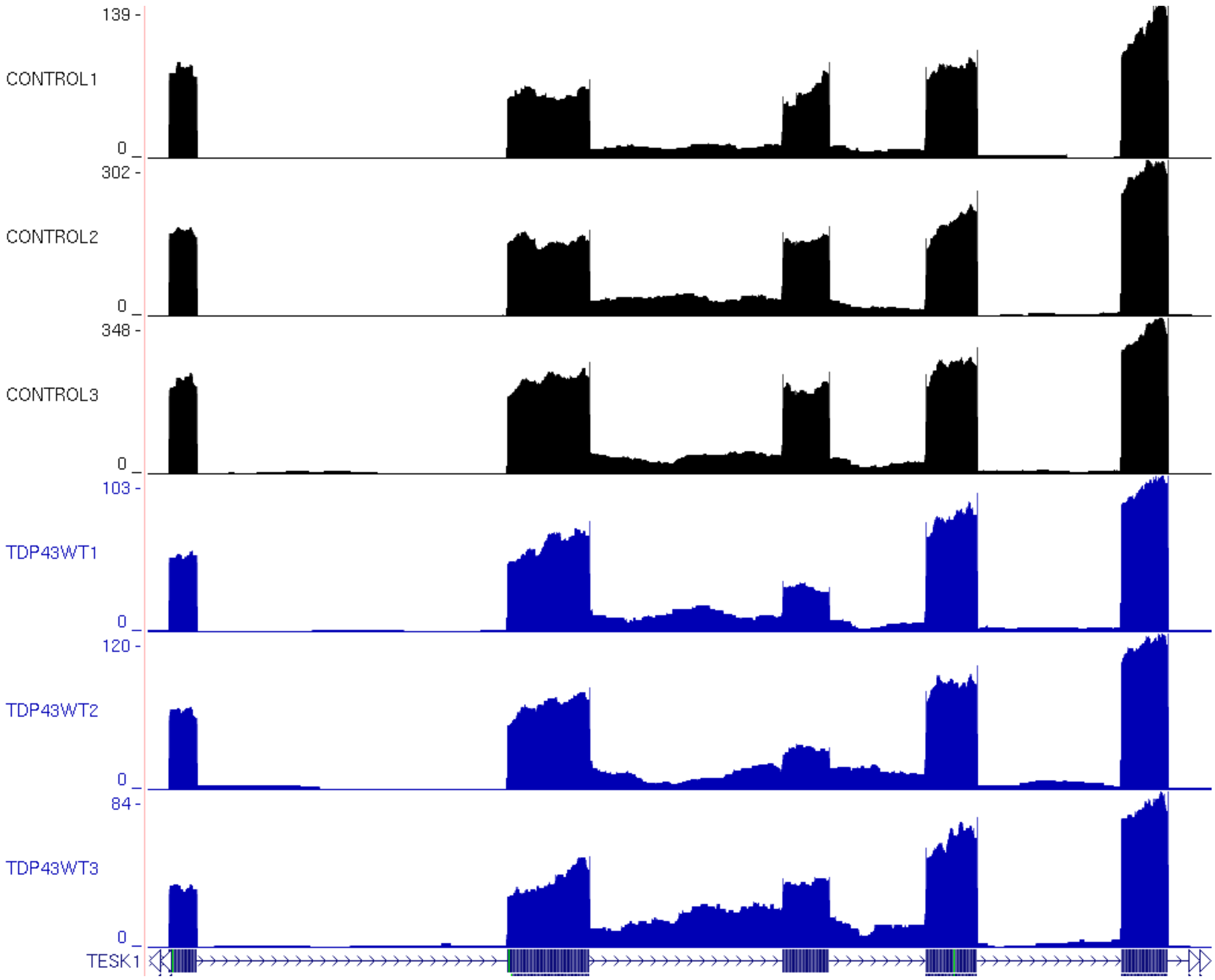
139 -
CONTROL1
0
302 -
CONTROL2
0
348 -
CONTROL3
0
103 -
TDP43WT1
0
120 -
TDP43WT2
0
84 -
TDP43WT3
0
TESK1

AAE

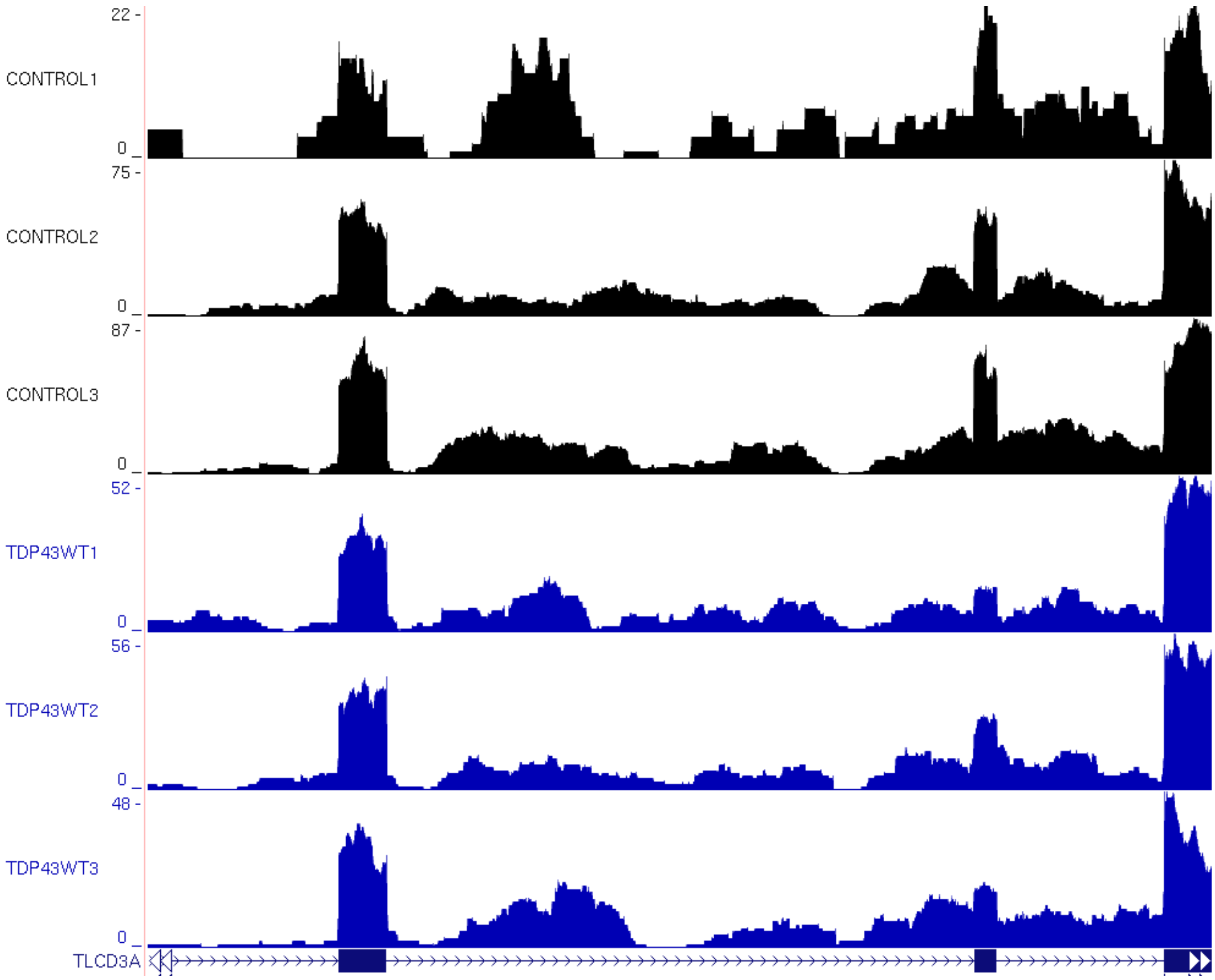
22 -
CONTROL1
0
75 -
CONTROL2
0
87 -
CONTROL3
0
52 -
TDP43WT1
0
56 -
TDP43WT2
0
48 -
TDP43WT3
0
TLCD3A

### AAF

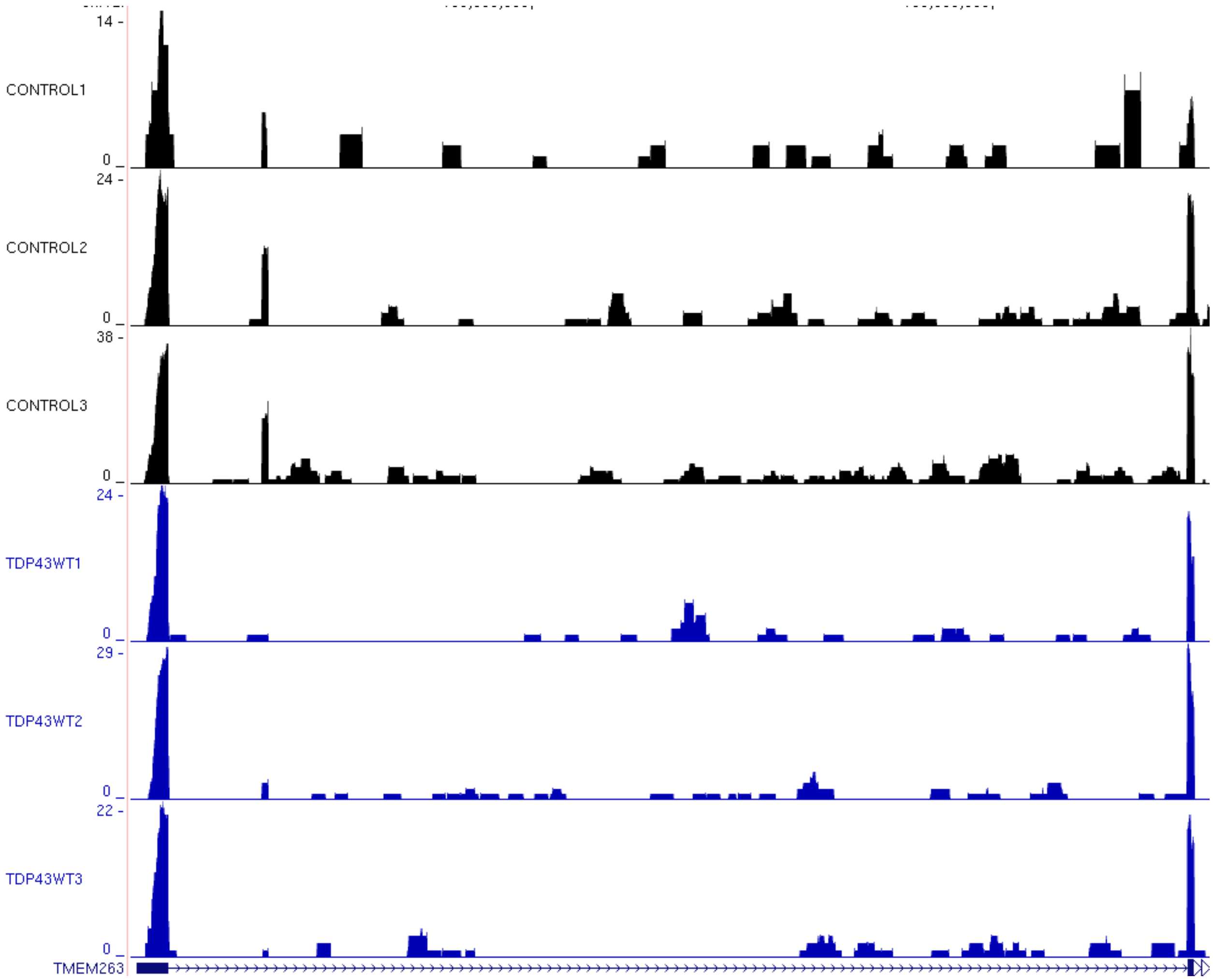
14 -
CONTROL1
0
24 -
CONTROL2
0
38 -
CONTROL3
0
24 -
TDP43WT1
0
29 -
TDP43WT2
0
22 -
TDP43WT3
0
TMEM263

### AAG

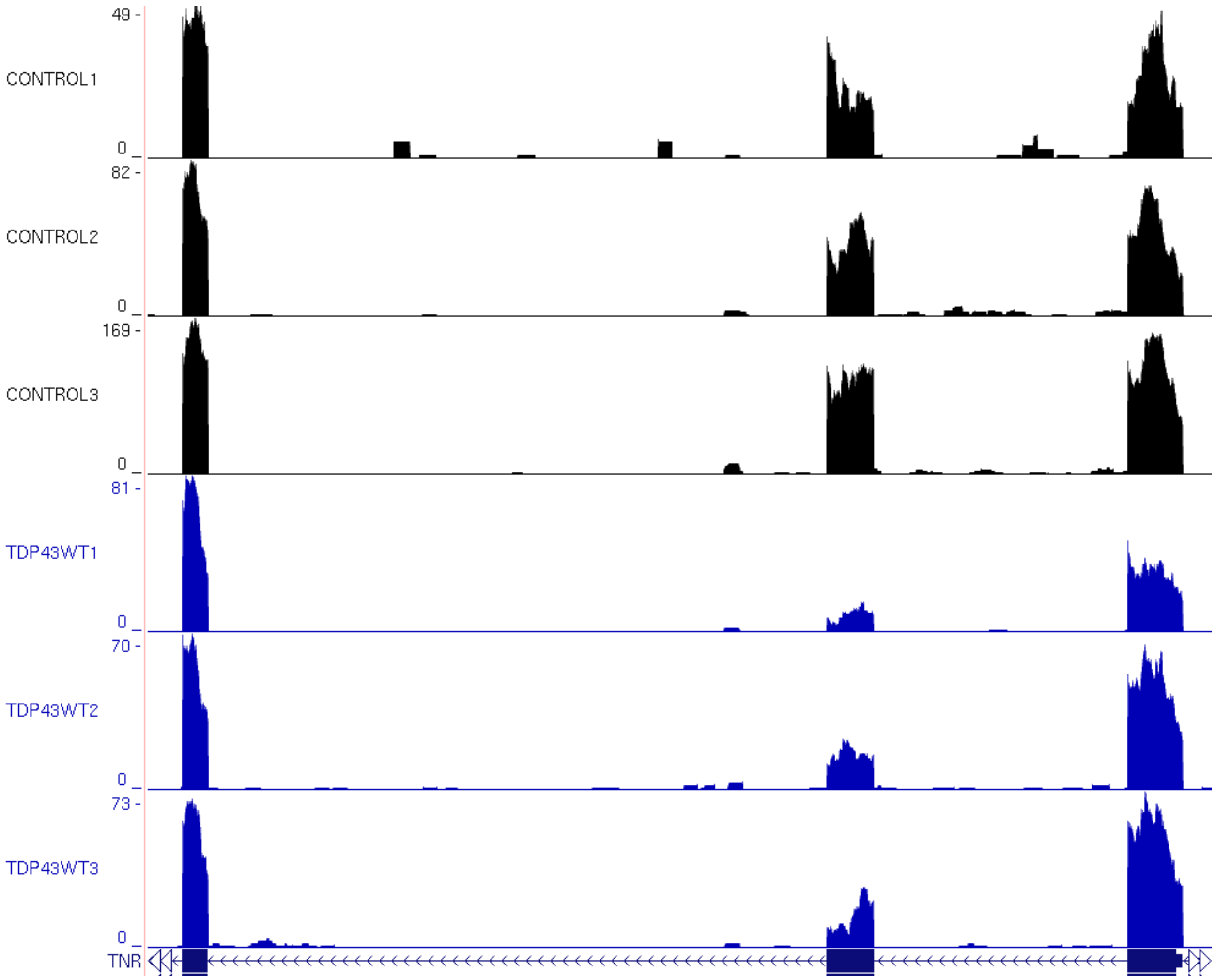

49 -
CONTROL1
0
82 -
CONTROL2
0
169 -
CONTROL3
0
81 -
TDP43WT1
0
70 -
TDP43WT2
0
73 -
TDP43WT3
0
TNR

### AAH

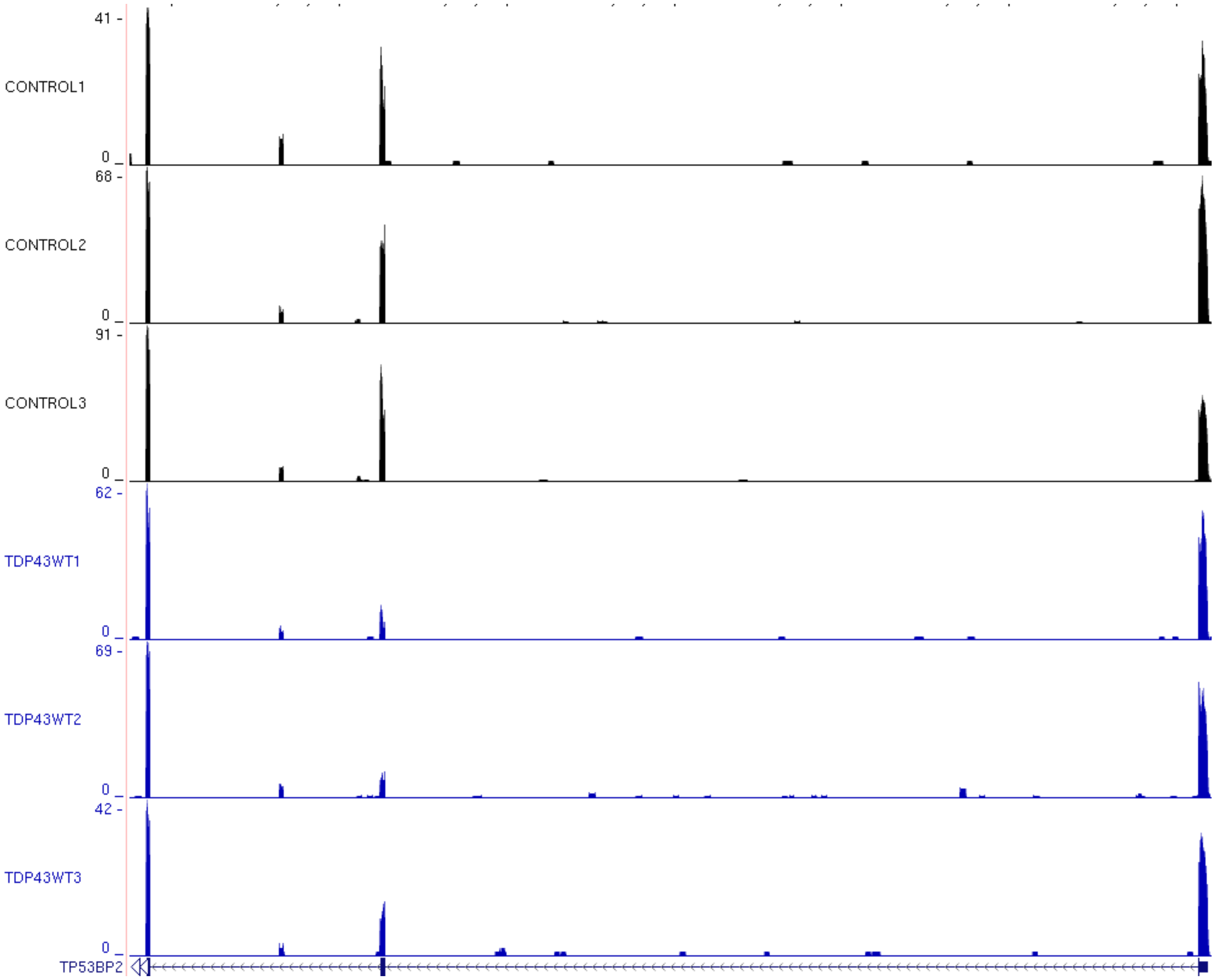

41 -
CONTROL1
0 _
68 -
CONTROL2
0 _
91 -
CONTROL3
0 _
62 -
TDP43WT1
0 _
69 -
TDP43WT2
0 _
42 -
TDP43WT3
0 _
TP53BP2

### AAI

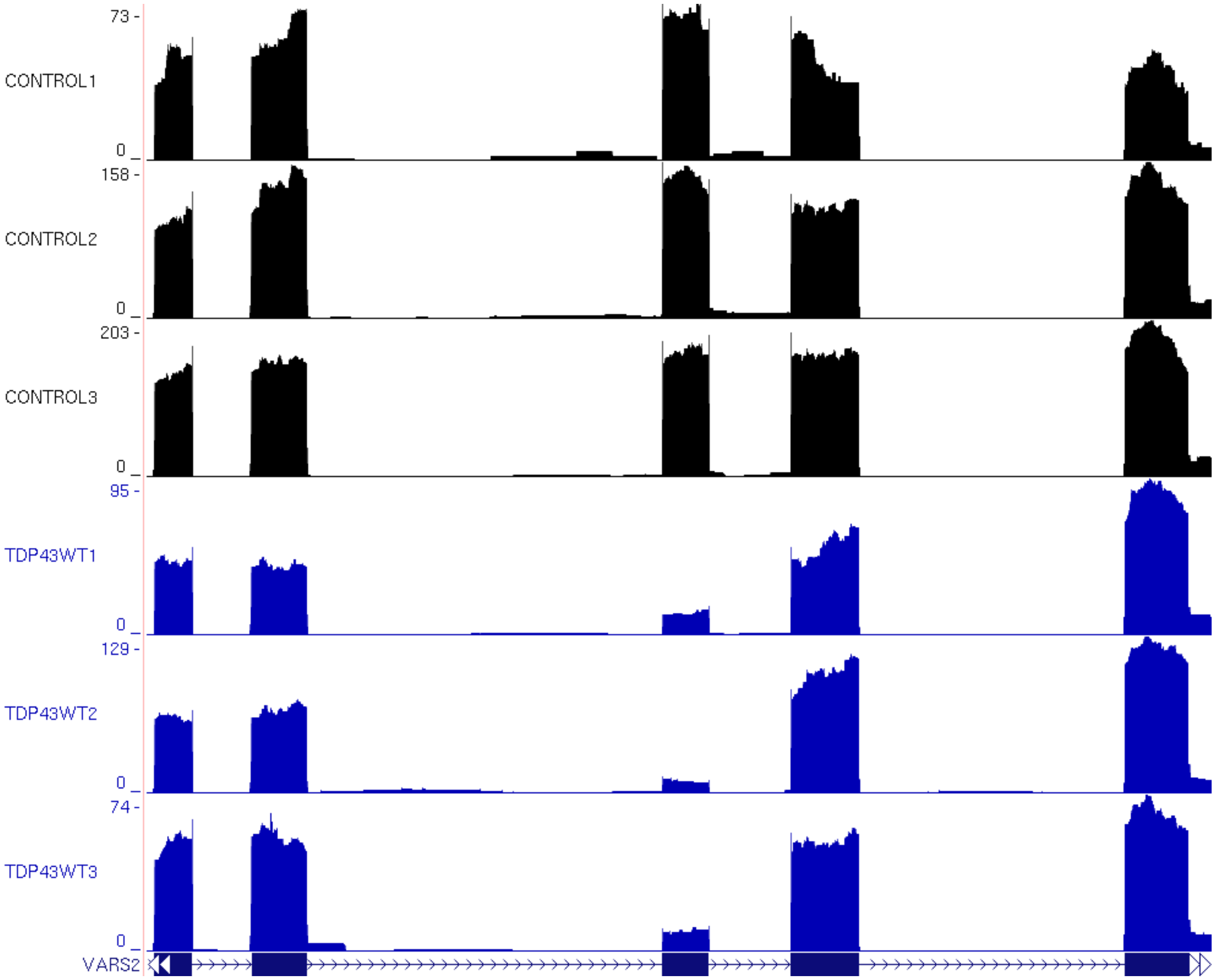
73 -
CONTROL1
0
158 -
CONTROL2
0
203 -
CONTROL3
0
95 -
TDP43WT1
0
129 -
TDP43WT2
0
74 -
TDP43WT3
0
VARS2

AAJ

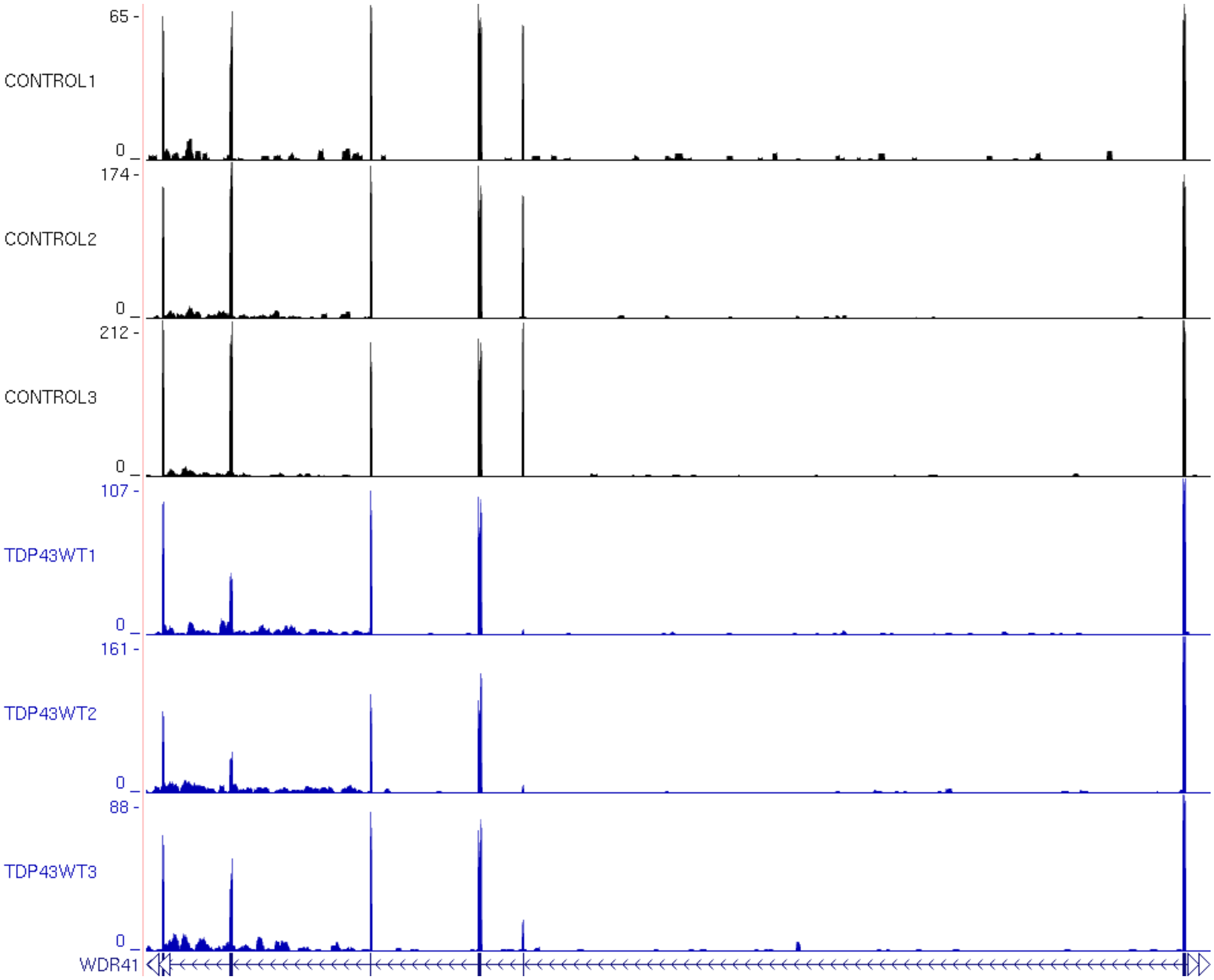

65 -
CONTROL1
0 _
174 -
CONTROL2
0 _
212 -
CONTROL3
0 _
107 -
TDP43WT1
0 _
161 -
TDP43WT2
0 _
88 -
TDP43WT3
0 _
WDR41

### AAK

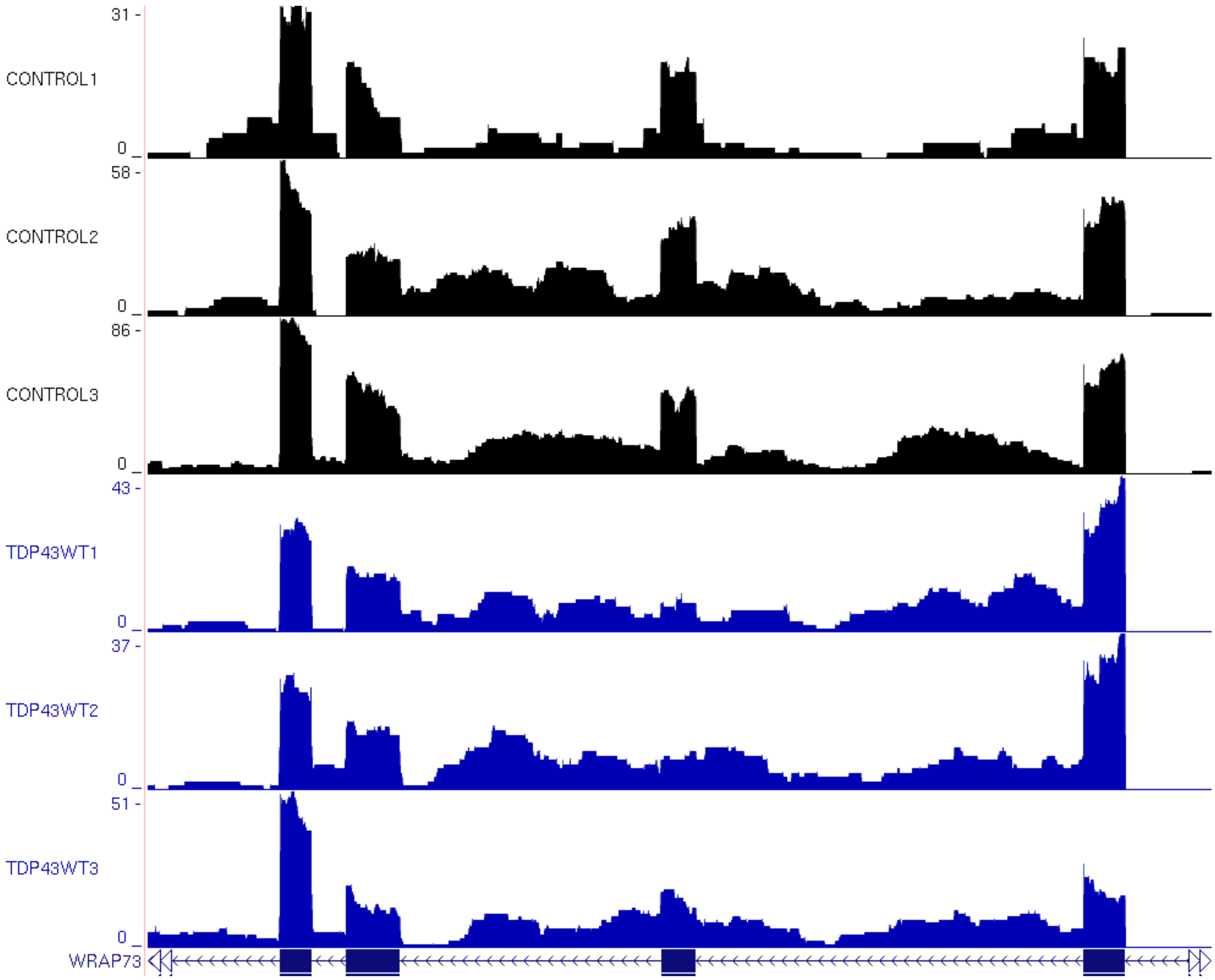

31 -
CONTROL1
0
58 -
CONTROL2
0
86 -
CONTROL3
0
43 -
TDP43WT1
0
37 -
TDP43WT2
0
51 -
TDP43WT3
0
WRAP73

### AAL

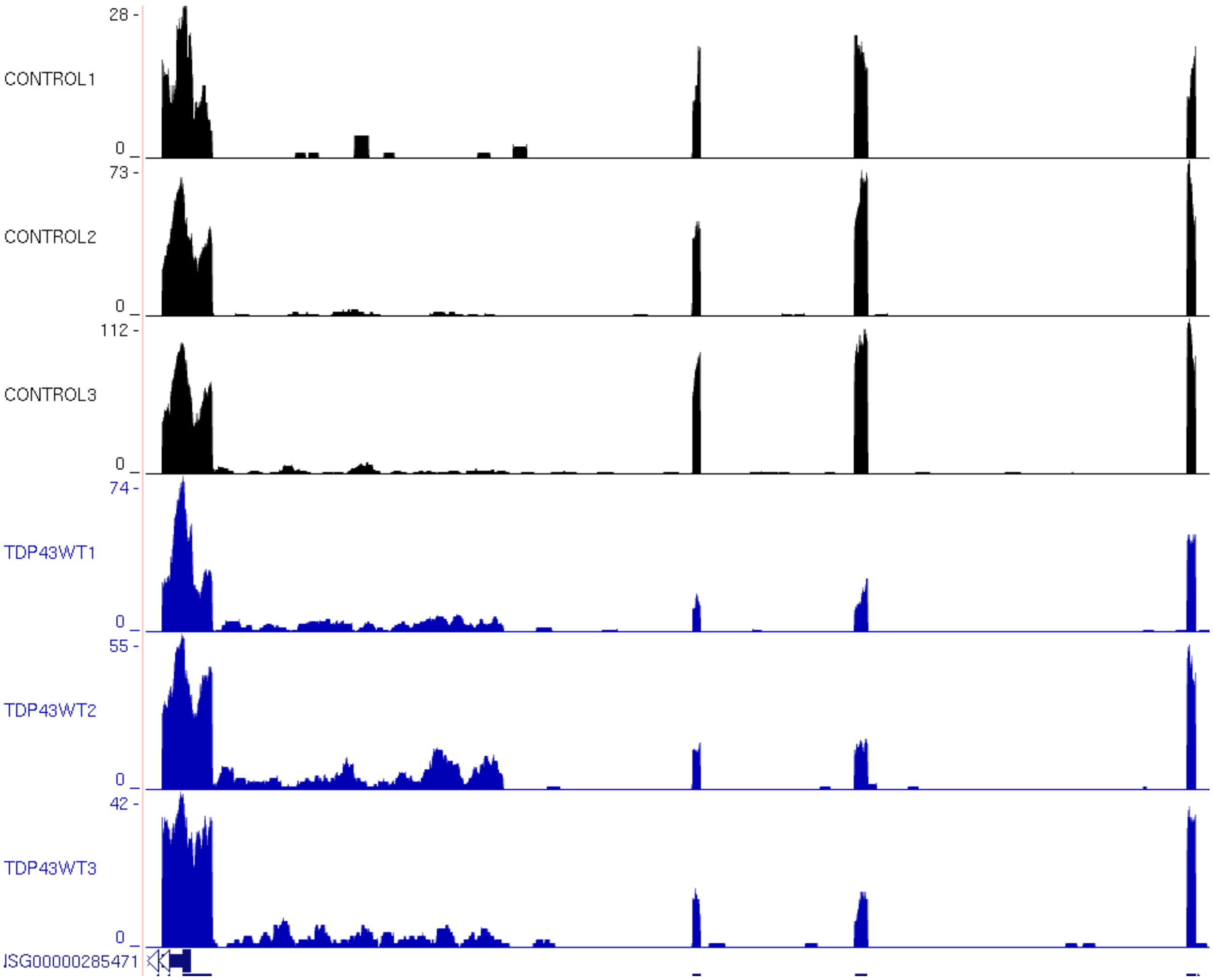

28 -
CONTROL1
0
73 -
CONTROL2
0
112 -
CONTROL3
0
74 -
TDP43WT1
0
55 -
TDP43WT2
0
42 -
TDP43WT3
0
JSG00000285471

AAM

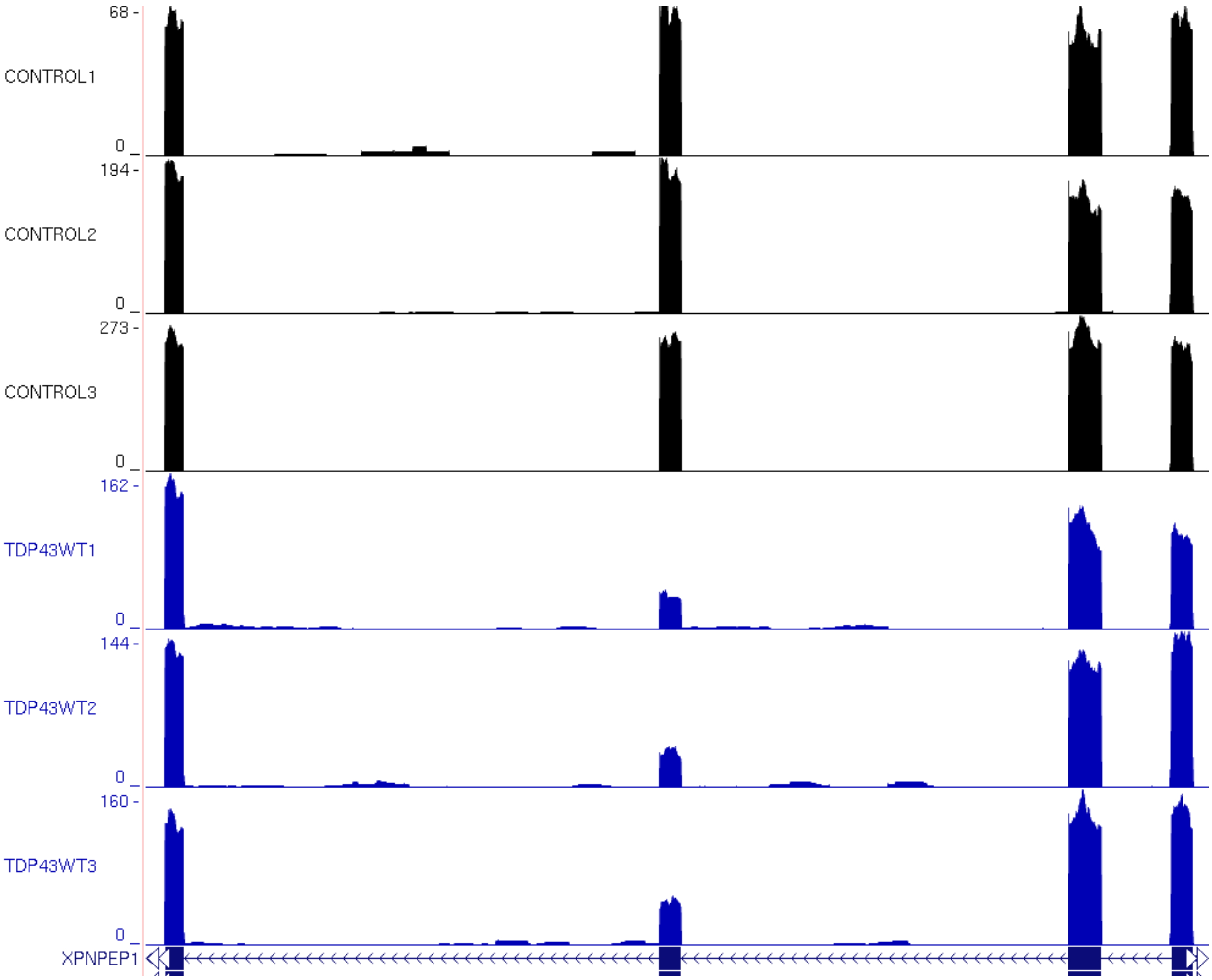
68 -
CONTROL1
0
194 -
CONTROL2
0
273 -
CONTROL3
0
162 -
TDP43WT1
0
144 -
TDP43WT2
0
160 -
TDP43WT3
0
XPNPEP1

AAN

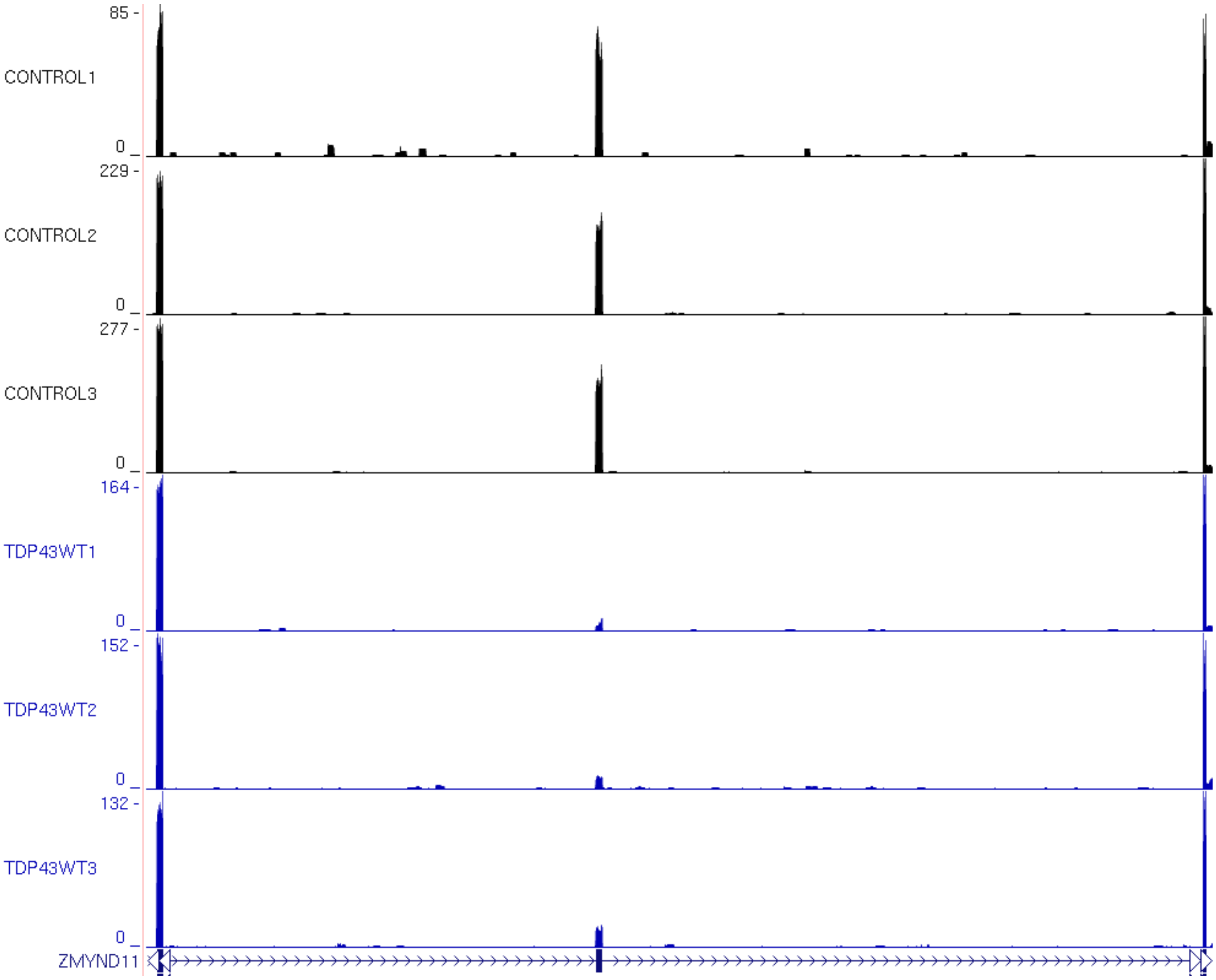

85 -
CONTROL1
0
229 -
CONTROL2
0
277 -
CONTROL3
0
164 -
TDP43WT1
0
152 -
TDP43WT2
0
132 -
TDP43WT3
0
ZMYND11

### AAO

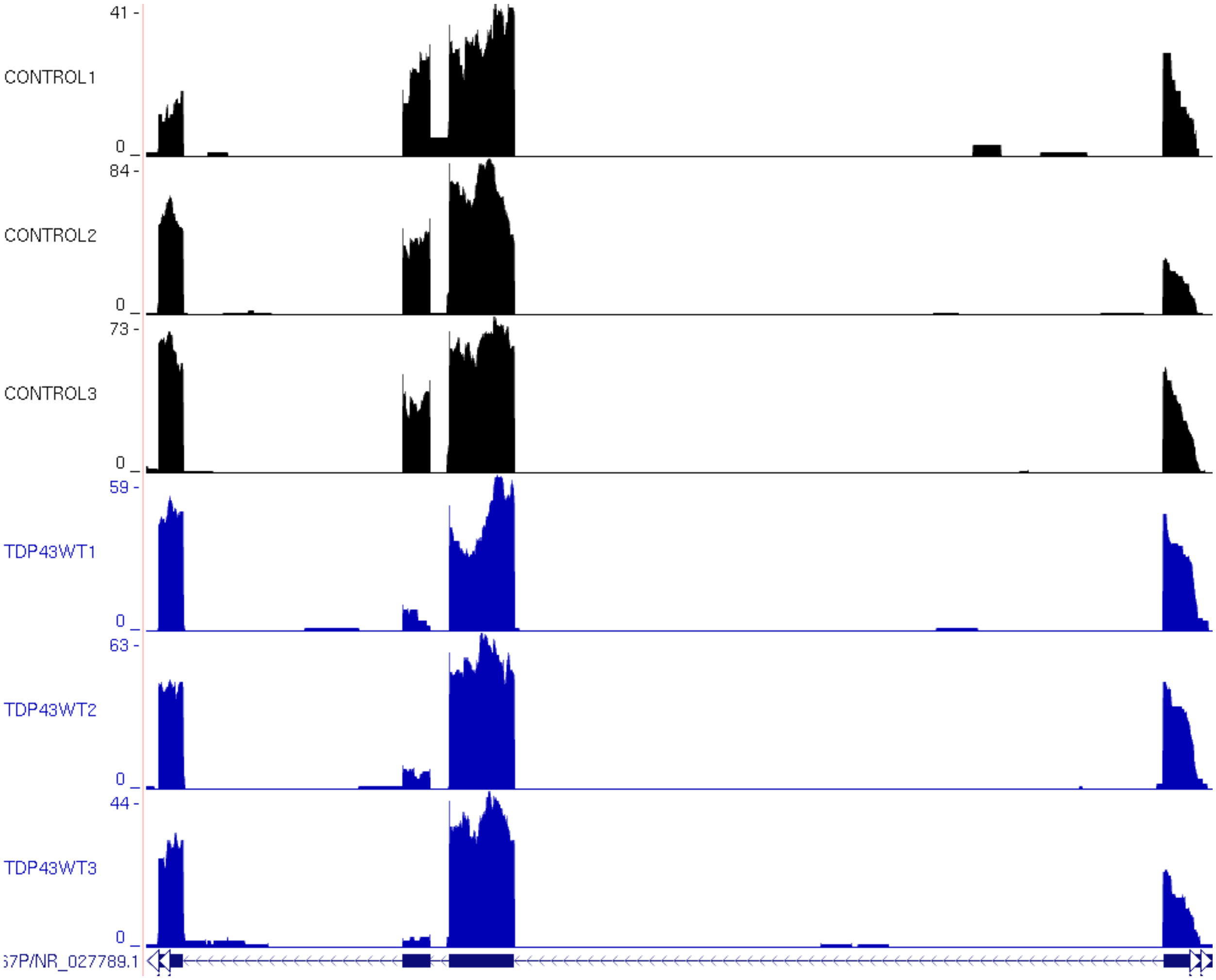

41 -
CONTROL1
0
84 -
CONTROL2
0
73 -
CONTROL3
0
59 -
TDP43WT1
0
63 -
TDP43WT2
0
44 -
TDP43WT3
0
57P/NR_027789.1

### AAP

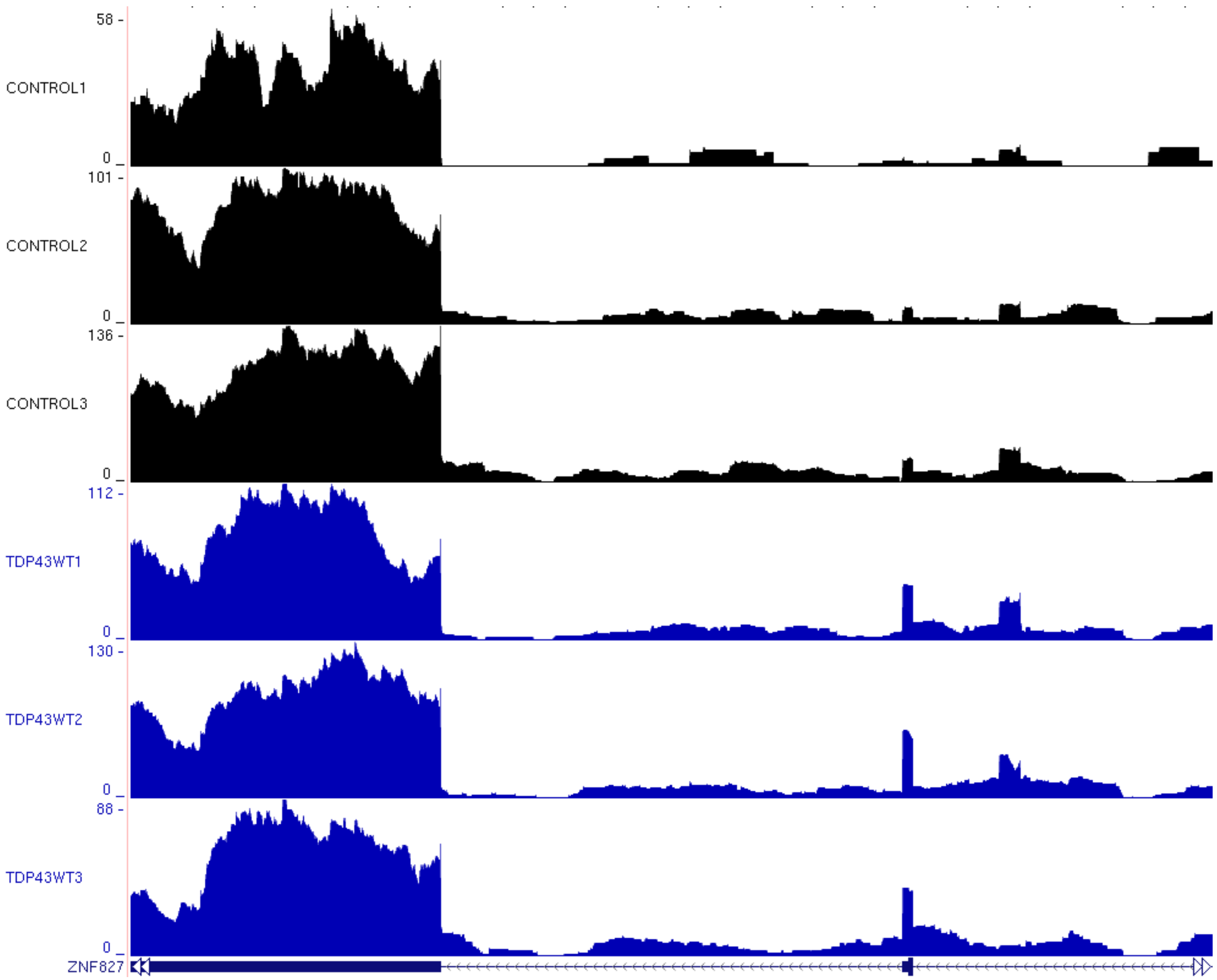
58 -
CONTROL1
0 _
101 -
CONTROL2
0 _
136 -
CONTROL3
0 _
112 -
TDP43WT1
0 _
130 -
TDP43WT2
0 _
88 -
TDP43WT3
0 _
ZNF827

**Supplementary file 2.** Examples of exon skipping events taken from UCSC genome browser from i3N human neurons infected with lentivirus overexpressing TDP-$43^{WT}$, (TDPWT1, TDPWT2, TDPWT3) and i3N non infected cells (CONTROL1, CONTROL2, CONTROL3). A = ADAM23, B = AMT, C = BMPR1A, D = BRI3BP, E = CANX, F = CCDC126, G = CERT1, H = CLASP2, I = COQ5, J = CSDE1, K = CSPP1, L = DDL2, M = DRGX (cryptic exon), N = ELP1, O = ELP2, P = ERMAD, Q = FAM66C, R = FAM102A, S = FAT1, T = FBRS, U = FBXO22, V = FRAS1, W = GART, X = HARS2, Y = HCFC2, Z = HIRA, AA = HYOU1, AB = INTS10, AC = ITGB1, AD = KCNMA1, AE = LGI4 (cryptic exon), AF = LINC00680-GUSBP4, AG = MYBBP1A, AH = NHLRC3, AI = NICN1, AJ = NIFK, AK = NIPAL3, AL = NRCAM, AM = NUP88, AN = NUP93, AO = PLXNB1, AP = PPA1, AQ = PTS, AR = RABGGTB, AS = RCHY1, AT = RHOT2, AU = RNF114, AV = SBF1, AW = SCN3A, AX = SCN9A, AY = SESN3, AZ = SLC17A5, AAA = SLC35A5, AAB = SRPK2, AAC = STC2, AAD = TESK1, AAE = TLCD3A, AAF = TMEM263, AAG = TNR, AAH = TP53BP2, AAI = VARS2, AAJ = WDR41, AAK = WRAP73, AAL = WSCD1, AAM = XPNPEP1, AAN = ZMYND11, AAO = ZNF767P, AAP = ZNF827.
